# Reduced sensitivity of infectious SARS-CoV-2 variant B.1.617.2 to monoclonal antibodies and sera from convalescent and vaccinated individuals

**DOI:** 10.1101/2021.05.26.445838

**Authors:** Delphine Planas, David Veyer, Artem Baidaliuk, Isabelle Staropoli, Florence Guivel-Benhassine, Maaran Michael Rajah, Cyril Planchais, Françoise Porrot, Nicolas Robillard, Julien Puech, Matthieu Prot, Floriane Gallais, Pierre Gantner, Aurélie Velay, Julien Le Guen, Najibi Kassis-Chikhani, Dhiaeddine Edriss, Laurent Belec, Aymeric Seve, Hélène Péré, Laura Courtellemont, Laurent Hocqueloux, Samira Fafi-Kremer, Thierry Prazuck, Hugo Mouquet, Timothée Bruel, Etienne Simon-Lorière, Felix A. Rey, Olivier Schwartz

**Affiliations:** Virus & Immunity Unit, Department of Virology, Institut Pasteur; CNRS UMR 3569, Paris, France; Vaccine Research Institute, Creteil, France; INSERM, Functional Genomics of Solid Tumors (FunGeST), Centre de Recherche des Cordeliers, Université de Paris and Sorbonne Université, Paris, France; Hôpital Européen Georges Pompidou, Laboratoire de Virologie, Service de Microbiologie, Paris, France; G5 Evolutionary genomics of RNA viruses, Department of Virology, Institut Pasteur, Paris, France; Université de Paris, Sorbonne Paris Cité, Paris, France; Laboratory of Humoral Immunology, Department of Immunology, Institut Pasteur, INSERM U1222, Paris, France; CHU de Strasbourg, Laboratoire de Virologie, Strasbourg, France; Université de Strasbourg, INSERM, IRM UMR_S 1109, Strasbourg, France; Hôpital Européen Georges Pompidou, Service de Gériatrie, Assistance Publique des Hôpitaux de Paris, Paris, France; Hôpital européen Georges Pompidou, Unité d’Hygiène Hospitalière, Service de Microbiologie, Assistance Publique-Hôpitaux de Paris, Paris, 75015, France; CHR d’Orléans, service de maladies infectieuses, Orléans, France; Structural Virology Unit, Department of Virology, Institut Pasteur; CNRS UMR 3569, Paris, France

## Abstract

The SARS-CoV-2 B.1.617 lineage emerged in October 2020 in India^1–6^. It has since then become dominant in some indian regions and further spread to many countries. The lineage includes three main subtypes (B1.617.1, B.1617.2 and B.1.617.3), which harbour diverse Spike mutations in the N-terminal domain (NTD) and the receptor binding domain (RBD) which may increase their immune evasion potential. B.1.617.2 is believed to spread faster than the other versions. Here, we isolated infectious B.1.617.2 from a traveller returning from India. We examined its sensitivity to monoclonal antibodies (mAbs) and to antibodies present in sera from COVID-19 convalescent individuals or vaccine recipients, in comparison to other viral lineages. B.1.617.2 was resistant to neutralization by some anti-NTD and anti-RBD mAbs, including Bamlanivimab, which were impaired in binding to the B.1.617.2 Spike. Sera from convalescent patients collected up to 12 months post symptoms and from Pfizer Comirnaty vaccine recipients were 3 to 6 fold less potent against B.1.617.2, relative to B.1.1.7. Sera from individuals having received one dose of AstraZeneca Vaxzevria barely inhibited B.1.617.2. Thus, B.1.617.2 spread is associated with an escape to antibodies targeting non-RBD and RBD Spike epitopes.

## Introduction

SARS-CoV-2 variants of concern (VOCs) with mutations increasing inter-individual transmission or allowing immune evasion have supplanted ancestral strains ^7–10^. While the mutations have arisen in different viral proteins, of particular interest is the Spike protein, which promotes viral binding to the ACE2 receptor, mediates viral fusion and entry, and is targeted by neutralizing antibodies ^10,11^. The first identified variant included the D614G mutation which enhanced viral infectivity ^7,10,12^. VOC then appeared in multiple countries. The B.1.1.7 variant was first identified in the UK, B.1.351 in South Africa and the P.1 lineage in Brazil ^7–10,13–15^. They harbour mutations that enhance Spike affinity to ACE2, such as N501Y, which is present in the three variants and K417N/T found in B.1.351 and P.1 ^10,16–21^. Antibody escape mutations in the RBD include E484K found in B.1.351 and P.1 ^10,16–21^. Another variant termed B.1.247/B.1.429 was originally detected in California. It lacks the E484K substitution and carries a different set of mutations in the NTD (S13I, W152C) and RBD (L452R). The L452R mutation reduces or abolishes neutralization by some monoclonal antibodies (mAb) or sera from convalescent persons or vaccine recipients ^22–25^.

Neutralizing antibodies also target non-RBD Spike epitopes, particularly in the NTD ^26^. The role of the NTD during viral entry is not fully understood. It has been proposed to recognize sugar moieties upon initial binding and might be involved in pre-fusion to post-fusion transition of the Spike ^27^. Some mAbs targeting the NTD may exhibit high neutralization potency without preventing Spike binding to ACE2^26,27^. These non-RBD antibodies are highly prevalent and may represent more than 80% of the anti-Spike IgG response, even if 90% of the neutralizing activity targets the RBD ^17,27^. VOC contain changes in the NTD that have been associated with antibody escape ^10,26,28^. These NTD modifications include the 69-70 and 144-145 deletions in B.1.1.7, the 241-243 deletion and R246I in B.1.351, W152C in B.1.429, and L18F substitution in a small proportion of B.1.351 and P.1 ^10,26,28^.

India is facing a large surge in COVID-19 cases since early 2021. Many cases are caused by a new lineage, termed B.1.617 ^1–6^. The epidemiology data are scarce, but B.1.617 has become the predominant strain in the state of Maharashtra and probably in other Indian regions ^4^. It has also been detected in many countries including UK. This lineage is under evolution and has been subdivided into three main sublineages, B.1.617.1, B.1.617.2 and B.1.617.3 ^29^ The B.1.617.2 sublineage has been recently classified as a VOC, since it is believed to be more transmissible than the two other sublineages. Recent pre-prints reported a reduced sensitivity of B.1.617 to certain monoclonal and polyclonal antibodies ^1–6,30^. Experiments were so far mostly performed with single-cycle pseudotyped viral particles, and very little is known about the sensitivity of replicative B.1.617.2 to the humoral immune response.

Here, we assessed the sensitivity of an authentic B.1.617.2 isolate to various mAbs targeting the NTD and the RBD of the Spike, including 4 clinically approved therapeutic antibodies. We also examined the neutralization potency of sera from convalescent individuals and vaccine recipients, demonstrating that the variant has indeed acquired a degree of resistance to them.

## Results

### Isolation and characterization of the SARS-CoV-2 B.1.617.2 variant

We isolated the B.1.617.2 variant from a nasopharyngeal swab of a symptomatic individual, a few days upon his return to France from India. The virus was amplified by two passages on Vero E6 cells. Sequences of the swab and the outgrown virus were identical and identified the B.1.617.2 variant (GISAID accession ID: EPI_ISL_2029113) (Fig. 1a). In particular, the Spike protein contained 9 mutations, when compared to the D614G strain (belonging to the basal B.1 lineage) used here as a reference, including five mutations in the NTD (T19R, G142D, Δ156, Δ157, R158G), two in the RBD (L452R, T478K), one mutation close to the furin cleavage site (P681R) and one in the S2 region (D950N) (Fig. 1a). Viral stocks were titrated using S-Fuse reporter cells and Vero cells ^21,31^. S-Fuse cells are convenient for rapid viral titration and measurement of neutralizing antibodies. They generate a GFP signal as soon as 6 hours post infection and the number of GFP+ cells correlates with the viral inoculum ^31, 21^. Viral titers were similar in the two target cells and reached 10^5^ to 10^6^ infectious units/ml. Large syncytia were observed in B.1.617.2-infected Vero and S-Fuse cells (Fig. S1). As expected, the syncytia were positive for Spike staining (Fig. S1). Future work will help determining whether B.1.617.2 is more fusogenic than D614G or other variants, as suggested here by the large size of B.1.617.2-induced syncytia.

**Fig. 1.**
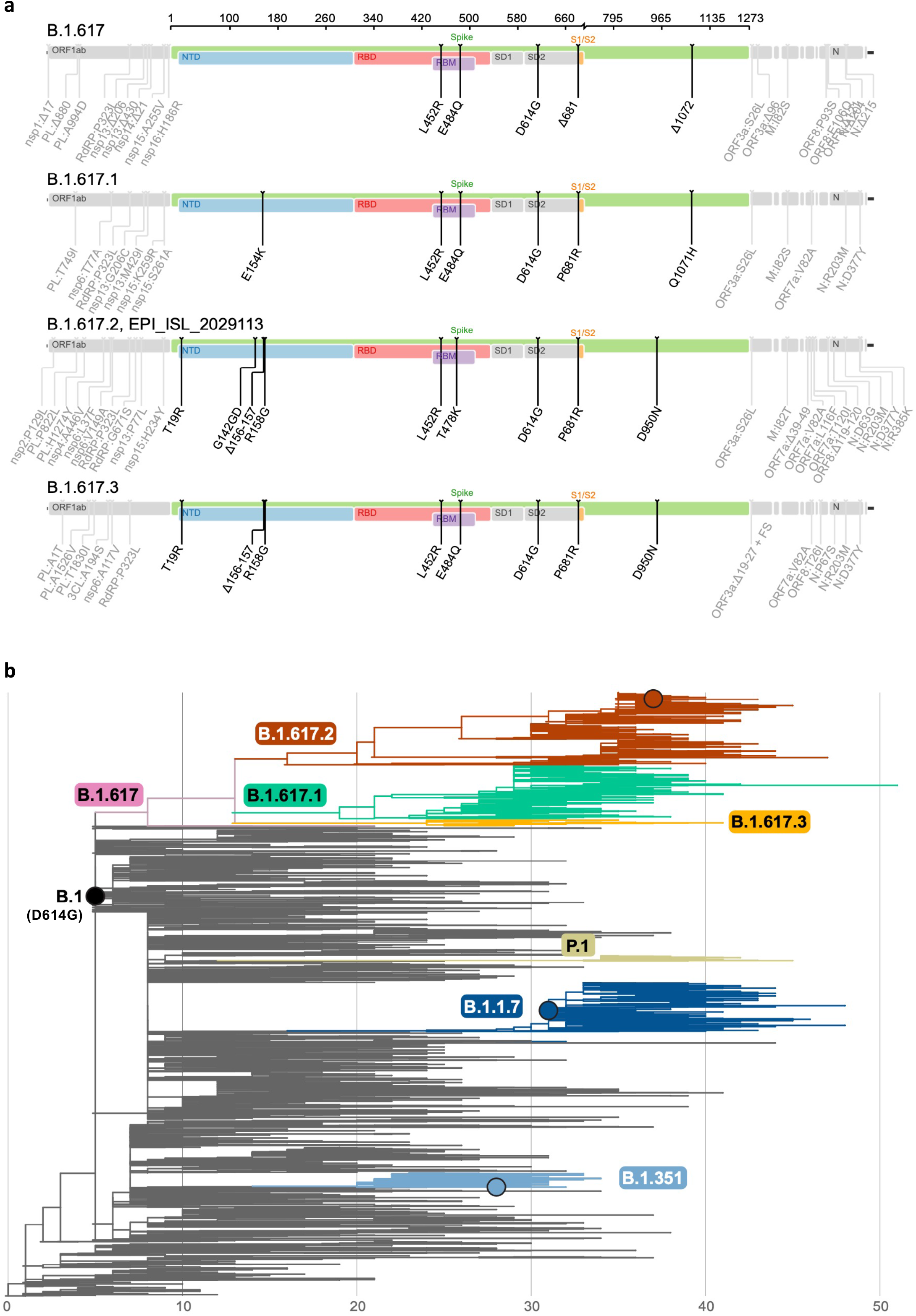
Phylogenetic analysis of the B.1.617 lineage. **a.** Schematic overview of B.1.617 sublineage consensus sequences with a focus on the Spike built with the Sierra tool ^58^. Amino acid modifications in comparison to the ancestral Wuhan-Hu-1 sequence (NC_045512) are indicated. **b.** Global phylogeny of SARS-CoV-2 highlighting the B.1.617 lineage. The maximum likelihood tree was inferred using IQ-Tree, as implemented in the Nextstrain pipeline on a subsampled dataset of 3794 complete genomes. Branch lengths are scaled according to the number of nucleotide substitutions from the root of the tree. The branches corresponding to key lineages are colored: B.1.1.7, dark blue; B.1.351, light blue; P.1, beige; B.1.617, pink; B.1.617.1, green; B.1.617.2, red; B.1.617.3, orange. A black circle indicate the position of the viruses characterized here.

### Phylogenetic analysis of the B.1.617 lineage

To contextualize the B.1.617.2 isolate reported here, we inferred a global phylogeny subsampling the diversity of SARS-CoV-2 sequences available on the GISAID EpiCoV database (Fig. 1). The B.1.617 lineage, subdivided into three sublineages according to the PANGO classification ^32^, derives from the B.1 lineage (D614G). All three sublineages present multiple changes in the Spike, including the L452R substitution in the RBD, already seen in B.1.429 and the P681R, located in the furin cleavage site and which may enhance the fusogenic activity of the Spike. The E484Q, which may be functionally similar to the antibody-escape mutation E484K found in B.1.351 and P.1, is present in both B.1.617.1 and B.1.617.3, and appears to have reverted in the rapidly growing B.1.617.2 sublineage, as it was present in a sequence (B.1.617) ancestral to the three sublineages (Fig. 1a,b) ^29^. Whether the absence of the E484Q mutation, the presence of the T478K mutation or other changes in the Spike or the rest of the genome may facilitate viral replication and transmissibility remains unknown. Interestingly, the B.1.617 lineage is not homogeneous, with multiple mutations fixed in a sublineage (e.g. Spike:T19R, G142D or D950N) that are also detected at lower frequencies in the other sublineages. This may reflect founder effects or similar selective pressures acting on these emerging variants.

### Mutational changes in B.1.617.2

Upon inspection, the locations of the spike mutations in the B.1.617.2 variant showed a similar overall distribution to those that appeared in other VOC. In particular, in addition to the D614G mutation common to all current strains, the D950N mutation mapped to the trimer interface (Fig. 2a), suggesting a potential contribution in regulating Spike dynamics, as has been shown with the D614G mutation ^10^. As with other VOC, a number of mutations in B.1.617.2 cluster in the NTD (Fig. 2b). The 156-157 deletion and G158R mutation map to the same surface as the 144 deletion in B.1.1.7 and the 241-243 deletion in B.1.351. Similarly, the T19R maps to the surface patch that has several mutations in B.1.1.7. Importantly, all these altered residues lie in the NTD “supersite” that is the target of the most potently anti-NTD neutralizing antibodies ^28^. In the RBD, examining the distribution of all mutations appearing in VOC detected so far indicates that they all map to the periphery of the ACE2 binding surface (Fig. 2c), suggesting that the virus accumulates mutations there in order to reduce or avoid antibody binding while maintaining binding to ACE2. For instance, the L452R mutation found in B.1.617.2 impairs neutralization by antibodies ^22, 23,24, 25^ and is located at this periphery. The only mutation within the ACE2 patch is at location 501, which increases the affinity of the RBD for ACE2 and is also involved in antibody escape ^10 33 34^. The RBD mutation T478K in the RBD is unique to B.1.617.2 and falls within the epitope region of potent neutralizing mAbs categorized as “Class 1” (Fig. 2c) ^35^. This mutation at location 478 is in proximity of the well characterized E484K mutation known to facilitate antibody escape ^10^. These observations prompted us to analyze the neutralization potential of well characterized monoclonal antibodies, as well as sera from convalescents and vaccinees, against the B.1.617.2 variant.

**Fig. 2.**
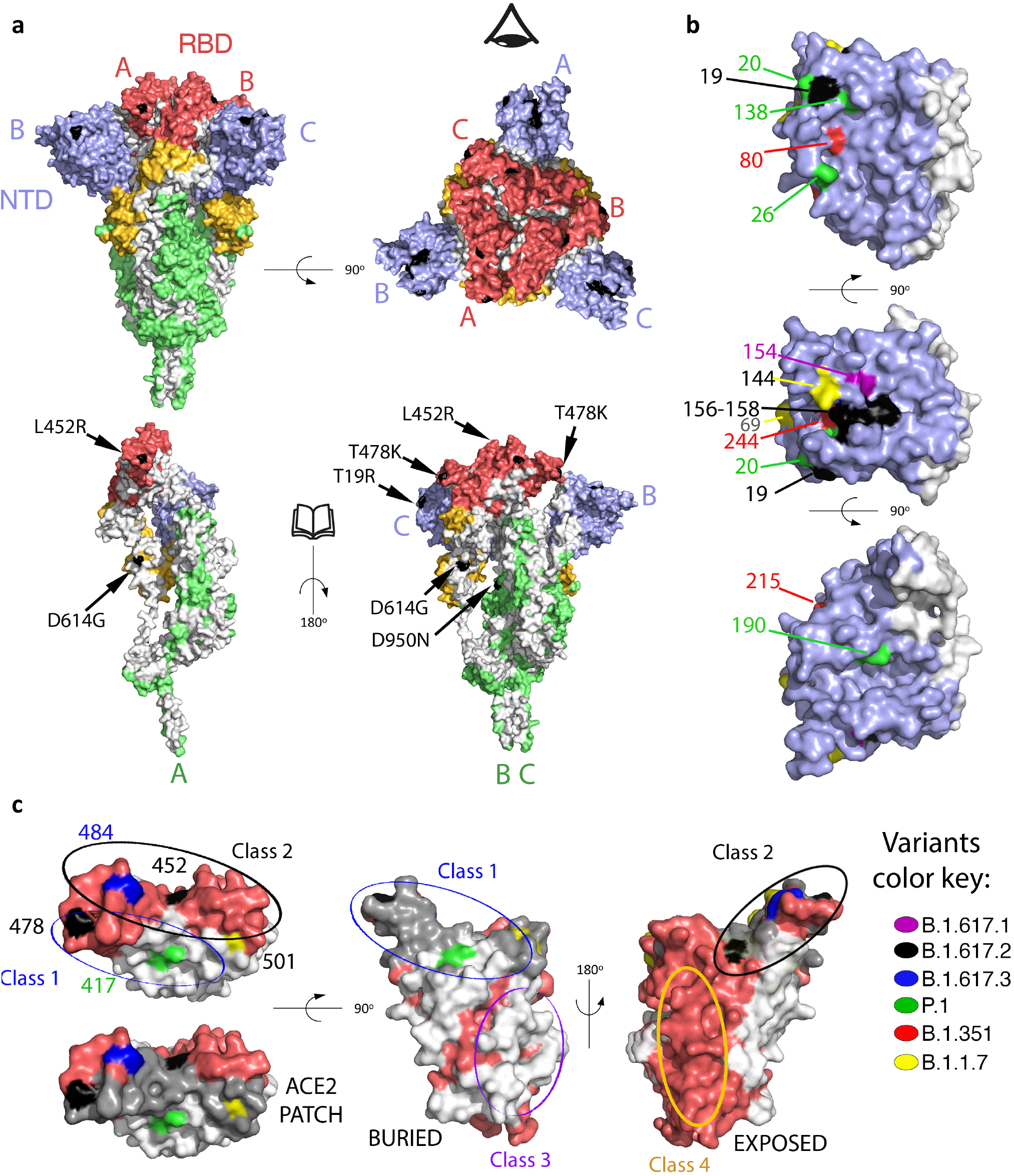
Mapping mutations on B1.617.2 and other variants of concern to the Spike surface. **a**. The Spike protein trimer (PDB:6XR8, corresponding to a closed spike trimer with all three RBDs in the “down” conformation) is shown with its surface coloured according to domains: NTD blue, RBD red, the remainder of S1 yellow and S2 green. Interfaces between protomers were left white to help visualize the protomers’ boundaries. The three polypeptide chains in the trimer were arbitrarily defined as A, B and C, labelled to identify NTD (blue) and RBD (red) in the same protomer. Surface patches corresponding to residues mutated in the B.617.2 variant are colored in black. The top panels display two orthogonal views. The bottom panels show the trimers with subunit A in the same orientation as in the panel on top, and subunits B and C rotated 180 degrees to show the trimer interface (buried regions in the trimer are left white). The eye icon on the top right panel serves to indicate the view point for the bottom right panel, after removing chain A to display internal surfaces. The mutations in B.617.2 are labelled in the bottom panel. **b.** Details of the NTD surface, with mutations that appeared in the indicated variants. **c.** RBD shown in three orthogonal views. the left panel is viewed down the surface that binds ACE2, with the surface buried in the complex with ACE2 superposed in grey in the lower panel (labelled “ACE2 patch”). The mutations on the variants are labelled. The middle panel shows the RBD surface buried in the closed spike trimer. The right panel shows its exposed surface. Note that the mutations on the RBD cluster all around the ACE2 patch. Panels were prepared with The PyMOL Molecular Graphics System, Version 2.1 Schrödinger, LLC.

### Neutralization of B.1.617.2 by monoclonal antibodies

We assessed the sensitivity of B.1.617.2 to a panel of human mAbs using the S-Fuse assay. We tested four clinically approved antibodies, Bamlanivimab (LY-CoV555), Etesevimab (LY-CoV016), Casirivimab (REGN10933) and Imdevimab (REGN10987) targeting the RBD ^20,36^ as well as four other anti-RBD (RBD-48, RBD-85, RBD-98 and RBD-109) and four anti-NTD (NTD-18, NTD-20, NTD-69 and NTD-71) antibodies that were derived from convalescent individuals (Planchais et al, in preparation). Neutralizing anti-SARS-CoV-2 mAbs targeting the RBD can be classified into 4 main categories depending on their binding epitope ^35,37^. RBD-48 and RBD-85 belong to the first category (“Class 1”) and act by blocking binding of the “up” conformation of RBD to ACE2 ^35^. The precise epitopes of RBD-98 and RBD-109 are not yet defined but overlap with those of RBD-48 and RBD-85. The anti-NTD antibodies bind uncharacterized epitopes within this domain, as assessed by Elisa (not shown).

We first measured the potency of the four therapeutic antibodies against B.1.617.2, and included as a comparison D614G (B.1), B.1.1.7 and B.1.351 strains. The four antibodies neutralized D614G, with IC50 varying from 1.2 × 10^−3^ to 6.5 × 10^−2^μg/mL (Fig. 3a). They were active against B.1.1.7, with the exception of Etesivimab, which displayed a 200-fold increase of IC50 against this variant. As previously reported, Bamlanivimab and Etesivimab did not neutralize B.1.351 ^38 39^. Bamlanivimab lost antiviral activity against B.1.617.2, in line with previous results demonstrating that L452R is an escape mutation for this mAb ^25 22^. Etesivimab, Casirivimab and Imdevimab remained active against B.1.617.2 (Fig. 3a).

The four other anti-RBD mAbs neutralized D614G. The IC50 of RBD-48 and RBD-98 were about 15-100-fold higher with B.1.1.7 than with D614G, whereas RBD-85 displayed increased activity against B.1.1.7. Three out of the four mAbs inhibited B.1.617.2, whereas RBD-85 was inactive (Fig. 3a). The four anti-NTD mAbs were globally less efficient than the anti-RBD mAbs. They all inhibited D614G, with high IC50 (1-60 μg/mL) (Fig. 3a). Three out of the four anti-NTD antibodies totally lost any activity against both B.1.1.7 and B.1.617.2, whereas the fourth (NTD-18) inhibited to some extent the two variants. Thus, B.1.617.2 escapes neutralization by some antibodies targeting either the RBD or NTD.

**Fig. 3.**
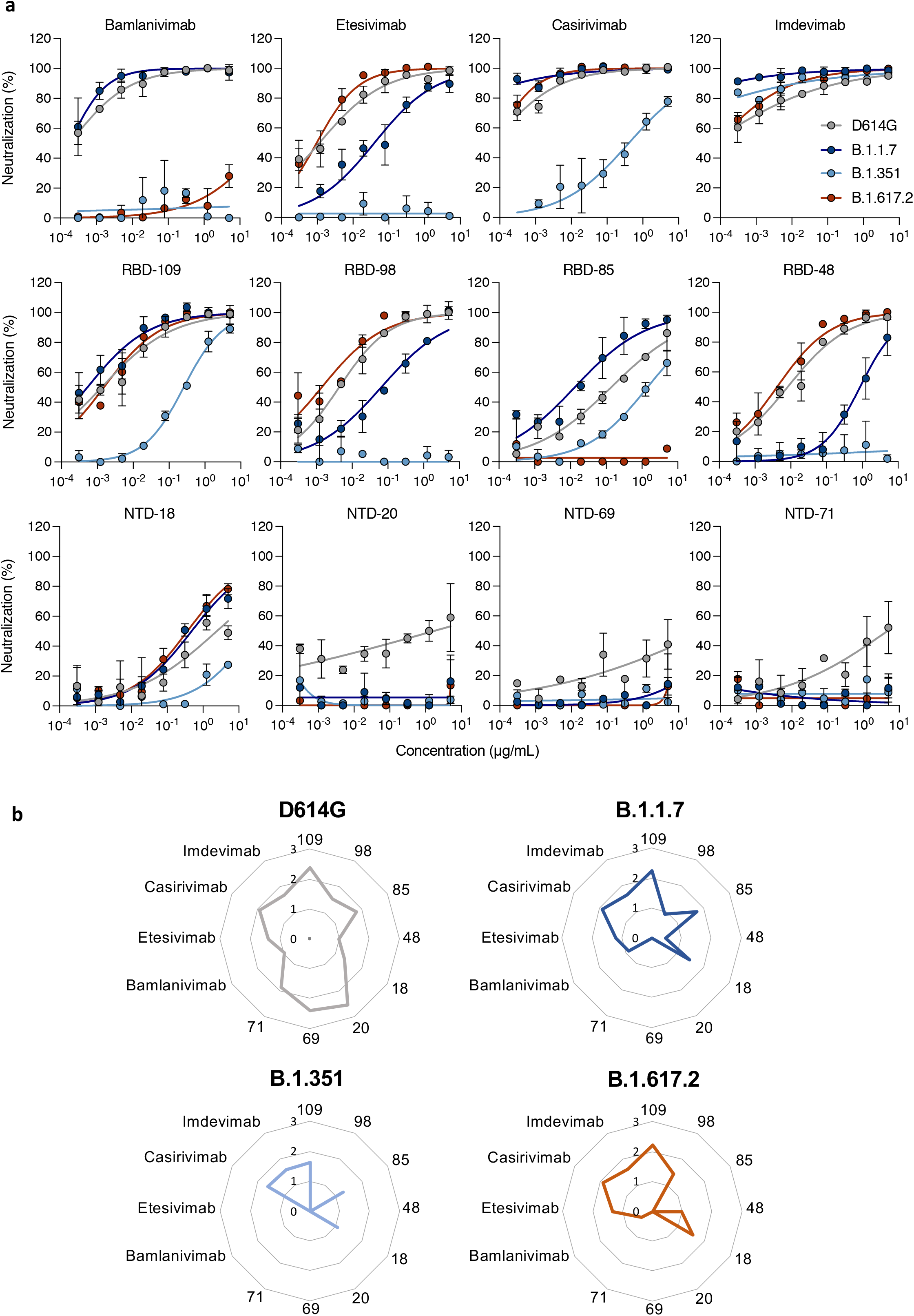
Neutralization of D614G, B.1.1.7, B.1.351 and B.1.617.2 variants by anti-SARS-CoV-2 mAbs. **a.** Neutralization curves of mAbs. Dose response analysis of the neutralization by four therapeutic mAbs (Bamlanivimab, Etesivimab, Casirivimab and Imdevimab), four anti-RBD and four anti-NTD, on the indicated SARAS-CoV-2 variants (D614G (grey), B.1.1.7 (dark blue), B.1.351 (light blue) and B.1.617.2 (orange) strains. Data are mean ± SD of three independent experiments. **b.** Binding of mAbs to infected Vero cells. Vero cells were infected with the indicated variants at a MOI of 0.1. After 48h, cells were stained with anti-SARS-CoV-2 mAbs (1μg/ml) and analyzed by flow-cytometry. Radar charts represent for each antibody the logarithm of the mean of fluorescent intensity of the staining, relative to the non-infected condition. Data are representative of three independent experiments.

We next examined by flow cytometry the binding of each antibody to Vero cells infected with the different variants. An example of binding with two therapeutic antibodies (Bamlanivimab and Imdevinab) and one anti-RBD antibody (RBD-85) is depicted in Fig. S3. Radar plots show the binding of all antibodies tested (Fig. 3b). D614G was recognized by all 12 mAbs tested. B.1.1.7 and B.1.617.2 were recognized by 9 and 7 mAb, respectively. Bamlanivimab no longer bound B.1.617.2. We also analyzed the binding of the 12 mAbs to the B.1.351 variant, which is known to be more resistant to neutralization. Only 5 of the antibodies bound this variant, whereas Bamlanivimab and Etesivimab lost their binding to B.1.351 (Fig. 3b). Altogether, these results indicate that the escape of B.1.617.2 and other variants to neutralization is due to a reduction or loss of binding of the antibodies to their target.

### Sensitivity of B.1.617.2 to sera from convalescent individuals at 6 and 12 months post-infection

We examined the neutralization ability of sera from convalescent subjects. We first selected samples from 56 donors in a cohort of infected individuals from the French city of Orléans. All individuals were diagnosed with SARS-CoV-2 infection by RT-qPCR or serology and included critical, severe, and mild-to-moderate COVID-19 cases (Table S1). They were not vaccinated at the sampling time. We recently characterized the potency of these sera against D614G, B.1.1.7 or B.1.351 ^21^. We analyzed individuals sampled at a median of 188 days (range 114-205), referred to as Month 6 (M6) samples. We incubated serially diluted sera with B.1.1.7 or B.1.617.2 strains, added the mixture to S-Fuse cells, and scored GFP+ cells after overnight infection. We then calculated ED50 (Effective Dose 50%) for each combination of serum and virus (Fig. 4a). With B.1.1.7, we obtained similar ED50 values in this series of experiments (median of 811) than in our previous analysis (median of 761) ^21^. This allowed us to compare the variants together across different experiments. With B.1.617.2, the neutralization titers were significantly decreased by 4 to 6-fold when compared to D614G and B.1.1.7 strains, respectively (Fig. 4a). This reduction in neutralizing titers was similar against B.1.617.2 and B.1.351 (Fig. 4a).

**Fig. 4.**
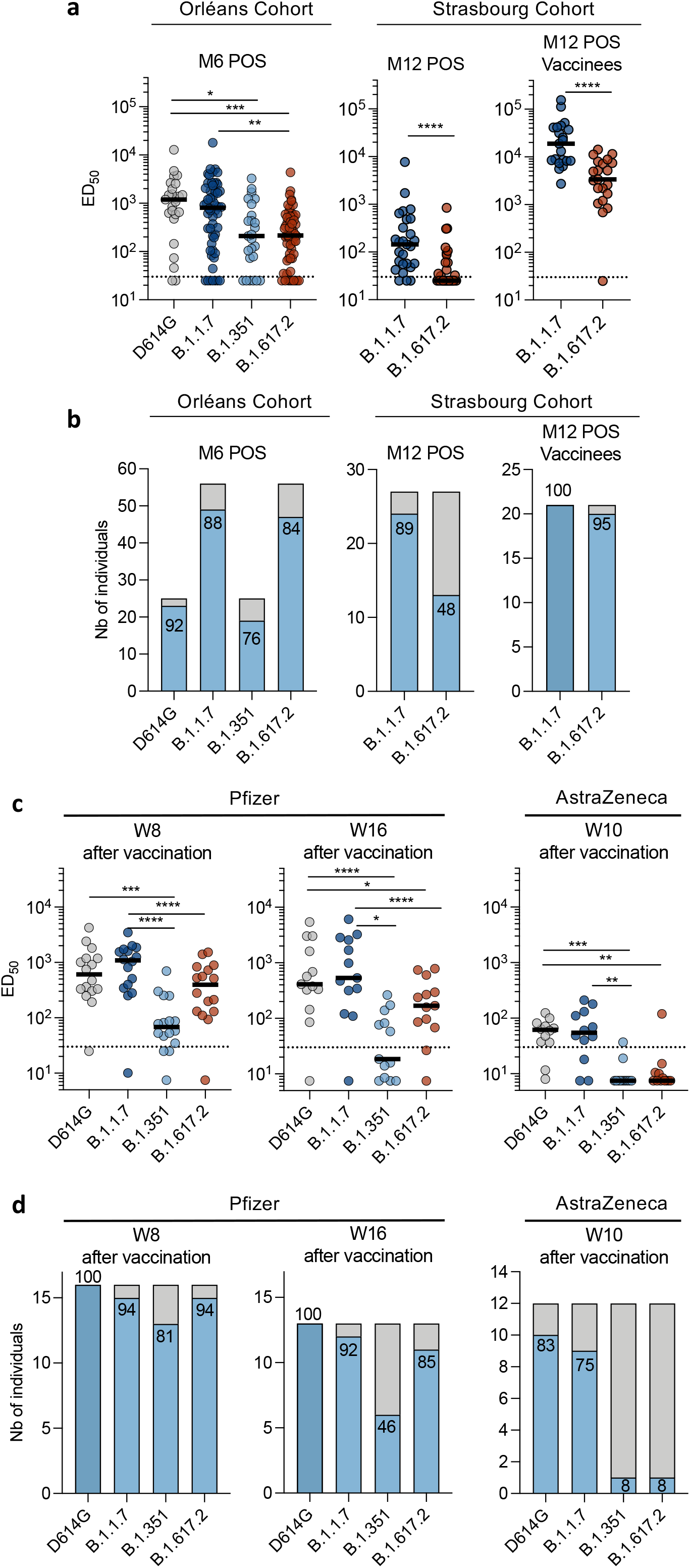
Sensitivity of SARS-CoV-2 D614G, B.1.1.7, B.1.351 and B.1.617.2 variants to sera from convalescent individuals and vaccine recipients. **a.** ED50 of neutralization of the four viral isolates. Sera from the Orleans cohort (n=25 to 56 convalescents) were sampled at 6 months post-infection (M6). Sera from Strasbourg cohort (n=26 convalescents and n=21 convalescent vaccinees) were sampled at M12. Data are mean from two independent experiments. The dotted line indicates the limit of detection (ED50=30). Two-sided Kruskal-Wallis or Friedman test with Dunn’s multiple comparison was performed between each viral strain. **b.** Each subject was arbitrarily defined as “neutralizer” (in blue) when a neutralizing activity was detected at the first (1/30) serum dilution or “non-neutralizer” (in grey) when no activity was detected. The numbers indicate the % of neutralizers. **c.** ED50 of neutralization of the four viral isolates. Sera from Pfizer vaccinated recipients were sampled at week 8 (W8) and W16 post-vaccination (W5 and W13 post second dose (n=16 and n=13, respectively). Sera from AstraZeneca vaccinated recipients were sampled at W10 post-vaccination (n=12). Data are mean from two independent experiments. The dotted line indicates an ED50 of 30. A two-sided Friedman test with Dunn’s multiple comparison was performed between each viral strain at the different time points. **d.** Each subject was arbitrarily defined as “neutralizer” (in blue) when a neutralizing activity was detected at the 1/30 serum dilution or “non-neutralizer” (in grey) when no activity was detected. The numbers indicate the % of neutralizers.

We asked whether this neutralization profile was maintained for longer periods of time. We analyzed sera from 48 individuals from another cohort of RT-qPCR-confirmed health care workers from Strasbourg University Hospitals who experienced mild disease ^40–42^. The samples were collected at a later time point (M12), with a median of 330 days (range 144-383) POS (Table S1) ^42^. We compared their potency against B.1.1.7 and B.1.617.2 strains. Twenty seven individuals were unvaccinated, whereas 21 received a single dose of vaccine 7-81 days before sampling. As previously reported, the neutralization activity against B.1.1.7 was globally low at M12 in unvaccinated individuals ^42^ (Fig. 4a). The neutralizing activity was even lower against B.1.617.2 at this time point, with a median ED50 of 25, representing a 6-fold decrease when compared to B.1.1.7 (Fig. 3a). The 13 single-dose vaccine recipients of the M12 cohort included 9 vaccinated with AstraZeneca, 9 with Pfizer and 3 with Moderna vaccines. Sera from these vaccinated participants showed a 130-fold increase in the median neutralizing antibody titer against both B.1.1.7 and B.1.617.2 (Fig. 3b). Therefore, our results extend previous results obtained with other variants, showing that a single dose of vaccine boosts cross-neutralizing antibody responses ^17,42–45^.

We then classified the cases as neutralizers (defined as harboring neutralizing antibodies detectable at the first serum dilution of 1/30) and non-neutralizers, for the different viral variants and the two cohorts (Fig. 4b). Between 76% and 92% of the individuals neutralized the four strains at M6. The fraction of neutralizers was lower in the second cohort at M12, a phenomenon which was particularly marked for B.1.617.2., 89% of individuals neutralized B.1.1.7 and only 48% neutralized B.1.617.2 (Fig. 4b). After vaccination, 100% and 95% of convalescent individuals neutralized B.1.1.7 and B.1.617.2, respectively (Fig. 4b).

Altogether, these results indicate that B.1.617.2 displays enhanced resistance to neutralization by sera from convalescent individuals, particularly one year after infection.

### Sensitivity of B.1.617.2 to sera from vaccine recipients

We next asked whether vaccine-elicited antibodies neutralized B.1.617.2, in individuals that were not previously infected with SARS-CoV-2. We thus selected 28 individuals from a cohort of vaccinated health care workers established in Orléans. The characteristics of vaccinees are depicted in Table S2. 16 individuals received the Pfizer vaccine. Sera were sampled at week 8 (W8) (16 individuals, corresponding to week 5 after the second dose) and W16 (13 individuals, corresponding to week 13 after the second dose). 12 individuals received the AstraZeneca vaccine. In France, vaccination with Vaxzevria started on February 2021. We were thus only able to test individuals that received a single dose of this vaccine. Their sera were sampled at W10 after administration. This allowed us to assess the early humoral response to vaccination. We measured the potency of the sera against D614G, B.1.1.7, B.1.351 and B.1.617.2 (Fig. 4c).

With the Pfizer vaccine, at W8, we observed a 3-fold and 16-fold reduction in the neutralization titers against B.1.617.2 and B.1.351, respectively, when compared to B.1.1.7 (Fig. 4c). Similar differences between strains were observed at a later time point (W16), although titers were globally slightly lower globally (Fig. 4c). The situation was different with the AstraZeneca vaccine, which induced particularly low levels of antibodies neutralizing B.1.617.2 and B.1.351, when compared to D.614G and B.1.1.7 (Fig. 4c).

We also classified the vaccine recipients as neutralizers (with neutralizing antibodies detectable at 1/30 dilution) and non-neutralizers, for the four viral strains (Fig. 4c,d). With the Pfizer vaccine, most of the individuals (81 to 100%) neutralized any of the four stains at W8. This fraction remained stable at W16, with the exception of B.1.351, which was neutralized by only 46% of the individuals. 75% and 83% of individuals that received the Astrazeneca vaccine neutralized the D614G and B.1.17 strains, respectively. This fraction sharply dropped with B.1.351 and B.617.2, which were inhibited by only 8% of the sera. Therefore, the Pfizer vaccine generated a neutralizing response that efficiently targeted the four viral strains, whereas a single dose of AstraZeneca vaccine triggered a neutralizing response against D614G and B.1.1.7 but was either poorly or not at all efficient against B.1.351 and B.617.2. These results are in line with the real-world efficacy of the two vaccines. The Pfizer vaccine is known to be protective against asymptomatic and symptomatic infections, including severe diseases, induced by B.1.1.7 and B.1.351, as described for instance in the UK, Israel and South Africa ^46–48^ In contrast, the efficacy of a two-dose regimen of AstraZeneca against mild-to-moderate disease dropped to 10% in South Africa, where the B.1.351 variant circulates ^49^. Altogether, our results suggest that a single dose of AstraZeneca vaccine will not display an optimal protection against B.1.617.2.

## Discussion

After one year of an intense circulation of SARS-CoV-2 worldwide, VOCs with enhanced transmissibility and resistance to antibodies were first identified in UK, South Africa, Brazil, USA (California) and spread to many other countries. Since early 2021, India has also faced a surge of cases associated with the emergence of a new lineage termed B.1.617. It includes 3 main sublineages, B.1.617.1, B.1.617.2 and B.1.617.3. Very little is known about the epidemiological and biological characteristics of this lineage. B.1.617.2 seems to be particularly worrying, and has been deemed a VOC by multiple public health bodies including the WHO. It represents up to 80% of sequenced strains in the Maharashtra region of India ^4^. B.1.617.2 has also been detected in dozens of other countries and represented about 20% of sequenced viruses circulating in UK between May 12 and May 19, 2021 ^50^. B.1.617.2 is characterized by the presence of 9 mutations in the Spike protein. We show here that these mutations map to regions of the Spike that potentially modify virus binding to the receptor and allow escape from the humoral immune response. Our results highlight the way the virus evolves in the face of herd immunity: it modifes the sensitive ACE2 binding surface at its periphery to escape antibody recognition while still efficiently interacting with ACE2. Furthermore, our preliminary experiment showed the presence of large syncytia in B.1.617.2 infected cells. Future work will help understanding whether the mutations modify the kinetics of viral fusion and improve B.1.617.2 fitness.

We studied here the cross-reactivity of mAbs to pre-existing SARS-CoV-2 viruses, sera from long-term convalescent individuals and recent vaccine recipients against B.1.617.2. We used an authentic clinical viral isolate, rather than pseudovirus particles, to assess viral sensitivity to neutralizing antibodies. We report that some therapeutic mAb, including Bamlavinimab, lost binding to the variant Spike and no longer neutralized B.1.617.2. It is thus of importance to identify the viral strain present in patients, before administration of therapeutic mAbs in individuals at-risk for severe forms of COVID-19. We further show that B.1.617.2 is less sensitive to sera from naturally immunized individuals. The convalescent sera sampled at M6 displayed a four-fold decrease in their efficacy against B.1.617.2, as compared to B.1.1.7. This extent of decrease is similar to that observed with B.1.351, another VOC. In a second cohort of convalescent individuals sampled later (M12), this process was more marked, with a 6-fold ED50 decrease between B.1.617.2 and B.1.1.7. However, vaccination of convalescent individuals boosted the humoral immune response against all variants by more than 130-fold, well above the threshold of neutralization. These results strongly suggest that vaccination of previously infected individuals will be most likely protective against a large array of circulating viral strains, including B.1.617.2.

We report that a two-dose regimen of the Pfizer vaccine generated high sero-neutralization levels against B.1.1.7, B.1.351 and B.1.617.2 in individuals that were not previously infected with SARS-CoV-2, when sampled at W8 and W16 post vaccination. We also tested sera from individuals that received a single dose of the AstraZeneca vaccine. At W10, 75% of the sera neutralized B.1.1.7, but less than 10% of the sera neutralized B.1.351 and B.1.617.2. Neutralizing antibody levels are highly predictive of immune protection from symptomatic SARS-CoV-2 infection ^51^. Our results are in line with the negligible efficacy of a two-dose regimen of AstraZeneca vaccine against B.1.351 ^49^. A recent report analyzing data on all sequenced symptomatic cases of COVID-19 in England was used to estimate the impact of vaccination on infection with B.1.617.2 compared to B.1.1.7 ^52^. Effectiveness was notably lower with B.1.617.2 (33.5%) than with B.1.1.7 (51 %) after 1 dose of AstraZeneca vaccine. The two-dose effectiveness against B.1.617.2 was estimated to be 60% and 88% for Astrazeneca and Pfizer vaccines, respectively ^52^. Our neutralization experiments indicate that Astrazeneca vaccine-elicited antibodies are less potent against B.1.351 and B.1.617.2 than those induced by the Pfizer vaccine.

Potential limitations of our work include a relatively low number of vaccine recipients analyzed and the absence of samples from individuals that received two-doses of AstraZeneca vaccine. We have not analyzed the impact of cellular immunity, which may display higher cross-reactivity than the humoral response. Future work with more individuals and longer survey periods will help characterize the role of humoral responses in vaccine efficacy against the panel of circulating variants.

In conclusion, our results demonstrate that the novel emerging B.617.2 variant partially but significantly escapes neutralizing antibodies targeting the NTD and RBD, and polyclonal antibodies elicited by previous SARS-CoV-2 infection or vaccination.

## Methods

No statistical methods were used to predetermine sample size. The experiments were not randomized and the investigators were not blinded to allocation during experiments and outcome assessment. Our research complies with all relevant ethical regulation.

### Orléans Cohort of convalescent and vaccinated individuals

Since August 27, 2020, a prospective, monocentric, longitudinal, interventional cohort clinical study enrolling 170 SARS-CoV-2-infected individuals with different disease severities, and 30 non-infected healthy controls is on-going, aiming to describe the persistence of specific and neutralizing antibodies over a 24-months period. This study was approved by the ILE DE FRANCE IV ethical committee. At enrolment, written informed consent was collected and participants completed a questionnaire which covered sociodemographic characteristics, virological findings (SARS-CoV-2 RT-PCR results, including date of testing), clinical data (date of symptom onset, type of symptoms, hospitalization), and data related to anti-SARS-CoV-2 vaccination if ever (brand product, date of first and second vaccination). Serological status of participants was assessed every 3 months. Those who underwent anti-SARS-CoV-2 vaccination had weekly blood and nasal sampling after first dose of vaccine for a 2 months period (ClinicalTrials.gov Identifier: NCT04750720). For the present study, we selected 56 convalescent and 28 vaccinated participants (16 with Pfizer and 12 with AstraZeneca). Study participants did not receive any compensation.

### Strasbourg Cohort of convalescent individuals

Since April 2020, a prospective, interventional, monocentric, longitudinal, cohort clinical study enrolling 308 RT-PCR-diagnosed SARS-CoV-2 infected hospital staff from the Strasbourg University Hospitals is on-going (ClinicalTrials.gov Identifier: NCT04441684). At enrolment (from April 17, 2020), written informed consent was collected and participants completed a questionnaire which covered sociodemographic characteristics, virological findings (SARS-CoV-2 RT-PCR results including date of testing) and clinical data (date of symptom onset, type of symptoms, hospitalization). This study was approved by Institutional Review Board of Strasbourg University Hospital. The serological status of the participants has been described at Months 3 (M3) and Months 6 (M6) POS ^40,41^. Laboratory identification of SARS-CoV-2 was performed at least 10 days before inclusion by RT-PCR testing on nasopharyngeal swab specimens according to current guidelines (Institut Pasteur, Paris, France; WHO technical guidance). The assay targets two regions of the viral RNA-dependent RNA polymerase (RdRp) gene with a threshold of detection of 10 copies per reaction. For the present study, we randomly selected 48 patients collected at M12 (27 unvaccinated and 21 vaccinated). Study participants did not receive any compensation.

### Phylogenetic analysis

All SARS-CoV-2 sequences available on the GISAID EpiCov™ database ^53^ as of May 21, 2021 were retrieved. A subset of complete and high coverage sequences, as indicated in GISAID, assigned to lineages B.1.617.1, B.1.617.2, B.1.617.3 were randomly subsampled to contain up to 5 sequences per country and epidemiological week in R with packages *tidyverse* <u>and *lubridate*</u>. Together with a single B.1.617 sequence this subset was included in the global SARS-CoV-2 phylogeny reconstructed with augur and visualized with auspice as implemented in the Nextstrain pipeline (https://github.com/nextstrain/ncov, version from 21 May 2021) ^54^. Within Nextstrain, a random subsampling approach capping a maximum number of sequences per global region was used for the contextual non-B.1.617 sequences. The acknowledgment of contributing and originating laboratories for all sequences used in the analysis is provided in Table S3.

### 3D representation of mutations on B1.617.2 and other variants to the Spike surface

Panels in Fig. 2 were prepared with The PyMOL Molecular Graphics System, Version 2.1 Schrödinger, LLC. The atomic model used (PDB:6XR8) has been previously described ^55^.

### S-Fuse neutralization assay

U2OS-ACE2 GFP1-10 or GFP 11 cells, also termed S-Fuse cells, become GFP+ when they are productively infected by SARS-CoV-2 ^31, 21^. Cells were tested negative for mycoplasma. Cells were mixed (ratio 1:1) and plated at 8×10^3^ per well in a μClear 96-well plate (Greiner Bio-One). The indicated SARS-CoV-2 strains were incubated with mAb, sera or nasal swabs at the indicated concentrations or dilutions for 15 minutes at room temperature and added to S-Fuse cells. The nasal swabs and sera were heat-inactivated 30 min at 56°C before use. 18 hours later, cells were fixed with 2% PFA, washed and stained with Hoechst (dilution 1:1,000, Invitrogen). Images were acquired with an Opera Phenix high content confocal microscope (PerkinElmer). The GFP area and the number of nuclei were quantified using the Harmony software (PerkinElmer). The percentage of neutralization was calculated using the number of syncytia as value with the following formula: 100 × (1 – (value with serum – value in “non-infected”)/(value in “no serum” – value in “non-infected”)). Neutralizing activity of each serum was expressed as the half maximal effective dilution (ED50). ED50 values (in μg/ml for mAbs and in dilution values for sera) were calculated with a reconstructed curve using the percentage of the neutralization at the different concentrations. We previously reported a correlation between neutralization titers obtained with the S-Fuse assay and a pseudovirus neutralization assay ^56^.

### Clinical history of the patient infected with B.1.617.2

A 54-year-old man was admitted April, 27, 2021 in the Emergency department of the Hôpital Européen Georges Pompidou hospital in Paris, France, for an acute respiratory distress syndrome with fever. He had no medical background, and came back from India (West Bengali and few days spent in Delhi) 10 days before (April, 17 2021), where he stayed 15 days for his work. Onset of symptoms (abdominal pain and fever) was approximately April 18, 2021. The nasopharyngeal swab was positive SARS-CoV-2 at his date of admission. Lung tomo-densitometry showed a mild (10-25%) COVID pneumonia without pulmonary embolism. He initially received oxygen therapy 2 L/min, dexamethasone 6mg/day and enoxaparin 0,4 ml twice a day. His respiratory state worsened on day 3 (April 30, 2021). He was transferred in an intensive care unit, where he received high flow oxygen therapy (maximum 12 L/min). His respiratory condition improved, and he was transferred back in a conventional unit on day 8 (May 5, 2021). He was discharged from hospital on day 15 (May 10, 2021).

### Virus strains

The reference D614G strain (hCoV-19/France/GE1973/2020) was supplied by the National Reference Centre for Respiratory Viruses hosted by Institut Pasteur (Paris, France) and headed by Pr. S. van der Werf. This viral strain was supplied through the European Virus Archive goes Global (Evag) platform, a project that has received funding from the European Union’s Horizon 2020 research and innovation program under grant agreement n° 653316. The variant strains were isolated from nasal swabs on Vero cells and amplified by one or two passages on Vero cells. B.1.1.7 originated from a patient in Tours (France) returning from United Kingdom. B.1.351 (hCoV-19/France/IDF-IPP00078/2021) originated from a patient in Creteil (France). B.1.617.2 was isolated from a nasopharyngeal swab of a hospitalized patient returning from India. The swab was provided and sequenced by the laboratory of Virology of Hopital Européen Georges Pompidou (Assistance Publique – Hopitaux de Paris). Both patients provided informed consent for the use of the biological materials. Titration of viral stocks was performed on Vero E6, with a limiting dilution technique allowing a calculation of TCID50, or on S-Fuse cells. Viruses were sequenced directly on nasal swabs, and after one or two passages on Vero cells. Sequences were deposited on GISAID immediately after their generation, with the following IDs: D614G: EPI_ISL_414631; B.1.1.7: EPI_ISL_735391; B.1.1.351: EPI_ISL_964916; B.1.617.2: ID: EPI_ISL_2029113.

### Flow Cytometry

Vero cells with the indicated viral strains at a multiplicity of infection (MOI) of 0.1. Two days after, cells were detached using PBS-EDTA and transferred into U-bottom 96-well plates (50,000 cell/well). Cells were fixed in 4% PFA for 15-30 min at RT. Cells were then incubated for 15-30 min at RT with the indicated mAbs (1 μg/mL) in PBS, 1% BSA, 0.05% sodium azide, and 0.05% Saponin. Cells were washed with PBS, and stained using anti-IgG AF647 (1:600 dilution) (ThermoFisher). Data were acquired on an Attune Nxt instrument (Life Technologies). Stainings were also performed on control uninfected cells. Results were analysed with FlowJo 10.7.1 (Becton Dickinson).

### Antibodies

The four therapeutic antibodies were kindly provided by CHR Orleans. Human anti-SARS-CoV2 mAbs were cloned from S-specific blood memory B cells of Covid19 convalescents (Planchais et al, manuscript in preparation). Recombinant human IgG1 mAbs were produced by co-transfection of Freestyle 293-F suspension cells (Thermo Fisher Scientific) as previously described ^57^ and purified by affinity chromatography using protein G sepharose 4 fast flow beads (GE Healthcare).

### Statistical analysis

Flow cytometry data were analyzed with FlowJo v10 software (TriStar). Calculations were performed using Excel 365 (Microsoft). Figures were drawn on Prism 9 (GraphPad Software). Statistical analysis was conducted using GraphPad Prism 9. Statistical significance between different groups was calculated using the tests indicated in each figure legend.

## Data availability

All data supporting the findings of this study are available within the paper and are available from the corresponding author upon request.

## Acknowledgments

We thank Nicoletta Casartelli for critical reading of the manuscript and Pablo Guardado Calvo for discussion. We thank patients who participated to this study, members of the Virus and Immunity Unit for discussions and help, Nathalie Aulner and the UtechS Photonic BioImaging (UPBI) core facility (Institut Pasteur), a member of the France BioImaging network, for image acquisition and analysis. The Opera system was co-funded by Institut Pasteur and the Région ile de France (DIM1Health))We thank the DRCI, CIC, Médecine du travail and Pôle de Biologie teams (CHU de Strasbourg) for the management of the Strasbourg cohort and serology testing. We thank the members of the Virus and Immunity Unit for discussion and help, the UTechS Photonic BioImaging (PBI) core facility (Institut Pasteur), a member of the France BioImaging network, for image acquisition and analysis (the Opera system was co-funded by Institut Pasteur and the Région ile de France (DIM1Health))

## Author contributions

Experimental strategy design, experiments: DP, DV, AB, IS, FGB, MMR, FP, TB, ESL, FR

Vital materials DV, CP, NR, JP, MP, FG, PG, AV, JLG, LC, NKC, DE, LB, AS, HP, LH, SFK, TP, HM

Manuscript writing: DP, TB, ESL, FR, OS

Manuscript editing: DV, MMR, HP, LH, SFK, TP, HM

---

**Fig. S1. SARS-CoV-2 variants induce syncytia in S-Fuse cells.** S-Fuse cells were exposed to the indicated SARS-CoV-2 variants (MOI 10^−3^). The cells become GFP+ when they fuse together. After 20 h, infected cells were stained with anti-Spike antibodies and with Hoechst dye to visualize the nuclei. Syncytia (green), Spike (red) and nuclei (blue) are shown. Data are representative of three independent experiments. Scale bar: 50 μm.

**Fig. S2. Schematic overview of the three other variants analyzed in this study.** The consensus sequences with a focus on the Spike were built with the Sierra tool ^58^. AA modifications in comparison to the ancestral Wuhan-Hu-1 sequence (NC_045512) are indicated.

**Fig. S3. Binding of anti-SARS-CoV-2 mAbs to Vero cells infected with D614G, B.1.1.7, B.1.351 and B.1.617.2 variants.** Vero cells were infected with the indicated variants at a MOI of 0.1. After 48h, cells were stained with anti-SARS-CoV-2 mAbs (1μg/ml) and analyzed by flow-cytometry. Data are representative of three independent experiments.

**Table S1. Characteristics of the two cohorts of convalescent individuals.**

**Table S2. Characteristics of the cohort of vaccinated recipients.**

**Table S3. Contributing and originating laboratories for all sequences used in Fig. 1b.**

