## Supplementary data for "Reduced sensitivity of infectious SARS-CoV-2 variant B.1.617.2 to monoclonal antibodies and sera from convalescent and vaccinated individuals"

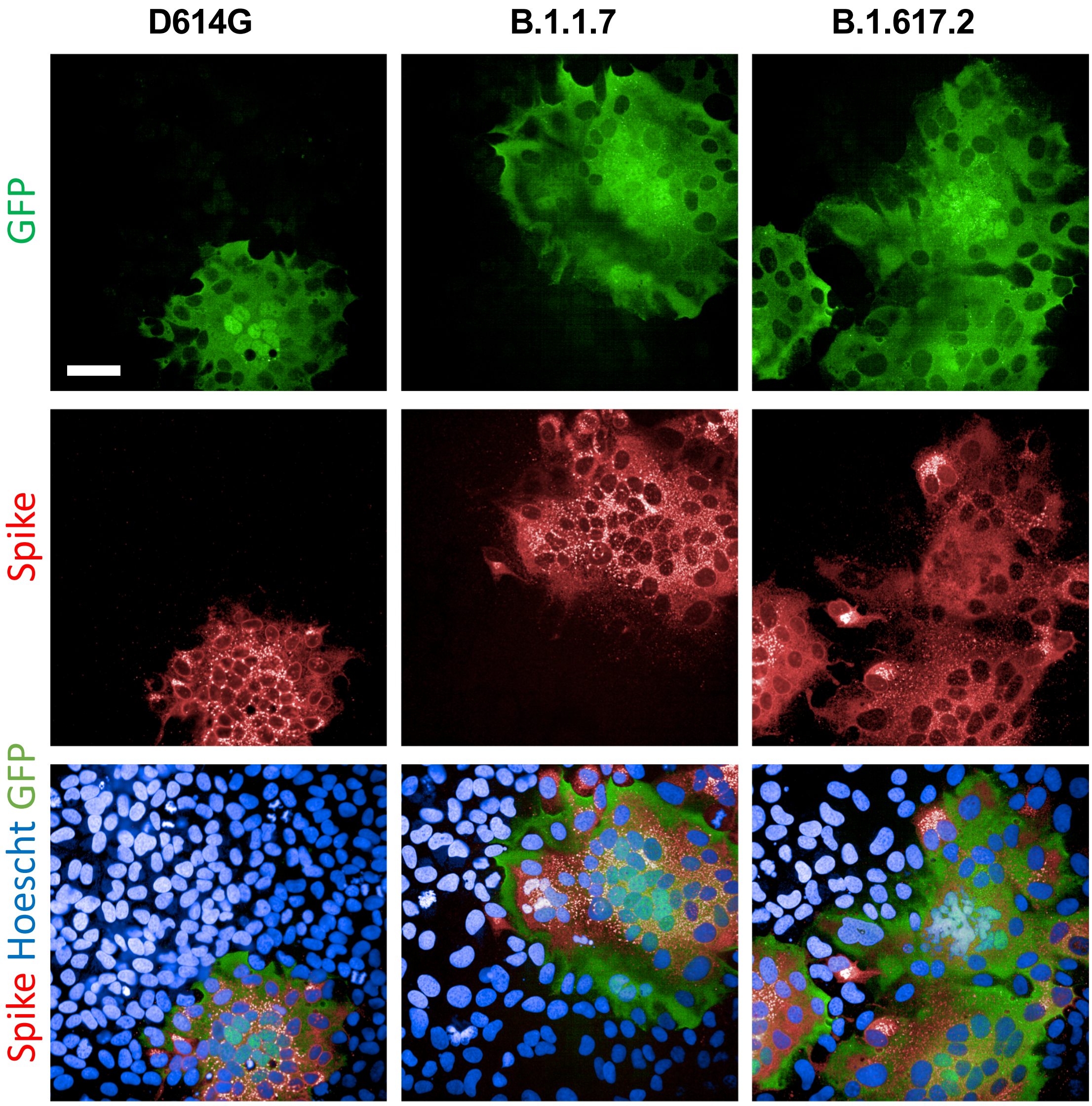


Figure S1

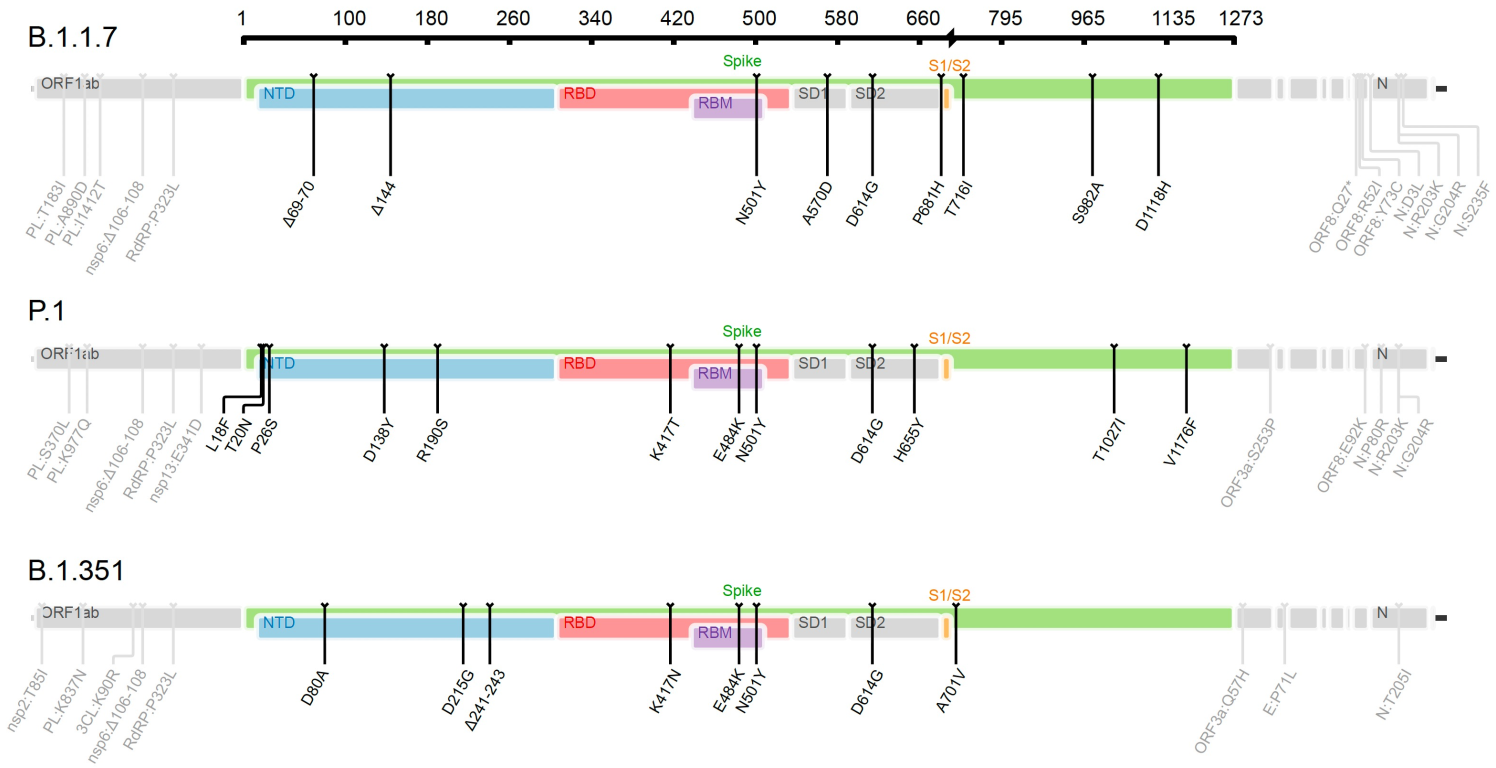
B.1.1.7
1
100
180
260
340
420
500
580
660
795
965
1135
1273
ORF1ab
Spike
NTD
RBD
RBM
SD1
SD2
S1/S2
N
PL:T183I
PL:A890D
PL:I1412T
nsp6:Δ106-108
RdRP:P323L
Δ69-70
Δ144
N501Y
A570D
D614G
P681H
T716I
S982A
D1118H
ORF8:Q27*
ORF8:R52I
ORF8:Y73C
N:D3L
N:R203K
N:G204R
N:S235F
P.1
PL:S370L
PL:K977Q
nsp6:Δ106-108
RdRP:P323L
nsp13:E341D
L18F
T20N
P26S
D138Y
R190S
K417T
E484K
N501Y
D614G
H655Y
T1027I
V1176F
ORF3a:S253P
ORF8:E92K
N:P80R
N:R203K
N:G204R
B.1.351
nsp2:T85I
PL:K837N
3CL:K90R
nsp6:Δ106-108
RdRP:P323L
D80A
D215G
Δ241-243
K417N
E484K
N501Y
D614G
A701V
ORF3a:Q57H
E:P71L
N:T205I

**Figure S2**

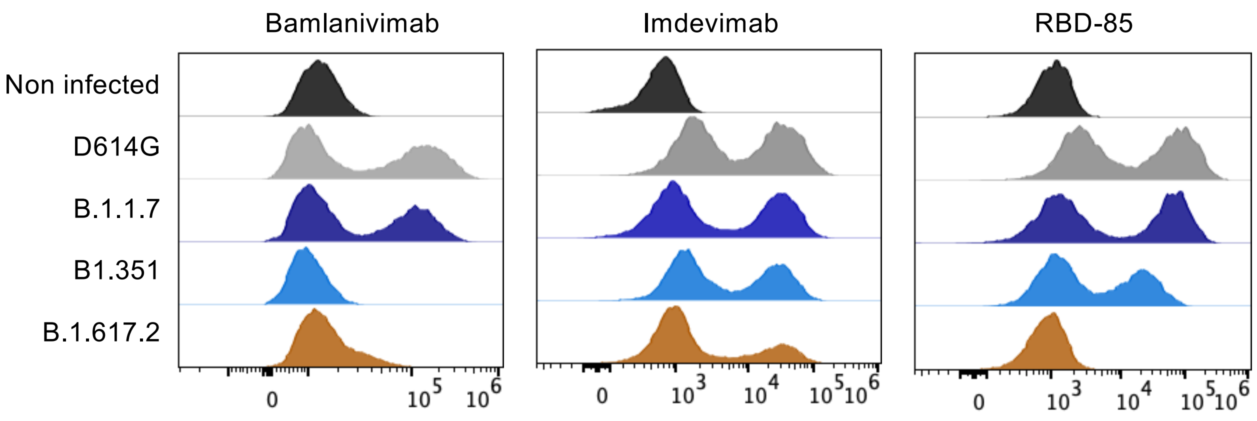
Bamlanivimab
Imdevimab
RBD-85
Non infected
D614G
B.1.1.7
B1.351
B.1.617.2
0
10^5
10^6
0
10^3
10^4
10^5
10^6
0
10^3
10^4
10^5
10^6

**Figure S3**

### a. Orléans Cohort: M6 POS

| | | n=56 |
|---|---|---|
| Sex | | |
| | Female | 29 |
| | Male | 27 |
| Age (Median; range) | | 53 (22;77) |
| Severity | | |
| | Critical | 13 |
| | Severe | 15 |
| | Mild-Moderate | 16 |
| | Asymptomatic | 12 |
| HIV | | 3 |
| PCR | | 53 |
| Anti-S (S-Flow) | | 56 |
| Sampling days POS (median; range) | | 188 (114;205) |

### b. Strasbourg Cohort : M12 POS

| | | Convalescent<br>n=26 | Vaccinated Convalescent<br>n=21 |
|---|---|---|---|
| Sex | | | |
| | Female | 21 | 18 |
| | Male | 5 | 3 |
| Age (median; range) | | 36 (24;60) | 44 (23;62) |
| Severity | | | |
| | Critical | 0 | 0 |
| | Severe | 0 | 0 |
| | Mild-Moderate | 26 | 21 |
| | Asymptomatic | 0 | 0 |
| PCR | | 26 | 21 |
| Anti-S (Abott) | | 26 | 21 |
| Vaccine | | | |
| | AstraZeneca | 0 | 9 |
| | Pfizer | 0 | 9 |
| | Moderna | 0 | 3 |
| Sampling days (median; range) | | | |
| | POS | 330 (144;383) | 359 (327;404) |
| | Post Vaccine | NA | 24 (7;81) |

**Table S1**

### Orléans cohort: Vaccinated recipients

| | | Pfizer<br>n=16 | AstraZeneca<br>n=12 |
|---|---|---|---|
| Sex | | | |
| | Female | 5 | 7 |
| | Male | 11 | 5 |
| Age (mediane; range) | | 60 (35;75) | 38.5 (29;51) |
| BMI (Mediane; Range) | | 24.2 (20;34.5) | 24.6 (21;33) |
| Immune deficiency | | 0 | 0 |
| Previous COVID-19 | | 0 | 0 |
| Anti-N | | 0 | 0 |
| 1st shot | | Jan 4- 8, 2021 | Feb 8 – March 4, 2021 |
| 2nd shot | | Jan 20 - Feb 5, 2021 | NA |
| Sampling days post-vaccination (mediane; range) | | | |
| | W8 | 57 (53;84) | |
| | W10 | | 69 (53;76) |
| | W16 | 118 (116;124) | |

**Table S2**

**Table S3 : Contributing and originating laboratories for all sequences used in Fig. 1b.**

| GISAID_accession | originating_lab | submitting_lab | authors |
|---|---|---|---|
| EPI_ISL_882760,EPI_ISL_954186 | 1.AO Universitaria 'S. Giovanni di Dio e Ruggi D'Aragona, Scuola Medica Salernitana' Hospital / 2.UOC di Virologia e Microbiologia, Università della Campania 'L. Vanvitelli' / 3.AO Universitaria 'Federico II' Napoli Hospital / 4.AORN 'San Giuseppe Moscati' Avellino Hospital / 5.AO 'San Pio - presidio G. Rummo' Benevento Hospital / 6.AO 'Sant'Anna e San Sebastiano' Caserta Hospital / | 1. Genome Research Center for Health (CRGS) / 2. Laboratory of Molecular Medicine and Genomics(LMMGe) / 3. Center for Research in Pure and Applied Mathematics (CRMPA) | Giorgio Giurato et al |
| EPI_ISL_699655,EPI_ISL_733499,EPI_ISL_733500,EPI_ISL_763067 | 1-Laboratory of Microbiology, National Reference Lab, Charles Nicolle Hospital; 2-University of Tunis ElManar, Faculty of Medicine of Tunis, LR99ES09, Tunis, Tunisia | 1-Clinical and Experimental Pharmacology Lab, LR16SP02, National Center of Pharmacovigilance, University of Tunis El Manar, Tunis, Tunisia. 2-Neurodegenerative diseases and psychiatric troubles, LR18SP03, Razi Hospital, University of Tunis El Manar, Tunis, | Ilhem Boutiba-Ben Boubaker et al |
| EPI_ISL_767931 | 4Cyte Pathology | NSW Health Pathology - Institute of Clinical Pathology and Medical Research; Westmead Hospital; University of Sydney | CIDM-PH et al. |

| | | | |
|---|---|---|---|
| EPI_ISL_517625,EPI_ISL_517616,EPI_ISL_517629,EPI_ISL_517637,EPI_ISL_517645,EPI_ISL_517646,EPI_ISL_517648,EPI_ISL_517662,EPI_ISL_518807,EPI_ISL_518815 | Academic Hospital Paramaribo | Erasmus Medical Center | Bas Oude Munnink et al |
| EPI_ISL_498479,EPI_ISL_498485,EPI_ISL_498520,EPI_ISL_498524,EPI_ISL_498534,EPI_ISL_498543,EPI_ISL_498544,EPI_ISL_1208401,EPI_ISL_1208402 | ACT Pathology | Schwessinger Lab | Ashley Jones et al |
| EPI_ISL_735503 | ACT Pathology | Schwessinger Lab | Robyn N Hall et al |

| | | | |
|---|---|---|---|
| EPI_ISL_427718 | ACT Pathology, The Canberra Hospital | NSW Health Pathology - Institute of Clinical Pathology and Medical Research; Westmead Hospital; University of Sydney | Gray K et al |
| EPI_ISL_1525226,EPI_ISL_1562451,EPI_ISL_1561855,EPI_ISL_1562503,EPI_ISL_1649291,EPI_ISL_1649351,EPI_ISL_1685730,EPI_ISL_1689967,EPI_ISL_1736936,EPI_ISL_1837612,EPI_ISL_1836347,EPI_ISL_2000472,EPI_ISL_1995160,EPI_ISL_1996453,EPI_ISL_1996867,EPI_ISL_2040187,EPI_ISL_2040461,EPI_ISL_203913 | Aegis Sciences Corporation | Centers for Disease Control and Prevention Division of Viral Diseases, Pathogen Discovery | Dakota Howard et al |
| EPI_ISL_1663516 | AIIMS, Patna | Institute of Life Sciences - INSACOG | Sunil K. Raghav et al |

| | | | |
|---|---|---|---|
| EPI_ISL_500781,EPI_ISL_2007357,EPI_ISL_2142788 | Akershus University Hospital, Department for Microbiology and Infectious Disease Control | Norwegian Institute of Public Health, Department of Virology | Kathrine Stene-Johansen et al |
| EPI_ISL_509688 | Alabama Department of Public Health Bureau of Clinical Laboratories | Pathogen Discovery, Respiratory Viruses Branch, Division of Viral Diseases, Centers for Disease Control and Prevention | Ying Tao et al |
| EPI_ISL_528530 | Alaska State Virology Laboratory | Alaska State Virology Laboratory | Chen J et al with Pathogenomics group Dagdag R et al |

| | | | |
|---|---|---|---|
| EPI_ISL_586258,EPI_ISL_586262 | Alaska State Virology Laboratory | Alaska State Virology Laboratory | Jack Chen et al |
| EPI_ISL_911693,EPI_ISL_1039732,EPI_ISL_1039716,EPI_ISL_1049507,EPI_ISL_1675113 | Alaska State Virology Laboratory | Alaska State Virology Laboratory | Stephanie DeRonde et al |
| EPI_ISL_805733,EPI_ISL_805749,EPI_ISL_806372 | Alberta Precision Labs (APL) | Alberta Precision Labs (APL) | Gordon P et al |

| | | | |
|---|---|---|---|
| EPI_ISL_2100700 | All India Institute of Medical Sciences, Ansari Nagar Delhi | CSIR-Institute of Genomics and Integrative Biology | Pooja Sharma* et al |
| EPI_ISL_739661 | Al-Quds Nutrition and Health Research Institute, Al-Quds University | Al-Quds Nutrition and Health Research Institute, Al-Quds University | Nasereddin et al |
| EPI_ISL_528707 | Alsafar - Khalifa University Abu Dhabi | Alsafar - Khalifa University Abu Dhabi | Andreas Henschel et al |

| | | | |
|---|---|---|---|
| EPI_ISL_1015796 | Altius Institute for Biomedical Sciences | Seattle Flu Study | Deborah A. Nickerson et al |
| EPI_ISL_1445266 | AMBULATORIO MEDICO DE ESPECIALIDADES DE PERUIBE | Instituto Butantan / Mendelics | Dimas Tadeu Covas et al |
| EPI_ISL_467455 | AMPATH-DBN | KRISP, KZN Research Innovation and Sequencing Platform | Giandhari J et al |

| | | | |
|---|---|---|---|
| EPI_ISL_1165068 | Anwar Medika General Hospital | Institute of Tropical Disease, Universitas Airlangga | Jezzy R Dewantari et al |
| EPI_ISL_576196 | AR Dept. of Health-Public Health Lab | Pathogen Discovery, Respiratory Viruses Branch, Division of Viral Diseases, Centers for Disease Control and Prevention | Ying Tao et al |
| EPI_ISL_770026 | Area De Salud Catedral Noreste | Inciensa, Instituto Costarricense de Investigación y Enseñanza en Nutrición y Salud | Francisco Duarte et al |

| | | | |
|---|---|---|---|
| EPI_ISL_770014 | Area De Salud Curridabat 2 | Inciensa, Instituto Costarricense de Investigación y Enseñanza en Nutrición y Salud | Francisco Duarte et al |
| EPI_ISL_512662,EPI_ISL_527760,EPI_ISL_770003 | Area De Salud La Cruz | Inciensa, Instituto Costarricense de Investigación y Enseñanza en Nutrición y Salud | Francisco Duarte et al |
| EPI_ISL_1196423 | AREA DE SALUD MATA REDONDA-HOSPITAL - CLINICA DR. MORENO CAÑAS | Inciensa, Instituto Costarricense de Investigación y Enseñanza en Nutrición y Salud | Francisco Duarte et al |

| | | | |
|---|---|---|---|
| EPI_ISL_770005 | Area De Salud Moravia | Inciensa, Instituto Costarricense de Investigación y Enseñanza en Nutrición y Salud | Francisco Duarte et al |
| EPI_ISL_1827544 | AREA DE SALUD SANTA CRUZ | Inciensa, Instituto Costarricense de Investigación y Enseñanza en Nutrición y Salud | Francisco Duarte et al |
| EPI_ISL_500657,EPI_ISL_500703,EPI_ISL_509522 | Area of Virology, Serology and Virology Division (SAViD), New South Wales Health Pathology Randwick | Area of Virology, Serology and Virology Division (SAViD), New South Wales Health Pathology Randwick | Rawlinson et al |

| | | | |
|---|---|---|---|
| EPI_ISL_678310,EPI_ISL_678312,EPI_ISL_678386,EPI_ISL_1005538,EPI_ISL_1005536,EPI_ISL_1293047,EPI_ISL_1494721,EPI_ISL_1543925,EPI_ISL_1615598,EPI_ISL_1615596,EPI_ISL_1615595,EPI_ISL_1633353,EPI_ISL_1633348,EPI_ISL_1911191,EPI_ISL_1911189 | Area of Virology, Serology and Virology Division (SAViD), New South Wales Health Pathology Randwick | Virology Research Laboratory; Area of Virology, Serology and Virology Division (SAViD), New South Wales Health Pathology Randwick | Foster et al |
| EPI_ISL_406223 | Arizona Department of Health Services | Pathogen Discovery, Respiratory Viruses Branch, Division of Viral Diseases, Centers for Disease Control and Prevention | Ying Tao et al |
| EPI_ISL_978405 | Arizona State Public Health Laboratory | Arizona State Public Health Laboratory | Trung Huynh et al |

| | | | |
|---|---|---|---|
| EPI_ISL_1595843 | Armed Forces Institute of Pathology (AFIP), Dhaka Cantonment | Genomic Research Lab, BCSIR | Md. Murshed Hasan Sarkar et al |
| EPI_ISL_1615663 | Armed Forces Institute of Pathology (AFIP), Dhaka Cantonment | Genomic Research Lab, BCSIR | Mohammad Mizanur Rahman et al |
| EPI_ISL_1593860 | Armed Forces Institute of Pathology (AFIP), Dhaka Cantonment | Genomic Research Lab, BCSIR | Susane Giti et al |

| | | | | |
|---|---|---|---|---|
| RCGEB - MASA et al | ASL Napoli 1 Centro | AMES Centro Polidiagnostico Strumentale S.r.l. | Giovanni Savarese et alEPI_ISL_1080712 Atlanta VA Medical CenterPathogen Discovery, Respiratory Viruses Branch, Division of Viral Diseases, Centers for Disease Control and PreventionYan Li et alEPI_ISL_1591248 Auckland HospitalInstitute of Environmental Science and Research (ESR)Matt Storey et alEPI ISL 413490 | EPI_ISL_516431 |
| EPI_ISL_2103642,EPI_ISL_2103689 | Clinical Microbiology, Infection Prevention and Control | Section for Molecular Diagnostics | Björn Hallström et al | |
| EPI_ISL_877630 | Clinical Molecular Microbiology Laboratory, UNC Hospital | Dirk Dittmer | Razia Moorad et al | |

| | | | |
|---|---|---|---|
| EPI_ISL_2023324 | Clinical Reference Laboratory | Kansas Health and Environmental Lab | Mike Grose et al |
| EPI_ISL_1833679 | Clinical Virology | Clinical Bacteriology | Tim Roloff et al |
| EPI_ISL_2035940 | Clinical Virology | Clinical Virology | Wasfi Fares et al |

| | | | |
|---|---|---|---|
| EPI_ISL_447408 | Clinical Virology Unit, Hadassah Hebrew University Medical Center | Stern Lab | Stern Lab et al |
| EPI_ISL_2151071 | Clínica Meta | Instituto de Virología-Universidad El Bosque | L. Johana Madroñero et al |
| EPI_ISL_445361 | CLINICA UC SAN CARLOS DE APOQUINDO | Instituto de Salud Publica de Chile | Andrés E Castillo et al |

| | | | |
|---|---|---|---|
| EPI_ISL_1494958 | Clinica Universitaria Medicina Integral - Laboratorio Molecular | Instituto Nacional de Salud- Dirección de Investigación en Salud Pública | Katherine Laiton-Donato et al |
| EPI_ISL_1599177 | Cliniques universitaires Saint-Luc | UCLouvain/IREC/MBLG | Jean Ruelle et al |
| EPI_ISL_676567,EPI_ISL_1788257 | CNR Virus des Infections Respiratoires - France SUD | CNR Virus des Infections Respiratoires - France SUD | Antonin Bal et al |

| | | | |
|---|---|---|---|
| EPI_ISL_1539438 | Colorado Department of Public Health and Environment | Colorado Department of Puplic Health and Environment | Laura Bankers et al |
| EPI_ISL_710321,EPI_ISL_1038842 | Colorado Department of Public Health and Environment | Colorado Department of Puplic Health and Environment | Laura Bankers et al |
| EPI_ISL_527466 | Colorado State University - Ebel Lab | Colorado State University - Ebel Lab | Greg Ebel et al. |
| EPI_ISL_1169508,EPI_ISL_1169518,EPI_ISL_1169542 | Commonwealth Healthcare Center | Genomics and Discovery, Respiratory Viruses Branch, Division of Viral Diseases, Centers for Disease Control and Prevention | Krista Queen et al |
| EPI_ISL_682300,EPI_ISL_682315,EPI_ISL_1660431,EPI_ISL_1660428,EPI_ISL_1660424,EPI_ISL_1660457,EPI_ISL_1660462,EPI_ISL_1660464,EPI_ISL_1660466 | Communicable Disease Laboratory, Public Health Directorate | Communicable Disease Laboratory, Public Health Directorate | Alwasti et al |

| | | | |
|---|---|---|---|
| EPI_ISL_632250,EPI_ISL_632284,EPI_ISL_632263 | Communicable Disease Laboratory, Public Health Directorate | Communicable Disease Laboratory, Public Health Directorate | AlWasti et al |
| EPI_ISL_845620 | compensar calle 63 | Instituto Nacional de Salud - Dirección de Investigación en Salud Pública | Katherine Laiton-Donato et al |
| EPI_ISL_738246,EPI_ISL_738312 | Connecticut Veterans' Affairs Hospital | Grubaugh Lab - Yale School of Public Health | Joseph Fauver et al |
| EPI_ISL_1017597,EPI_ISL_1017613 | Connecticut Veterans' Affairs Hospital | Grubaugh Lab - Yale School of Public Health | Mary Petrone et al |
| EPI_ISL_1040821 | CoVid WC Cape Town Metro | NHLS/UCT | Arash Iranzadeh et al |
| EPI_ISL_910144,EPI_ISL_910281 | CSIR-Centre for Cellular and Molecular Biology | CSIR-Centre for Cellular and Molecular Biology | Payel Mukherjee et al |
| EPI_ISL_539771 | CSIR-Centre for Cellular and Molecular Biology | CSIR-Centre for Cellular and Molecular Biology | Pratheusa Maccha et al |
| EPI_ISL_2156965,EPI_ISL_2156966 | CSIR-Centre for Cellular and Molecular Biology | CSIR-Centre for Cellular and Molecular Biology-INSACOG | Ara Sreenivas et al |
| EPI_ISL_2156869,EPI_ISL_2156882,EPI_ISL_2156946,EPI_ISL_2156951 | CSIR-Centre for Cellular and Molecular Biology | CSIR-Centre for Cellular and Molecular Biology-INSACOG | Blessy B John et al |
| EPI_ISL_2157041,EPI_ISL_2157048 | CSIR-Centre for Cellular and Molecular Biology | CSIR-Centre for Cellular and Molecular Biology-INSACOG | Lamuk Zaveri et al |
| EPI_ISL_1838377,EPI_ISL_1838545,EPI_ISL_1838552,EPI_ISL_1838604,EPI_ISL_1838655,EPI_ISL_1838670,EPI_ISL_1838761,EPI_ISL_1838764,EPI_ISL_1838820 | CSIR-Centre for Cellular and Molecular Biology | CSIR-Centre for Cellular and Molecular Biology-INSACOG | Payel Mukherjee et al |
| EPI_ISL_2156990,EPI_ISL_2157061 | CSIR-Centre for Cellular and Molecular Biology | CSIR-Centre for Cellular and Molecular Biology-INSACOG | Tulasi Nagabandi et al |
| EPI_ISL_1360316,EPI_ISL_1360361,EPI_ISL_1916473,EPI_ISL_1916475 | CSIR-National Environmental Engineering Research Institute | CSIR-Centre for Cellular and Molecular Biology - INSACOG | Ara Sreenivas et al |
| EPI_ISL_1360306,EPI_ISL_1360329,EPI_ISL_1360330,EPI_ISL_1360338 | CSIR-National Environmental Engineering Research Institute | CSIR-Centre for Cellular and Molecular Biology - INSACOG | Lamuk Zaveri et al |

| | | | |
|---|---|---|---|
| EPI_ISL_1360304,EPI_ISL_1360352 | CSIR-National Environmental Engineering Research Institute | CSIR-Centre for Cellular and Molecular Biology - INSACOG | Onkar Kulkarni et al |
| EPI_ISL_1360318,EPI_ISL_1360317,EPI_ISL_1360342,EPI_ISL_1360364 | CSIR-National Environmental Engineering Research Institute | CSIR-Centre for Cellular and Molecular Biology - INSACOG | Sofia Banu et al |
| EPI_ISL_722139 | DB Diagnosticos do Brasil | Laboratório de Parasitologia Médica - Instituto de Medicina Tropical - Universidade de São Paulo | Brazil-UK Centre for Arbovirus Discovery Diagnosis Genomics et al |
| EPI_ISL_804816 | DB Diagnosticos do Brasil | Laboratório de Parasitologia Médica - Instituto de Medicina Tropical - Universidade de São Paulo | Nuno Faria et al |
| EPI_ISL_476365 | DB Diagnósticos do Brasil | Instituto de Medicina Tropical da Univesidade de São Paulo | Samples: Nelson Gaburo Jr et al |
| EPI_ISL_2018399 | DC Public Health Lab/ Dept. of Forensic Sciences | Centers for Disease Control and Prevention Division of Viral Diseases, Pathogen Discovery | Mili Sheth et al |
| EPI_ISL_1502075 | DC Public Health Lab/ Dept. of Forensic Sciences | DC Public Health Lab/ Dept. of Forensic Sciences | Janis Doss et al |
| EPI_ISL_1168445 | DC Public Health Lab/ Dept. of Forensic Sciences | DC Public Health Lab/ Dept. of Forensic Sciences | Scott Nguyen et al |
| EPI_ISL_826923,EPI_ISL_826729,EPI_ISL_828668,EPI_ISL_828691,EPI_ISL_829136,EPI_ISL_830194,EPI_ISL_829701 | deCODE genetics | deCODE genetics | Daniel F Gudbjartsson et al |
| EPI_ISL_560406,EPI_ISL_1137042 | Delaware Public Health Lab | Delaware Public Health Lab | Gregory Hovan et al |
| EPI_ISL_693711,EPI_ISL_767006 | Delaware Public Health Laboratory | Delaware Public Health Laboratory | Gregory Hovan et al |

| | | | |
|---|---|---|---|
| EPI_ISL_2100645 | Delhi North District | CSIR-Institute of Genomics and Integrative Biology | Pooja Sharma* et al |
| EPI_ISL_430802 | Departamento de Biología y genética molecular, IACA Laboratorios. | Área de Secuenciación del Laboratorio de Virología del Hospital de Niños Dr. Ricardo Gutierrez on behalf of 'Proyecto Argentino Interinstitucional de genomica de SARS-CoV-2' (PAIS Consortium) | Nabaes Jodar et al |
| EPI_ISL_1340750,EPI_ISL_1340751,EPI_ISL_1340752,EPI_ISL_1340753,EPI_ISL_1340754,EPI_ISL_1340755,EPI_ISL_1340758,EPI_ISL_1340759,EPI_ISL_1340760,EPI_ISL_1340761,EPI_ISL_1340764 | Departamento de Virologia, Laboratorio Central de Salud Pública, Avenida Venezuela y Teniente Escurra, Asunción, Paraguay | Laboratory of Respiratory Viruses and Measles, Oswaldo Cruz Institute, FIOCRUZ | Paola Resende et al |
| EPI_ISL_1137609 | Département de Maladies Infectieuses, CHU Farhat Hached Sousse, Tunisie | Laboratoire des Procédés de Criblage Moléculaire et Cellulaire-Centre de Biotechnologie de Sfax | Souissi et al |
| EPI_ISL_516930 | Department for Molecular Diagnostics, Centre for Medical Microbiology, Institute of Public Health of Montenegro | Charité Universitätsmedizin Berlin, Institut für Virologie | Victor M Corman et al |
| EPI_ISL_481380 | Department for Virology, Molecular Biology and Genome Research, R. G. Lugar Center for Public Health Research, National Center for Disease Control and Public Health (NCDC) of Georgia. | Department for Virology, Molecular Biology and Genome Research, R. G. Lugar Center for Public Health Research, National Center for Disease Control and Public Health (NCDC) of Georgia. | Ana Papkiauri et al |

| | | | |
|---|---|---|---|
| EPI_ISL_1914785 | Department for Virology, Molecular Biology and Genome Research, R. G. Lugar Center for Public Health Research, National Center for Disease Control and Public Health (NCDC) of Georgia. | Department for Virology, Molecular Biology and Genome Research, R. G. Lugar Center for Public Health Research, National Center for Disease Control and Public Health (NCDC) of Georgia. | Giorgi Gogoladze et al |
| EPI_ISL_763062,EPI_ISL_1914604 | Department for Virology, Molecular Biology and Genome Research, R. G. Lugar Center for Public Health Research, National Center for Disease Control and Public Health (NCDC) of Georgia. | Department for Virology, Molecular Biology and Genome Research, R. G. Lugar Center for Public Health Research, National Center for Disease Control and Public Health (NCDC) of Georgia. | Giorgi Tomashvili et al |
| EPI_ISL_470877,EPI_ISL_1048367 | Department for Virology, Molecular Biology and Genome Research, R. G. Lugar Center for Public Health Research, National Center for Disease Control and Public Health (NCDC) of Georgia. | Department for Virology, Molecular Biology and Genome Research, R. G. Lugar Center for Public Health Research, National Center for Disease Control and Public Health (NCDC) of Georgia. | Gvantsa Brachveli et al |
| EPI_ISL_471529,EPI_ISL_754181,EPI_ISL_1921859,EPI_ISL_1914783 | Department for Virology, Molecular Biology and Genome Research, R. G. Lugar Center for Public Health Research, National Center for Disease Control and Public Health (NCDC) of Georgia. | Department for Virology, Molecular Biology and Genome Research, R. G. Lugar Center for Public Health Research, National Center for Disease Control and Public Health (NCDC) of Georgia. | Meri Pantsulaia et al |

| | | | |
|---|---|---|---|
| EPI_ISL_754180 | Department for Virology, Molecular Biology and Genome Research, R. G. Lugar Center for Public Health Research, National Center for Disease Control and Public Health (NCDC) of Georgia. | Department for Virology, Molecular Biology and Genome Research, R. G. Lugar Center for Public Health Research, National Center for Disease Control and Public Health (NCDC) of Georgia. | Salome Javashvili et al |
| EPI_ISL_1914578 | Department for Virology, Molecular Biology and Genome Research, R. G. Lugar Center for Public Health Research, National Center for Disease Control and Public Health (NCDC) of Georgia. | Department for Virology, Molecular Biology and Genome Research, R. G. Lugar Center for Public Health Research, National Center for Disease Control and Public Health (NCDC) of Georgia. | Tata Imnadze et al |
| EPI_ISL_515085,EPI_ISL_515096,EPI_ISL_515103,EPI_ISL_515105 | Department of Biochemistry, Cell and Molecular Biology | WACCBIP, University of Ghana | Ngoi et al |
| EPI_ISL_944681,EPI_ISL_944701 | Department of Biochemistry, Cell and Molecular Biology, West African Centre for Cell Biology of Infectious Pathogens (WACCBIP), University of Ghana | Department of Biochemistry, Cell and Molecular Biology, West African Centre for Cell Biology of Infectious Pathogens (WACCBIP), University of Ghana | Morang'a et al |
| EPI_ISL_884850,EPI_ISL_884838,EPI_ISL_884841 | Department of Biochemistry, Cell and Molecular Biology, West African Centre for Cell Biology of Infectious Pathogens (WACCBIP), University of Ghana | Department of Biochemistry, Cell and Molecular Biology, West African Centre for Cell Biology of Infectious Pathogens (WACCBIP), University of Ghana | Ngoi et al |
| EPI_ISL_907075,EPI_ISL_956332 | Department of Biology, University of Basrah | Department of Biology, University of Basrah | Abu-Ali et al |
| EPI_ISL_484708,EPI_ISL_581664,EPI_ISL_661247,EPI_ISL_722965,EPI_ISL_737349 | Department of Clinical Microbiology | GIGA Medical Genomics | Keith Durkin et al |

| | | | |
|---|---|---|---|
| EPI_ISL_1805696 | Department of Genetic Engineering and Biotechnology, Shahjalal University of Science and Technology | Genomic Research Lab, BCSIR | Md. Murshed Hasan Sarkar et al |
| EPI_ISL_1280136 | Department of Genetics, Medirex | Laboratory of Genomics and Bioinformatics, Comenius University Science Park | Tatiana Sedláčková et al |
| EPI_ISL_417444 | Department of Healthcare Biotechnology, National University of Sciences and Technology (NUST) | Department of Healthcare Biotechnology, National University of Sciences and Technology (NUST) | Javed et al |
| EPI_ISL_1018325,EPI_ISL_1797619,EPI_ISL_1797620,EPI_ISL_1797622,EPI_ISL_1797623,EPI_ISL_1797624,EPI_ISL_1797625,EPI_ISL_1797626 | Department of Health Technology and Informatics, The Hong Kong Polytechnic University | Department of Health Technology and Informatics, The Hong Kong Polytechnic University | Gilman Kit-Hang Siu et al |
| EPI_ISL_1920640 | Department of Hygiene, Epidemiology and Medical Statistics, Medical School, National and Kapodistrian University of Athens | Central Public Health Laboratory, National Public Health Organization | Gkikas Magiorkinis et al |
| EPI_ISL_410984 | Department of Infectious and Tropical Diseases, Bichat Claude Bernard Hospital, Paris | National Reference Center for Viruses of Respiratory Infections, Institut Pasteur, Paris | Mélanie Albert et al |
| EPI_ISL_1942010 | Department of Infectious Diseases, Kobe Institute of Health | Department of Infectious Diseases, Kobe Institute of Health | Ryohei Nomoto et al |
| EPI_ISL_1306140,EPI_ISL_1306146 | Department of Laboratory Medicine, Clinical Center, National Institutes of Health | Laboratory of Parasitic Diseases, Systems Genomics Section, National Institute of Allergy and Infectious Diseases, National Institutes of Health | Allison Roder et al |

| | | | |
|---|---|---|---|
| EPI_ISL_934571,EPI_ISL_1008368,EPI_ISL_2137114 | Department of Laboratory Medicine, Division of Clinical Virology, University of Medicine, Vienna | Bergthaler laboratory, CeMM Research Center for Molecular Medicine of the Austrian Academy of Sciences | Lukas Endler et al |
| EPI_ISL_447621,EPI_ISL_463007,EPI_ISL_534336,EPI_ISL_1010729,EPI_ISL_1020316,EPI_ISL_1039160,EPI_ISL_1041957,EPI_ISL_1667469,EPI_ISL_1667472 | Department of Laboratory Medicine, National Taiwan University Hospital | Microbial Genomics Core Lab, National Taiwan University Centers of Genomic and Precision Medicine | Shiou-Hwei Yeh et al |
| EPI_ISL_477175 | Department of Laboratory Medicine Tan Tock Seng Hospital | Department of Laboratory Medicine Tan Tock Seng Hospital | Chen YYC et al |
| EPI_ISL_648106,EPI_ISL_648667,EPI_ISL_648685,EPI_ISL_648724,EPI_ISL_648782,EPI_ISL_648725 | Department of Laboratory Medicine, Tan Tock Seng Hospital | Department of Laboratory Medicine, Tan Tock Seng Hospital | Chen YYC et al |
| EPI_ISL_477170 | Department of Laboratory, Medicine Tan Tock Seng Hospital | Department of Laboratory Medicine Tan Tock Seng Hospital | Chen YYC et al |
| EPI_ISL_1678790,EPI_ISL_1678810,EPI_ISL_1678636 | Department of Medical Microbiology & Infection prevention, Amsterdam University Medical Centers location AMC | Department of Medical Microbiology & Infection prevention, Amsterdam University Medical Centers location AMC | Matthijs Welkers et al |
| EPI_ISL_1919364 | Department of Medical Microbiology - section Molde, Molde Hospital | Norwegian Institute of Public Health, Department of Virology | Kathrine Stene-Johansen et al |
| EPI_ISL_501215,EPI_ISL_501196 | Department of Medical Microbiology, University Malaya Medical Centre | Department of Medical Microbiology, Faculty of Medicine, University of Malaya | Yoong Min CHONG et al |
| EPI_ISL_1716719 | Department of Microbiology | Greek Genome Center, Biomedical Research Foundation of the Academy of Athens (BRFAA) | Emmanouil Athanasiadis et al |

| | | | |
|---|---|---|---|
| EPI_ISL_1745698 | Department of Microbiology, Faculty of Medicine and Dentistry, Palacky University and University Hospital Olomouc | Institute of Molecular and Translational Medicine / Laboratory of Experimental Medicine, Faculty of Medicine and Dentistry, Palacky University | Rastislav Slavkovský et al |
| EPI_ISL_1027639,EPI_ISL_1027648 | Department of Microbiology, National Institute for Public Health of Kosova | Charité Universitätsmedizin Berlin, Institut für Virologie | Victor M Corman et al |
| EPI_ISL_498270,EPI_ISL_497795,EPI_ISL_1034373,EPI_ISL_1034641,EPI_ISL_1034718,EPI_ISL_1055454,EPI_ISL_1197084 | Department of Microbiology, The University of Hong Kong | Department of Microbiology, The University of Hong Kong | Kelvin K.W. To et al |
| EPI_ISL_1008285 | Department of Microbiology, University Innsbruck | Bergthaler laboratory, CeMM Research Center for Molecular Medicine of the Austrian Academy of Sciences | Lukas Endler et al |
| EPI_ISL_463748,EPI_ISL_1164627,EPI_ISL_1164628,EPI_ISL_1164644,EPI_ISL_1164704,EPI_ISL_1164673,EPI_ISL_1164716 | Department of Molecular Virology, Cyprus Institute of Neurology and Genetics | Department of Molecular Virology, Cyprus Institute of Neurology and Genetics | Jan Richter et al |
| EPI_ISL_594187,EPI_ISL_596452,EPI_ISL_596455 | Department of Pathology, School of Medicine, Imam Khomeini Hospital, Tehran University of Medical Sciences | Genetics Research Center, University of Social Welfare and Rehabilitation Sciences | Zohreh Fattahi et al |
| EPI_ISL_2099869 | Department of Public Health Bucharest | National Institute of Infectious Diseases-Prof. Dr. Matei Bals Molecular Diagnostics Laboratory | Corina Casangiu et al |

| | | | |
|---|---|---|---|
| EPI_ISL_1034300,EPI_ISL_1289910 | Department of Public Health Microbiology Ljubljana, National Laboratory for Health, Environment and Food | Department of Public Health Microbiology Ljubljana, National Laboratory for Health, Environment and Food | José Gonçalves et al |
| EPI_ISL_2032161 | Department of Public Health Microbiology Ljubljana, National Laboratory for Health, Environment and Food | Department of Public Health Microbiology Ljubljana, National Laboratory for Health, Environment and Food | Tom Koritnik et al |
| EPI_ISL_582509 | Department of Respiratory and other Viral Infections of L.V.Gromashevsky Institute of Epidemiology & Infectious Diseases NAMS of Ukraine | Department of Respiratory and other Viral Infections of L.V.Gromashevsky Institute of Epidemiology & Infectious Diseases NAMS of Ukraine, JSC "Farmak" | Alla Mironenko et al |
| EPI_ISL_1122014 | Department of Respiratory and other Viral Infections of L.V.Gromashevsky Institute of Epidemiology & Infectious Diseases NAMS of Ukraine | Department of Respiratory and other Viral Infections of L.V.Gromashevsky Institute of Epidemiology & Infectious Diseases NAMS of Ukraine, JSC "Farmak" | Alla Mironenko et al |
| EPI_ISL_576147,EPI_ISL_576149 | Department of Respiratory & Other Viral Infections of L.V. Gromashevsky Institute of Epidemiology & Infectious Diseases NAMS of Ukraine | Department of Respiratory & Other Viral Infections of L.V. Gromashevsky Institute of Epidemiology & Infectious Diseases NAMS of Ukraine, JSC "Farmak" | Alla Mironenko et al |
| EPI_ISL_450200 | Department of Virology | Department of Virology | Boehmer et al |
| EPI_ISL_1385786,EPI_ISL_1385794,EPI_ISL_1397960,EPI_ISL_1406394,EPI_ISL_1385791,EPI_ISL_1385784,EPI_ISL_1710510,EPI_ISL_1732274 | Department of Virology | Department of Virology | Massab Umair et al |

| | | | |
|---|---|---|---|
| EPI_ISL_732642,EPI_ISL_737211,EPI_ISL_757291,EPI_ISL_757339,EPI_ISL_757378,EPI_ISL_862055,EPI_ISL_862062,EPI_ISL_862093,EPI_ISL_1082032,EPI_ISL_1219740,EPI_ISL_1231217,EPI_ISL_1496876,EPI_ISL_1842257 | Department of Virology and Immunology, University of Helsinki and Helsinki University Hospital, Huslab Finland | Department of Virology, Faculty of Medicine, University of Helsinki, Helsinki, Finland | Teemu Smura et al |
| EPI_ISL_407084 | Department of Virology III, National Institute of Infectious Diseases | Pathogen Genomics Center, National Institute of Infectious Diseases | Tsuyoshi Sekizuka et al |
| EPI_ISL_1015173 | Department of Virology, Pitié-Salpêtrière hospital | Department of Virology, Pitié-Salpêtrière hospital | Valentin Leducq et al |
| EPI_ISL_855565,EPI_ISL_855572 | Department of Virology, Principal Military Hospital of Instruction of Tunis | Bundeswehr Institute of Microbiology | Susann Handrick et al |
| EPI_ISL_929069,EPI_ISL_1890934,EPI_ISL_1876278,EPI_ISL_1868349,EPI_ISL_1872528,EPI_ISL_1869673,EPI_ISL_1885850,EPI_ISL_1871129,EPI_ISL_1885762,EPI_ISL_1874455,EPI_ISL_1880910,EPI_ISL_2027204,EPI_ISL_2026084,EPI_ISL_2026527,EPI_ISL_2026828,EPI_ISL_2028113,EPI_ISL_2027848,EPI_ISL_2024848,EPI_ISL_2025094,EPI_ISL_2024994,EPI_ISL_2024762 | Department of Virus and Microbiological Special Diagnostics, Statens Serum Institut, Copenhagen, Denmark | Aalborg University | Danish Covid-19 Genome Consortium et al |
| EPI_ISL_670146,EPI_ISL_714931 | Department of Virus and Microbiological Special Diagnostics, Statens Serum Institut, Copenhagen, Denmark | Albertsen Lab, Department of Chemistry and Bioscience, Aalborg University, Denmark | Danish Covid-19 Genome Consortium et al |

| | | | |
|---|---|---|---|
| EPI_ISL_429450 | Department of Virus and Microbiological Special Diagnostics, Statens Serum Institut, Copenhagen, Denmark, Artillerivej 5, 2300 Copenahgen S | Albertsen lab, Department of Chemistry and Bioscience, Aalborg University, Denmark | Rasmus Kirkegaard et al |
| EPI_ISL_618919 | Department of Virus and Microbiological Special Diagnostics, Statens Serum Institut, Denmark | Albertsen lab, Department of Chemistry and Bioscience, Aalborg University, Denmark | Danish Covid-19 Genome Consortia et al |
| EPI_ISL_1118134,EPI_ISL_1994923 | Dept. of Microbiology and Infection Control, Akershus University Hospital HF | Dept. of Microbiology and Infection Control, Akershus University Hospital HF | Hege Vangstein Aamot et al |
| EPI_ISL_410532,EPI_ISL_410531 | Dept. of Pathology, National Institute of Infectious Diseases | Pathogen Genomics Center, National Institute of Infectious Diseases | Tsuyoshi Sekizuka et al |
| EPI_ISL_408667 | Dept. of Virology III, National Institute of Infectious Diseases | Pathogen Genomics Center, National Institute of Infectious Diseases | Tsuyoshi Sekizuka et al |
| EPI_ISL_774933 | Designated Reference Institute for Chemical Measurements (DRICM) | DNA SOLUTION LTD. | Md. Imran Khan et al |
| EPI_ISL_1447336 | Diagnostic and Research Center of Infectious Diseases, Medical Faculty, Andalas University | Diagnostic and Research Center of Infectious Diseases, Medical Faculty, Andalas University | Andani Eka Putra et al |
| EPI_ISL_2140367 | Diagnostyka Sp. z o.o. | 1. Tricity SARS-CoV-2 sequencing consortium: University of Gdansk, Medical University of Gdansk, Vaxican Ltd., Invicta Ltd. 2. National Institute of Public Health - National Institute of Hygiene, Warsaw, Poland | Maciej Kosinski et al |
| EPI_ISL_1145662,EPI_ISL_2115136 | Dianovis GmbH Greiz | Robert Koch Institute | ? |

| | | | |
|---|---|---|---|
| EPI_ISL_1469836 | Diretoria de Vigilância em Saúde | Epiclin | Fernando Hayashi Sant'Anna et al |
| EPI_ISL_760222,EPI_ISL_850184,EPI_ISL_850415,EPI_ISL_995771,EPI_ISL_1138964,EPI_ISL_1138902,EPI_ISL_1165052,EPI_ISL_1252433,EPI_ISL_1252434,EPI_ISL_1315343,EPI_ISL_1315379,EPI_ISL_1490064,EPI_ISL_1647348,EPI_ISL_1647349,EPI_ISL_1647350,EPI_ISL_1647351,EPI_ISL_1647352,EPI_ISL_1675277,EPI_ISL_1934722,EPI_ISL_1934823,EPI_ISL_1936604 | Division of Emerging Infectious Diseases, Bureau of Infectious Diseases Diagnosis Control, Korea Disease Control and Prevention Agency | Division of Emerging Infectious Diseases, Bureau of Infectious Diseases Diagnosis Control, Korea Disease Control and Prevention Agency | Ae Kyung Park et al |
| EPI_ISL_1591413,EPI_ISL_1608102 | Division of Medical Virology, National Health Laboratory Service (NHLS), Tygerberg Hospital / Stellenbosch University | Division of Medical Virology, Stellenbosch University and NHLS Tygerberg Hospital | Susan Engelbrecht et al |
| EPI_ISL_426163,EPI_ISL_425118,EPI_ISL_498044,EPI_ISL_506992,EPI_ISL_510599,EPI_ISL_515031,EPI_ISL_522534 | Division of Viral Diseases, Center for Laboratory Control of Infectious Diseases, Korea Centers for Diseases Control and Prevention | Division of Viral Diseases, Center for Laboratory Control of Infectious Diseases, Korea Centers for Diseases Control and Prevention | Jeong-Min Kim et al |
| EPI_ISL_1337507 | DOHMH Corona | New York City Public Health Laboratory | Jade Wang et al |
| EPI_ISL_1167047,EPI_ISL_1167061,EPI_ISL_1167054,EPI_ISL_1167082 | Dr. Andrija Stampar Teaching Institute of Public Health, Department of Clinical Microbiology | Istituto di Genomica Applicata | Jasmina Vranes et al |
| EPI_ISL_481746,EPI_ISL_481751 | Dr. Georges-L.-Dumont University Hospital Centre | National Microbiology Laboratory | Anna Majer et al |

| | | | |
|---|---|---|---|
| EPI_ISL_583803,EPI_ISL_583827 | Dr. Gernot Walder GmbH | Bergthaler laboratory, CeMM Research Center for Molecular Medicine of the Austrian Academy of Sciences | Alexandra Popa et al |
| EPI_ISL_2106197 | Dr. Gernot Walder GmbH | Dr. Gernot Walder GmbH | Sissy T. Lamprecht-Sonnleitner et al |
| EPI_ISL_1055386,EPI_ISL_1055408,EPI_ISL_1055422,EPI_ISL_1055437,EPI_ISL_1055447 | Dr. Leonard A. Miller Centre for Health Services | National Microbiology Laboratory (NML) | Anna Majer et al |
| EPI_ISL_417211,EPI_ISL_417212 | Dunedin Hospital | University of Otago | M.E. Quiñones-Mateu et al |
| EPI_ISL_461372,EPI_ISL_523464,EPI_ISL_1310926,EPI_ISL_1311220,EPI_ISL_2154204 | Dutch COVID-19 response team | Erasmus Medical Center | Bas Oude Munnink et al |
| EPI_ISL_523056 | Dutch COVID-19 response team | Erasmus Medical Center | OH consortium et al |
| EPI_ISL_1120207 | Dutch COVID-19 response team | Medical Microbiology, Maastricht University Medical Centre | Jozef Dingemans* et al |

| | | | |
|---|---|---|---|
| EPI_ISL_547447,EPI_ISL_547449,EPI_ISL_547451,EPI_ISL_547450,EPI_ISL_547452,EPI_ISL_547453,EPI_ISL_636513,EPI_ISL_636516,EPI_ISL_636517,EPI_ISL_636514,EPI_ISL_636518,EPI_ISL_636519,EPI_ISL_636515,EPI_ISL_636520,EPI_ISL_1014187,EPI_ISL_1014432,EPI_ISL_1014552,EPI_ISL_1014349,EPI_ISL_1014557,EPI_ISL_1014562,EPI_ISL_1014565,EPI_ISL_1014566,EPI_ISL_1014567,EPI_ISL_1014572,EPI_ISL_1014574,EPI_ISL_1014575,EPI_ISL_1014576,EPI_ISL_1014577,EPI_ISL_1014578,EPI_ISL_1014584,EPI_ISL_1014595,EPI_ISL_1014646,EPI_ISL_1014657,EPI_ISL_1014671,EPI_ISL_1014661,EPI_ISL_1059911,EPI_ISL_1089775,EPI_ISL_1089997,EPI_ISL_1090036,EPI_ISL_1165598,EPI_ISL_1165516,EPI_ISL_1232313,EPI_ISL_1232268,EPI_ISL_1232248,EPI_ISL_1232260,EPI_ISL_1232328,EP | Dutch COVID-19 response team | National Institute for Public Health and the Environment (RIVM) | Adam Meijer et al |
| EPI_ISL_1312683,EPI_ISL_1613792 | E. Gulbja laboratorija | Latvian Biomedical Research and Study Centre | Janis Pjalkovskis et al |
| EPI_ISL_501829,EPI_ISL_639690 | E. Gulbja Laboratorija | Latvian Biomedical Research and Study Centre | Ivars Silamiķelis et al |
| EPI_ISL_477161,EPI_ISL_478672 | Egyptian National Cancer Institute (ENCI) | Egyptian National Cancer Institute (ENCI) | Zekri et al |

| | | | |
|---|---|---|---|
| EPI_ISL_1483794 | El Camino Hospital | Santa Clara County Public Health Laboratory | Santa Clara County Public Health Department et al |
| EPI_ISL_1398364,EPI_ISL_1398925 | Emam Ali Hospital | Razi Vaccine and Serum Research Institute | Amir Kaffashi et al |
| EPI_ISL_1582978 | E.S.E. HOSPITAL SAN JOSE DE MAICAO | Instituto Nacional de Salud- Dirección de Investigación en Salud Pública | Katherine Laiton-Donato et al |
| EPI_ISL_1899281,EPI_ISL_1899485,EPI_ISL_1897646,EPI_ISL_1898814 | Ethiopian Biotechnology Institute (EBTI) | International Centre for Genetic Engineering and Biotechnology (ICGEB) and ARGO Open Lab for Genome Sequencing | Molalegne Bitew et al |
| EPI_ISL_875628,EPI_ISL_979481 | Eurofins Diatherix | Hudsonalpha Genome Sequencing Center | Jane Grimwood et al |
| EPI_ISL_1571307,EPI_ISL_1643945,EPI_ISL_1845854,EPI_ISL_1846153,EPI_ISL_1846077,EPI_ISL_1977952,EPI_ISL_2110737,EPI_ISL_2110489,EPI_ISL_2110488,EPI_ISL_2110499,EPI_ISL_2110532, EPI_ISL_2110453 | Eurofins LifeCodexx GmbH | Robert Koch Institute | ? |
| EPI_ISL_848582,EPI_ISL_848606 | Evandro Chagas Institute | Evandro Chagas Institute | Santos et al |
| EPI_ISL_1815257 | EXCITE Lab | Andersen lab at Scripps Research | Nicole L Washington et al |
| EPI_ISL_548971 | Expo2020 Emergency Center | Agiomix | Walaa Allam et al |
| EPI_ISL_1273102 | Faculty of Medicine, Al-Quds University | Faculty of Medicine, Al-Quds University | Ereqat et al |
| EPI_ISL_640050 | False Bay Hospital wc FBH | NHLS/UCT | Arash Iranzadeh et al |
| EPI_ISL_2038893 | Federal Budget Health Care Institution "Center of Hygiene and Epidemiology in Tver region" | Group of Genomics and Postgenomic Technologies of Central Research Institute of Epidemiology | Samoilov AE et al |
| EPI_ISL_2153095,EPI_ISL_2153110 | Fimlab Laboratories | Fimlab Laboratories | Minna Paloniemi et al |

| | | | |
|---|---|---|---|
| EPI_ISL_419560 | FL Bureau of Public Health Laboratories-Tampa | Pathogen Discovery, Respiratory Viruses Branch, Division of Viral Diseases, Centers for Disease Control and Prevention | Anna Uehara et al |
| EPI_ISL_489721,EPI_ISL_526569,EPI_ISL_653321 | Florida Bureau of Public Health Laboratories | Florida Bureau of Public Health Laboratories | Sarah Schmedes et al |
| EPI_ISL_476139 | Folkhalsomyndigheten | The Public Health Agency of Sweden | Oskar Karlsson Lindsjo et al |
| EPI_ISL_581487,EPI_ISL_581491,EPI_ISL_581492,EPI_ISL_581493 | Fondation Congolaise pour la recherche medicale (FCRM) | NGS Competence Center Tübingen, Institut für Medizinische Mikrobiologie und Hygiene, Universitätsklinikum Tübingen | Angel Angelov et al |
| EPI_ISL_1671923,EPI_ISL_1671925,EPI_ISL_1671926 | Fondation Congolaise pour la recherche medicale (FCRM), Francine Ntoumi | Institute of Tropical Medicine | Prof. Dr. Thirumalaisamy P. Velavan et al |
| EPI_ISL_1654214,EPI_ISL_1854763,EPI_ISL_1854768,EPI_ISL_1857284,EPI_ISL_1857282 | Fondation Congolaise pour la recherche medicale (FCRM), Francine Ntoumi | Institute of Tropical Medicine | Prof. Francine Ntoumi et al |
| EPI_ISL_912353,EPI_ISL_912358,EPI_ISL_912363,EPI_ISL_912364 | Fondation Congolaise pour la recherche medicale (FCRM), Francine Ntoumi | NGS Competence Center Tuebingen, Institut für Medizinische Mikrobiologie und Hygiene, Universitaetsklinikum Tübingen | Angel Angelov et al |
| EPI_ISL_2125018 | Fraunhofer-Institut fur Zelltherapie und Immunologie IZI AG Next-Generation Diagnostics | Robert Koch Institute | ? |
| EPI_ISL_431780 | Fujian Center for Disease Control and Prevention | Fujian Center for Disease Control and Prevention | Lin Qi et al |

| | | | |
|---|---|---|---|
| EPI_ISL_1613245,EPI_ISL_1667321,EPI_ISL_1667332,EPI_ISL_1992696,EPI_ISL_2134065 | Fulgent Genetics | Centers for Disease Control and Prevention Division of Viral Diseases, Pathogen Discovery | Dakota Howard et al |
| EPI_ISL_1182549,EPI_ISL_1182611 | Fundação Ezequiel Dias (FUNED) | Coordenação Geral de Laboratórios de Saúde Pública (CGLAB/DAEVS/SVS/MS) | Vagner Fonseca et al |
| EPI_ISL_493370,EPI_ISL_493372 | Furst Medical Laboratory | Norwegian Institute of Public Health, Department of Virology | Kathrine Stene-Johansen et al |
| EPI_ISL_1225321 | GA Department of Public Health | GA Department of Public Health | Stacy Reeves et al |
| EPI_ISL_730569,EPI_ISL_730572,EPI_ISL_730574,EPI_ISL_984742 | Gazi University Faculty of Medicine, Medical Virology Laboratory | Gazi University Faculty of Medicine, Medical Virology Laboratory | Erdem Şahin et al |
| EPI_ISL_2157199 | Gencore - Universidad de los Andes | Gencore - Universidad de los Andes | Marcela Guevara et al |
| EPI_ISL_1185245,EPI_ISL_1184582 | Genelabs Medical (Pvt) Ltd | Genelabs Medical (Pvt) Ltd | Chandanamali Punchihewa et al |
| EPI_ISL_2029122 | General Hospital "Abdulah Nakas" | Alea Genetic Centre | Rijad Konjhodzic et al |
| EPI_ISL_406798 | General Hospital of Central Theater Command of People's Liberation Army of China | BGI & Institute of Microbiology, Chinese Academy of Sciences & Shandong First Medical University & Shandong Academy of Medical Sciences & General Hospital of Central Theater Command of People's Liberation Army of China | Weijun Chen et al |
| EPI_ISL_1142575 | General Hospital - Ohrid | Research Center for Genetic Engineering and Biotechnology "Georgi D. Efremov” , Macedonian Academy of Sciences and Arts | Aleksandar J. Dimovski et al |

| | | | |
|---|---|---|---|
| EPI_ISL_677726,EPI_ISL_677722 | General Hospital - Ohrid | Research Center for Genetic Engineering and Biotechnology "Georgi D. Efremov” , Macedonian Academy of Sciences and Arts | RCGEB - MASA et al |
| EPI_ISL_677674 | General Hospital - Prilep | Research Center for Genetic Engineering and Biotechnology "Georgi D. Efremov” , Macedonian Academy of Sciences and Arts | RCGEB - MASA et al |
| EPI_ISL_2001028 | General Hospital - Tetovo | Research Center for Genetic Engineering and Biotechnology "Georgi D. Efremov” , Macedonian Academy of Sciences and Arts | Aleksandar J. Dimovski et al |

| | | | |
|---|---|---|---|
| EPI_ISL_746483,EPI_ISL_746489,EPI_ISL_746491,EPI_ISL_746511,EPI_ISL_746528,EPI_ISL_746586,EPI_ISL_746603,EPI_ISL_746616,EPI_ISL_746632,EPI_ISL_746691,EPI_ISL_746695,EPI_ISL_746708,EPI_ISL_746714,EPI_ISL_746763,EPI_ISL_746795,EPI_ISL_1167669,EPI_ISL_1167674,EPI_ISL_1167710,EPI_ISL_1167723,EPI_ISL_1167773,EPI_ISL_1167796,EPI_ISL_1167821,EPI_ISL_1167823,EPI_ISL_1167847,EPI_ISL_1167861,EPI_ISL_1167902,EPI_ISL_1167907,EPI_ISL_1167740,EPI_ISL_1167924,EPI_ISL_1300492,EPI_ISL_1300502,EPI_ISL_1321466,EPI_ISL_1321494,EPI_ISL_1321511,EPI_ISL_1321533,EPI_ISL_1321538,EPI_ISL_1321555,EPI_ISL_1470484,EPI_ISL_1470456,EPI_ISL_1470529,EPI_ISL_1633408,EPI_ISL_1633498,EPI_ISL_1633477,EPI_ISL_2009228,EPI_ISL_2009630 | Genetica Molecular and Subdepartamento de Virologia ISP Chile | Instituto de Salud Publica de Chile | Javier Tognarelli et al |
| EPI_ISL_936385,EPI_ISL_936386,EPI_ISL_936383,EPI_ISL_936384 | Genetica y Virologia, Facultad de Ciencias | Genetica y Virologia, Facultad de Ciencias | Panzera et al |
| EPI_ISL_2153105 | GENETICS DEPARTMENT, ZHEEN INTERNATIONAL HOSPITAL | GENETICS DEPARTMENT, ZHEEN INTERNATIONAL HOSPITAL | Khailany et al |

| | | | |
|---|---|---|---|
| EPI_ISL_2080269 | Genome Analysis Center, Kamma Memorial Hospital | Genome Analysis Center, Kamma Memorial Hospital | Hanako Yazawa et al |
| EPI_ISL_2036272 | Genome Centre | Genome Centre | Shovon Lal Sarkar et al |
| EPI_ISL_730221 | Genomica Lab Molecular, M√©xico | Andersen lab at Scripps Research | SEARCH Alliance San Diego with Jonathan Gonzalez Garcia et al |
| EPI_ISL_735350 | Genomic Laboratory (GLAB) (Conjoint lab of Health Directorate of Istanbul and Istanbul Technical University) | Genomic Laboratory (GLAB), Istanbul Technical University | Ilker Karacan et al |
| EPI_ISL_632908 | Genomic Sciences, Rehman Medical Institute | Genomic Sciences, Rehman Medical Institute | Ali et al |
| EPI_ISL_812847,EPI_ISL_812852,EPI_ISL_812790 | Genomics Program, Children Cancer Hospital | Genomics Program, Children Cancer Hospital | Hatem et al |
| EPI_ISL_1008758 | genXone SA, Molecular Diagnostics Laboratory / NZOZ | genXone SA, Research & Development Laboratory | Maciej Sykulski et al |
| EPI_ISL_745492 | Ginkgo Bioworks Clinical Laboratory | Utah Public Health Laboratory | Erin L. Young et al |
| EPI_ISL_1677798 | GMERS Medical Collage, GMERS Medical College,Civil Hospital, Sola, Ahmedabad | Gujarat Biotechnology Research Centre | Nitin Savaliya et al |
| EPI_ISL_775217,EPI_ISL_775019 | Gonoshasthya-RNA Molecular Research Center | Gonoshasthya-RNA Molecular Research Center | Nihad Adnan et al |
| EPI_ISL_1001457 | Gorgas memorial Institute For Health Studies | Gorgas memorial Institute For Health Studies | Díaz Y et al |
| EPI_ISL_496609,EPI_ISL_496719,EPI_ISL_496843 | Gorgas Memorial Laboratory of Health Studies | Gorgas Memorial Laboratory of Health Studies | Danilo Franco et al |
| EPI_ISL_1502844,EPI_ISL_1502957,EPI_ISL_1502995,EPI_ISL_1503069,EPI_ISL_1503137 | Gorgas Memorial Laboratory of Health Studies | Gorgas Memorial Laboratory of Health Studies | Gonzalez Claudia et al |

| | | | |
|---|---|---|---|
| EPI_ISL_1225330,EPI_ISL_1225342,EPI_ISL_1225429,EPI_ISL_1225449,EPI_ISL_1225452,EPI_ISL_1225468,EPI_ISL_1225481,EPI_ISL_1225489,EPI_ISL_1225501,EPI_ISL_1225506,EPI_ISL_1225513,EPI_ISL_1225529,EPI_ISL_1225539,EPI_ISL_1225542 | Gorgas Memorial Laboratory of Health Studies | Gorgas Memorial Laboratory of Health Studies | Yamilka Diaz et al |
| EPI_ISL_508250 | Government Medical College | National Institute of Biomedical Genomics | Arindam Maitra et al |
| EPI_ISL_455021 | Government Medical College, Vadodara | Gujarat Biotechnology Research Centre | Ankit Hinsu et al |
| EPI_ISL_2105484,EPI_ISL_2155974 | Governor Celestino Gallares Memorial Hospital | Philippine Genome Center | Francis A. Tablizo et al |
| EPI_ISL_794818 | Greek Genome Center, Biomedical Research Foundation of the Academy of Athens (BRFAA) | Greek Genome Center, Biomedical Research Foundation of the Academy of Athens (BRFAA) | Emmanouil Athanasiadis et al |
| EPI_ISL_640130,EPI_ISL_1040768,EPI_ISL_1706553 | Groote Schuur Hospital wc GSH | NHLS/UCT | Arash Iranzadeh et al |
| EPI_ISL_1273049,EPI_ISL_1273061,EPI_ISL_1273050,EPI_ISL_1273053,EPI_ISL_1273071,EPI_ISL_1273070,EPI_ISL_1273073 | Guam Public Health Laboratory | Centers for Disease Control and Prevention Division of Viral Diseases, Pathogen Discovery | Krista Queen et al |
| EPI_ISL_1798906,EPI_ISL_1818073,EPI_ISL_1823589,EPI_ISL_1823588,EPI_ISL_1823583 | Guam Public Health Laboratory | Centers for Disease Control and Prevention Division of Viral Diseases, Pathogen Discovery | Mili Sheth et al |
| EPI_ISL_413888 | Guangdong Provincial Institution of Public Health, Guangdong Provinical Center for Disease Control and Prevention | Guangdong Provincial Institution of Public Health | Jing Lu et al |

| | | | |
|---|---|---|---|
| EPI_ISL_509710 | Guatemala Ministry of Public Health | Pathogen Discovery, Respiratory Viruses Branch, Division of Viral Diseases, Centers for Disease Control and Prevention | Jing Zhang et al |
| EPI_ISL_509695 | Guatemala Ministry of Public Health | Pathogen Discovery, Respiratory Viruses Branch, Division of Viral Diseases, Centers for Disease Control and Prevention | Ying Tao et al |
| EPI_ISL_700503,EPI_ISL_700576 | Guguletu CHC wc GDH | NHLS/UCT | Arash Iranzadeh et al |
| EPI_ISL_426161,EPI_ISL_547657 | Gundersen Molecular Diagnostics Laboratory | Kabara Cancer Research Institute | Craig S. Richmond et al |
| EPI_ISL_2096775,EPI_ISL_2096776,EPI_ISL_2096891 | Hadassah Medical Center Clinical Virology Laboratory, Hadassah Ein Kerem | Hadassah Hebrew University Viral Sequencing Group, Hadassah Hebrew University Medical Center | Hadar Golan Berman et al |
| EPI_ISL_1268834 | Hannover Medical School, Institute of Virology | Hannover Medical School, Institute of Virology | Lars Steinbrück et al |
| EPI_ISL_668392,EPI_ISL_1058051 | Haukeland University Hospital, Dept. of Microbiology | Norwegian Institute of Public Health, Department of Virology | Kathrine Stene-Johansen et al |
| EPI_ISL_796014 | Hebei Provincial Center for Disease Control and Prevention, Shijiazhuang, Hebei Province; National Institute for Viral Disease Control and Prevention, China CDC | Hebei Provincial Center for Disease Control and Prevention, Shijiazhuang, Hebei Province; National Institute for Viral Disease Control and Prevention, | Shunxiang Qi et al |
| EPI_ISL_1500961,EPI_ISL_2029113 | HEGP - Laboratoire de Virologie | HEGP - Laboratoire de Virologie | David Veyer et al |

| | | | |
|---|---|---|---|
| EPI_ISL_1512225,EPI_ISL_1581188,EPI_ISL_1592421,EPI_ISL_1679967,EPI_ISL_1796460,EPI_ISL_1804291,EPI_ISL_1907237,EPI_ISL_2010581,EPI_ISL_2144594 | Helix/Illumina | Centers for Disease Control and Prevention Division of Viral Diseases, Pathogen Discovery | Dakota Howard et al |
| EPI_ISL_1340428 | Helix/Illumina | Centers for Disease Control and Prevention Division of Viral Diseases, Pathogen Discovery | Peter W. Cook et al |
| EPI_ISL_966962,EPI_ISL_978698 | Helix/Illumina | Respiratory Viruses Branch, Division of Viral Diseases, Centers for Disease Control and Prevention | Peter W. Cook et al |
| EPI_ISL_733027,EPI_ISL_1372346,EPI_ISL_1400527 | HELIX LLC | WHO National Influenza Centre Russian Federation | Andrey Komissarov et al |
| EPI_ISL_487381,EPI_ISL_501233,EPI_ISL_501235 | Hellenic Pasteur Institute, National Influenza Reference laboratory of Southern Greece & Unit of Bioinformatics and Applied Genomics | Hellenic Pasteur Institute, National Influenza Reference laboratory of Southern Greece & Unit of Bioinformatics and Applied Genomics | Vasiliki Pogka et al |
| EPI_ISL_428235 | Hematology Laboratory, Section of Molecular Diagnostics, University Clinical Centre, Medical University of Gdansk | Department of Virology, Faculty of Medicine, University of Helsinki, Helsinki, Finland | Marlena Robakowska et al |
| EPI_ISL_906752 | Hematology Laboratory, Section of Molecular Diagnostics, University Clinical Centre, Medical University of Gdansk | Laboratory of Recombinant Vaccines | Lukasz Rabalski et al |
| EPI_ISL_699979,EPI_ISL_700307 | Hematopathology Laboratory, ACTREC, TMC | Hematopathology Laboratory, ACTREC, TMC | Hematopathology Laboratory et al |
| EPI_ISL_955255 | HGSMF 26 CABO SAN LUCAS | BIOBANCO / COCTI | Borja-Aburto VH et al |

| | | | |
|---|---|---|---|
| EPI_ISL_845796 | Histopath | NSW Health Pathology - Institute of Clinical Pathology and Medical Research; Westmead Hospital; University of Sydney | CIDM-PH et al. |
| EPI_ISL_1170958 | HIV Molecular Laboratory, Ethiopian Public Health Institute, Ethiopia | HIV molecular lab, Ethiopian Public Health Institute, Ethiopian | Weldemariam et al |
| EPI_ISL_770032 | Hle-Asociacion De Atencion Integral Del Anciano San Cayetano | Inciensa, Instituto Costarricense de Investigación y Enseñanza en Nutrición y Salud | Francisco Duarte et al |
| EPI_ISL_476804 | Hong Kong Department of Health | School of Public Health, The University of Hong Kong | Dominic N.C. Tsang et al |
| EPI_ISL_964916,EPI_ISL_1110933 | Hopital | National Reference Center for Viruses of Respiratory Infections, Institut Pasteur, Paris | Marion Barbet et al |
| EPI_ISL_940534 | Hôpital Bichat Claude Bernard, Laboratoire de Virologie | IAME UMR1137 Inserm, Université de Paris, Hôpital Bichat | Antoine Bridier-Nahmias et al |
| EPI_ISL_710541,EPI_ISL_710575 | Hôpital Fattouma-Bourguiba de Monastir | Laboratoire des Procédés de Criblage Moléculaire et Cellulaire-Centre de Biotechnologie de Sfax | Souissi et al |
| EPI_ISL_961666 | Hôpital Georges L. Dumont | National Microbiology Laboratory (NML) | Anna Majer et al |
| EPI_ISL_1336649 | HOPITAL PRINCESSE GRACE | CNR Virus des Infections Respiratoires - France SUD | Antonin Bal et al |
| EPI_ISL_414631 | Hôpital Robert Debré Laboratoire de Virologie | National Reference Center for Viruses of Respiratory Infections, Institut Pasteur, Paris | Mélnie Albert et al |
| EPI_ISL_1904989,EPI_ISL_2142689 | HOPITAL SAINT ANDRE | CNR Virus des Infections Respiratoires - France SUD | Antonin Bal et al |

| | | | |
|---|---|---|---|
| EPI_ISL_2029931 | Hospital | National Reference Center for Viruses of Respiratory Infections, Institut Pasteur, Paris | Marion Barbet et al |
| EPI_ISL_1629762 | Hospital Carlos Alberto Seguín Escobedo - EsSalud | Laboratorio de Genómica Microbiana, Universidad Peruana Cayetano Heredia | Lenin Maturrano et al |
| EPI_ISL_1917450 | Hospital Center Luxembourg | Laboratoire national de sante, Microbiology, Microbial Genomics Platform | Anke Wienecke-Baldacchino et al |
| EPI_ISL_476384 | Hospital da Clínicas da Faculdade de Medicina da Universidade de São Paulo | Instituto de Medicina Tropical da Univesidade de São Paulo | Samples: Ingra Morales Claro et al |
| EPI_ISL_527750 | Hospital De Niños Dr. Carlos Saenz Herrera [San Jose/San Jose] | Inciensa, Instituto Costarricense de Investigación y Enseñanza en Nutrición y Salud | Francisco Duarte et al |
| EPI_ISL_1494944 | HOSPITAL DEPARTAMENTAL SAN VICENTE DE PAUL | Instituto Nacional de Salud- Dirección de Investigación en Salud Pública | Katherine Laiton-Donato et al |
| EPI_ISL_1469795 | Hospital Dia e Pronto Atendimento | Epiclin | Fernando Hayashi Sant'Anna et al |
| EPI_ISL_1201438 | HOSPITAL DR. FERNANDO ESCALANTE PRADILLA | Inciensa, Instituto Costarricense de Investigación y Enseñanza en Nutrición y Salud | Francisco Duarte et al |
| EPI_ISL_770028 | Hospital Dr. Raul Blanco Cervantes | Inciensa, Instituto Costarricense de Investigación y Enseñanza en Nutrición y Salud | Francisco Duarte et al |
| EPI_ISL_861657 | Hospital e Maternidade Sino Brasileiro | Instituto Adolfo Lutz, Interdiciplinary Procedures Center, Strategic Laboratory | Claudio Tavares Sacchi et al |

| | | | |
|---|---|---|---|
| EPI_ISL_1084623 | Hospital for Infectious Diseases, Molecular Diagnostics Laboratory, Warsaw, Poland | 28. Laboratory of Recombinant Vaccines, Intercollegiate Faculty of Biotechnology University of Gdansk and Medical University of Gdansk, 2. ViroGenetics - BSL3 Laboratory of Virology, Małopolska Centre of Biotechnology, Jagiellonian University | Lukasz Rabalski et al |
| EPI_ISL_1084629 | Hospital for Infectious Diseases, Molecular Diagnostics Laboratory, Warsaw, Poland | 31. Laboratory of Recombinant Vaccines, Intercollegiate Faculty of Biotechnology University of Gdansk and Medical University of Gdansk, 2. ViroGenetics - BSL3 Laboratory of Virology, Małopolska Centre of Biotechnology, Jagiellonian University | Lukasz Rabalski et al |
| EPI_ISL_424731 | Hospital General Regional No.66, Ciudad Juárez, Chihuahua. | Laboratorio Central de Epidemiología-DLVIE / Laboratorio de Secuenciación-Centro de Instrumentos. Instituto Mexicano del Seguro Social | Muñoz-Medina JE et al |
| EPI_ISL_654277 | Hospital General Universitario Gregorio Marañón | SeqCOVID-SPAIN consortium/IBV(CSIC) | Darío García de Viedma et al |
| EPI_ISL_1527021 | Hospital Guápiles | Inciensa, Instituto Costarricense de Investigación y Enseñanza en Nutrición y Salud | Pérez-Corrales C et al |
| EPI_ISL_481245,EPI_ISL_481247 | Hospital IESS Babahoyo | Institute of Microbiology, Universidad San Francisco de Quito | Belén Prado-Vivar et al |

| | | | |
|---|---|---|---|
| EPI_ISL_413016 | Hospital Israelita Albert Einstein | Instituto Adolfo Lutz, Interdisciplinary Procedures Center, Strategic Laboratory | Jaqueline Goes de Jesus et al |
| EPI_ISL_769992 | Hospital Metropolitano | Inciensa, Instituto Costarricense de Investigación y Enseñanza en Nutrición y Salud | Francisco Duarte et al |
| EPI_ISL_1067614 | HOSPITAL MEXICO | Inciensa, Instituto Costarricense de Investigación y Enseñanza en Nutrición y Salud | Francisco Duarte et al |
| EPI_ISL_539496 | Hospital Nostra Senyora de Meritxell | Instituto de Salud Carlos III | Iglesias-Caballero et al |
| EPI_ISL_1511405 | Hospital of the University of Pennsylvania Molecular Pathology Lab | Bushman Lab - University of Pennsylvania | John Everett et al |
| EPI_ISL_471270 | Hospital Oncológico Solca Núcleo de Quito | Institute of Microbiology, Universidad San Francisco de Quito | Sully Márquez et al |
| EPI_ISL_1503129 | Hospital Regional Anita Moreno Los Santos | Gorgas Memorial Laboratory of Health Studies | Gonzalez Claudia et al |
| EPI_ISL_1502822,EPI_ISL_1502837 | Hospital Regional Dr. Luis "Chicho" Fábrega | Gorgas Memorial Laboratory of Health Studies | Gonzalez Claudia et al |
| EPI_ISL_593774 | HOSPITAL REGIONAL LAMBAYEQUE | GENOMA MAYOR | Franklin R. Aguilar-Gamboa et al |
| EPI_ISL_682273,EPI_ISL_2103373 | HOSPITAL SAN JUAN DE DIOS | Inciensa, Instituto Costarricense de Investigación y Enseñanza en Nutrición y Salud | Francisco Duarte et al |
| EPI_ISL_491448 | Hospital San Rafael de Alajuela | Inciensa, Instituto Costarricense de Investigación y Enseñanza en Nutrición y Salud | Francisco Duarte et al |

| | | | |
|---|---|---|---|
| EPI_ISL_1972570,EPI_ISL_2001050 | Hospital Sharp | Microbial Genomics Laboratory | Bruno Gomez-Gil et al |
| EPI_ISL_1908862 | Hospital Universitari Arnau de Vilanova | Hospital Universitari Vall d'Hebron - Vall d'Hebron Institut de Recerca | Cristina Andrés et al |
| EPI_ISL_1063792 | Hospital Universitari Joan XXIII de Tarragona | Hospital Universitari Vall d'Hebron - Vall d'Hebron Institut de Recerca | Cristina Andrés et al |
| EPI_ISL_1017708 | Hospital Universitario Hernando Moncaleano Perdomo | Instituto Nacional de Salud- Dirección de Investigación en Salud Pública | Katherine Laiton-Donato et al |
| EPI_ISL_831086 | Hospital Universitario La Paz (Madrid) | SeqCOVID-SPAIN consortium/IBV(CSIC) | María Rodríguez-Tejedor et al |
| EPI_ISL_2134878,EPI_ISL_2134892,EPI_ISL_2134882,EPI_ISL_2134901,EPI_ISL_2134933 | HOSPITAL UNIVERSITARIO SON ESPASES | HOSPITAL UNIVERSITARIO SON ESPASES | Carla López-Causapé et al |
| EPI_ISL_2047772,EPI_ISL_2047773,EPI_ISL_2162276 | Hospital Universitari Vall d'Hebron - Vall d'Hebron Institut de Recerca | Hospital Universitari Vall d'Hebron - Vall d'Hebron Institut de Recerca | Cristina Andrés et al |
| EPI_ISL_693645,EPI_ISL_693651 | Hospital Vila Franca de Xira | Instituto Nacional de Saude (INSA) | Borges et al |
| EPI_ISL_2105676 | HOSP MUN DE MOGI DAS CRUZES PREF WALDEMAR COSTA FILHO | Instituto Butantan / Mendelics | Instituto Butantan: Dimas Tadeu Covas et al |
| EPI_ISL_787376,EPI_ISL_1077143 | Houston Methodist Hospital | Houston Methodist Hospital | S. Wesley Long et al |
| EPI_ISL_645109,EPI_ISL_645040,EPI_ISL_645009 | Human Genome Variation Research Group, Malopolska Centre of Biotechnology | Human Genome Variation Research Group, Malopolska Centre of Biotechnology | Kowalski et al |
| EPI_ISL_526225 | Hungarian Defence Forces Military Medical Centre | National Laboratory of Virology, Szentágothai Research Centre | Endre Gábor Tóth et al |
| EPI_ISL_412971 | HUS Diagnostiikkakeskus, Hallinto | Department of Virology Faculty of Medicine, Medicum University of Helsinki | Teemu Smura et al |

| | | | |
|---|---|---|---|
| EPI_ISL_1381056 | IAL Regional de Santo Andre | Instituto Adolfo Lutz, Interdiciplinary Procedures Center, Strategic Laboratory | Claudio Tavares Sacchi et al |
| EPI_ISL_1303530 | IAL Regional de São Jose do Rio Preto | Instituto Adolfo Lutz, Interdiciplinary Procedures Center, Strategic Laboratory | Claudio Tavares Sacchi et al |
| EPI_ISL_648000 | IA State Hygienic Laboratory | Pathogen Discovery, Respiratory Viruses Branch, Division of Viral Diseases, Centers for Disease Control and Prevention | Ying Tao et al |

| | | | |
|---|---|---|---|
| EPI_ISL_1703901,EPI_ISL_1703971,EPI_ISL_1704351,EPI_ISL_1704352,EPI_ISL_1704353,EPI_ISL_1704354,EPI_ISL_1704358,EPI_ISL_1704360,EPI_ISL_1704371,EPI_ISL_1704414,EPI_ISL_1704443,EPI_ISL_1704466,EPI_ISL_1704467,EPI_ISL_1704515,EPI_ISL_1704517,EPI_ISL_1704520,EPI_ISL_1704527,EPI_ISL_1704535,EPI_ISL_1704541,EPI_ISL_1704543,EPI_ISL_1704549,EPI_ISL_1704596,EPI_ISL_1704617,EPI_ISL_1704620,EPI_ISL_1704623,EPI_ISL_1704627,EPI_ISL_1704635,EPI_ISL_1704638,EPI_ISL_1841238,EPI_ISL_1841236,EPI_ISL_1841306,EPI_ISL_1841329,EPI_ISL_1841253,EPI_ISL_1841379,EPI_ISL_1841356,EPI_ISL_1928407,EPI_ISL_1928408,EPI_ISL_1928411,EPI_ISL_1928427,EPI_ISL_1928430,EPI_ISL_1928432,EPI_ISL_1928444,EPI_ISL_1928459,EPI_ISL_1928462,EPI_ISL_1928481,EPI_IS | ICMR-National Institute of Virology - INSACOG | NIV Influenza | Dr. Varsha Potdar et al |
| EPI_ISL_751618 | ID Bureau of Laboratories | Genomics and Discovery, Respiratory Viruses Branch, Division of Viral Diseases, Centers for Disease Control and Prevention | Krista Queen et al |

| | | | |
|---|---|---|---|
| EPI_ISL_848232,EPI_ISL_1323228,EPI_ISL_1494247,EPI_ISL_1494521,EPI_ISL_2080003 | Illinois Department of Public Health | Gagnon Lab, Southern Illinois University | Keith Gagnon et al |
| EPI_ISL_2109475,EPI_ISL_2109472 | IMD - Institut fur Medizinische Diagnostik Berlin-Potsdam | Robert Koch Institute | ? |
| EPI_ISL_1722241 | IMD - MVZ Labor Martinsried | Robert Koch Institute | ? |
| EPI_ISL_1372093,EPI_ISL_1662451,EPI_ISL_1663562 | Immunogenomics lab, Institute of Life Sciences, Bhubaneswar | Institute of Life Sciences - INSACOG | Sunil K. Raghav et al |
| EPI_ISL_1018099,EPI_ISL_1018094,EPI_ISL_1018079,EPI_ISL_1018086,EPI_ISL_2001092,EPI_ISL_2001075 | Immunology, Noguchi Memorial Institute for Medical Research | Immunology, Noguchi Memorial Institute for Medical Research | Adu et al |
| EPI_ISL_2001066,EPI_ISL_2001065 | Immunology, Noguchi Memorial Institute for Medical Research | Immunology, Noguchi Memorial Institute for Medical Research | Campbell et al |
| EPI_ISL_1213358 | IMT-UFRN/RN | Bioinformatics Laboratory / LNCC | Alessandra P Lamarca et al |
| EPI_ISL_648218 | INBIRS-UBA | Laboratorio Mixto de Biotecnología Acuática (LMBA) | Joaquín Ezpeleta et al |
| EPI_ISL_1517393,EPI_ISL_1517399,EPI_ISL_1517401,EPI_ISL_1517402,EPI_ISL_1517403,EPI_ISL_1517405,EPI_ISL_1517421,EPI_ISL_1517433 | Inciensa, Instituto Costarricense de Investigación y Enseñanza en Nutrición y Salud | Inciensa, Instituto Costarricense de Investigación y Enseñanza en Nutrición y Salud | Cristian Peréz-Corrales et al |

| | | | |
|---|---|---|---|
| EPI_ISL_476704,EPI_ISL_476702 | Incubadora Venezolana de Ciencia, Venezuela | Incubadora Venezolana de Ciencia, Venezuela / Instituto Nacional de Salud, Bogotá, Colombia / Grupo de Investigaciones Microbiológicas-UR (GIMUR), Departamento de Biología, Facultad de Ciencias Naturales, Universidad del Rosario, Bogotá, Colombia / Icahn School of Medicine at Mount Sinai, New York, USA | Alberto Paniz-Mondolfi et al |
| EPI_ISL_413523 | Indian Council of Medical Research-National Institute of Virology | National Influenza Center, Indian Council of Medical Research-National Institute of Virology | Potdar V et al |
| EPI_ISL_2032642 | Infectious Diseases Hospital № 2 | Group of Genomics and Postgenomic Technologies of Central Research Institute of Epidemiology | Samoilov AE et al |
| EPI_ISL_2151337 | Infectious Diseases, King Faisal Hospital Research Center | Infectious Diseases, King Faisal Hospital Research Center | Alhamlan F et al |
| EPI_ISL_884386,EPI_ISL_884409,EPI_ISL_884411 | Infectious Diseases, Quest Diagnostics | Infectious Diseases, Quest Diagnostics | Rosenthal et al |
| EPI_ISL_496372 | Infectolab | Andersen lab at Scripps Research | SEARCH Alliance San Diego with Samuel Navarro Alvarez et al |
| EPI_ISL_1692611 | Infinity Biologix | Centers for Disease Control and Prevention Division of Viral Diseases, Pathogen Discovery | Dakota Howard et al |

| | | | |
|---|---|---|---|
| EPI_ISL_1576833,EPI_ISL_1576834,EPI_ISL_1577798,EPI_ISL_1577817,EPI_ISL_1577815,EPI_ISL_1577816 | INHRR | Laboratorio de Virología Molecular | Loureiro CL et al |
| EPI_ISL_2000618 | INMI Lazzaro Spallanzani IRCCS | INMI Lazzaro Spallanzani IRCCS | M Rueca et al |
| EPI_ISL_410546 | INMI Lazzaro Spallanzani IRCCS | Laboratory of Virology, INMI Lazzaro Spallanzani IRCCS | Maria R. Capobianchi et al |
| EPI_ISL_2151101 | INSACOG-Sikkim | National Institute of Biomedical Genomics – INSACOG | Arindam Maitra et al |
| EPI_ISL_1357700 | INSACOG-WB | National Institute of Biomedical Genomics | Arindam Maitra et al |
| EPI_ISL_1589885,EPI_ISL_1589927 | INSACOG-WB | National Institute of Biomedical Genomics – INSACOG | Arindam Maitra et al |
| EPI_ISL_940781 | INSPI-CRN de Influenza y otros virus respiratorios | INSPI-Centro de Investigación Multidisciplinaria de la DTIDI | Leandro Patiño et al |
| EPI_ISL_826811,EPI_ISL_826814,EPI_ISL_826820,EPI_ISL_826827,EPI_ISL_826832,EPI_ISL_826836 | INSPI-CRN DE INFLUENZA Y OTROS VIRUS RESPIRATORIOS | Instituto de Salud Publica de Chile | Javier Tognarelli et al |
| EPI_ISL_955159,EPI_ISL_955165 | Institute for Biocides and Medical Ecology | Institute of microbiology and Immunology, Faculty of Medicine, University of Belgrade | Knezevic et al |
| EPI_ISL_1633467,EPI_ISL_1827950 | Institute for Health Research, Epidemiological Surveillance and Training (IRESSEF) | Abbott | Souleymane Mboup et al |

| | | | |
|---|---|---|---|
| EPI_ISL_459956,EPI_ISL_718277,EPI_ISL_718290,EPI_ISL_718282,EPI_ISL_728157,EPI_ISL_944099,EPI_ISL_962526,EPI_ISL_1055264,EPI_ISL_1406185,EPI_ISL_1406194,EPI_ISL_1406250,EPI_ISL_1673683,EPI_ISL_1787254,EPI_ISL_2090886,EPI_ISL_2090887 | Institute for Medical Research, Infectious Disease Research Centre, National Institutes of Health, Ministry of Health Malaysia | Institute for Medical Research, Infectious Disease Research Centre, National Institutes of Health, Ministry of Health Malaysia | Suppiah J et al |
| EPI_ISL_430441 | Institute for Medical Research, Infectious Disease Research Centre, National Institutes of Health, Ministry of Health Malaysia | Institute for Medical Research, Infectious Disease Research Centre, National Institutes of Health, Ministry of Health Malaysia | Suppiah.J et al |
| EPI_ISL_455791 | Institute for Medical Research, Infectious Disease Research Centre, National Institutes of Health, Ministry of Health Malaysia | Malaysia Genome Institute | Mohd Noor Mat Isa et al |
| EPI_ISL_1491570,EPI_ISL_1511133 | Institute for Urban Disease Control and Prevention | COVID-19 Network Investigations (CONI) Alliance | Elizabeth Batty et al |
| EPI_ISL_707707,EPI_ISL_733546,EPI_ISL_733564,EPI_ISL_812932 | Institute for Urban Disease Control and Prevention | COVID-19 Network Investigations (CONI) Alliance | Kamolthip Atsawawaranunt et al |
| EPI_ISL_1061425,EPI_ISL_1061431 | Institute of Biocides and Medical Ecology, Belgarde, Serbia | Virology Department Institute of Microbiology and Immunology Faculty of Medicine University of Belgrade | Banko Ana et al |
| EPI_ISL_909737 | Institute of Biocides and Medical Ecology, Belgrade, Serbia | Virology Department, Institute of microbiology and immunology, Faculty of Medicine University of Blegrade | Banko Ana et al |

| | | | |
|---|---|---|---|
| EPI_ISL_1938477 | Institute of Epidemiology, Disease Control and Research (IEDCR) | Institute for Developing Science and Health Initiatives (ideSHi) | Hassan Afrad et al |
| EPI_ISL_1181824,EPI_ISL_1181826,EPI_ISL_1510968,EPI_ISL_1511055,EPI_ISL_1914765,EPI_ISL_1964394 | Institute of Microbiology and Immunology, Faculty of Medicine, University of Ljubljana | Institute of Microbiology and Immunology, Faculty of Medicine, University of Ljubljana | Alen Suljič et al |
| EPI_ISL_1098758 | Institute of Microbiology and Immunology, Faculty of Medicine, University of Ljubljana | Institute of Microbiology and Immunology, Faculty of Medicine, University of Ljubljana | Samo Zakotnik et al |
| EPI_ISL_635272,EPI_ISL_635263 | Institute of Microbiology and Immunology, Faculty of Medicine, University of Ljubljana | Institute of Microbiology and Immunology, Faculty of Medicine, University of Ljubljana | Tomaž Mark Zorec et al |
| EPI_ISL_539788 | Institute of Microbiology, Universidad San Francisco de Quito | Institute of Microbiology, Universidad San Francisco de Quito | Andrea Macias et al |
| EPI_ISL_486842,EPI_ISL_486843,EPI_ISL_486844,EPI_ISL_491940,EPI_ISL_527810,EPI_ISL_824292 | Institute of Microbiology, Universidad San Francisco de Quito | Institute of Microbiology, Universidad San Francisco de Quito | Belén Prado-Vivar et al |
| EPI_ISL_516648 | Institute of Microbiology, Universidad San Francisco de Quito | Institute of Microbiology, Universidad San Francisco de Quito | Juan José Guadalupe et al |
| EPI_ISL_516650,EPI_ISL_516652 | Institute of Microbiology, Universidad San Francisco de Quito | Institute of Microbiology, Universidad San Francisco de Quito | Prado-Vivar et al |
| EPI_ISL_417482,EPI_ISL_660530,EPI_ISL_660532,EPI_ISL_660534,EPI_ISL_728202,EPI_ISL_1896686,EPI_ISL_2086703,EPI_ISL_2100441 | Institute of Microbiology, Universidad San Francisco de Quito | Institute of Microbiology, Universidad San Francisco de Quito | Sully Márquez et al |
| EPI_ISL_1443648,EPI_ISL_1443652,EPI_ISL_1443657,EPI_ISL_1443662 | Institute of Microbiology, Universidad San Francisco de Quito | Omics Sciences Laboratory | Derly Andrade Molina et al |

| | | | |
|---|---|---|---|
| EPI_ISL_577740 | Institute of Virology, Biomedical Research Center of the Slovak Academy of Sciences, Bratislava | Faculty of Natural Sciences, Comenius University, Bratislava | Broňa Brejová et al |
| EPI_ISL_1805024 | Institute of Virology, Medical Center, University of Freiburg, Freiburg, Germany | Institute of Virology, Clinial Virus Genomics, Medical Center, University of Freiburg, Freiburg, Germany | Jonas Fuchs et al |
| EPI_ISL_1654812 | Institute of Virology, Vaccines and Sera "Torlak" | Institute of microbiology and Immunology, Faculty of Medicine, University of Belgrade | Knezevic et al |
| EPI_ISL_1138693 | Institut für Medizinische Virologie, Universitätsklinikum Frankfurt | Institut für Medizinische Virologie, Universitätsklinikum Frankfurt | Barbara Muehlemann et al |
| EPI_ISL_475922 | Institut für Virologie am Department für Hygiene, Mikrobiologie und Public Health | Bergthaler laboratory, CeMM Research Center for Molecular Medicine of the Austrian Academy of Sciences | Alexandra Popa et al |
| EPI_ISL_508688,EPI_ISL_508696 | Institut für Virologie und Epidemiologie der Viruskrankheiten, Universitätsklinikum Tübingen | NGS Competence Center Tübingen, Institut für Medizinische Mikrobiologie und Hygiene, Universitätsklinikum Tübingen | Angel Angelov et al |
| EPI_ISL_1416192,EPI_ISL_1415425,EPI_ISL_1443001 | Institut National d'hygiène | Unité Mixte Internationale TransVIHMI (UMI 233 IRD – U1175 INSERM - Université de Montpellier) IRD (Institut de recherche pour le développement) | Mounerou SALOU et al |

| | | | |
|---|---|---|---|
| EPI_ISL_1508958 | Institut National d'hygiène | Unité Mixte Internationale TransVIHMI (UMI 233 IRD – U1175 INSERM - Université de Montpellier) IRD (Institut de recherche pour le développement) | Mounerou SALOU et al |
| EPI_ISL_1913082 | Institut National d'Hygiène | Laboratoire de Biotechnologie | Mouna Ouadghiri et al |
| EPI_ISL_1404615,EPI_ISL_1436817,EPI_ISL_1434443,EPI_ISL_1435763,EPI_ISL_1437749 | Institut National d'Hygiène | Unité Mixte Internationale TransVIHMI (UMI 233 IRD – U1175 INSERM - Université de Montpellier) IRD (Institut de recherche pour le développement) | Mounerou SALOU et al |
| EPI_ISL_1406177 | Institut National d’Hygiène (INH) | Unité Mixte Internationale TransVIHMI (UMI 233 IRD – U1175 INSERM - Université de Montpellier)IRD (Institut de recherche pour le développement) | Mounerou SALOU et al |
| EPI_ISL_875544 | Instituto de Biotecnologia - UNESP-Botucatu-SP | Instituto de Biotecnologia - UNESP-Botucatu-SP | Leila Sabrina Ullmann et al |
| EPI_ISL_872097 | Instituto de Diagnostico y Referencia Epidemiologicos (INDRE) | Instituto de Diagnostico y Referencia Epidemiologicos (INDRE) | Abril Rodriguez-Maldonado et al |
| EPI_ISL_576273,EPI_ISL_576277,EPI_ISL_658901 | Instituto de Diagnostico y Referencia Epidemiologicos (INDRE) | Instituto de Diagnostico y Referencia Epidemiologicos (INDRE) | Ernesto Ramirez-Gonzalez et al |
| EPI_ISL_516620,EPI_ISL_576275 | Instituto de Diagnostico y Referencia Epidemiologicos (INDRE) | Instituto de Diagnostico y Referencia Epidemiologicos (INDRE) | Gisela Barrera-Badillo et al |

| | | | |
|---|---|---|---|
| EPI_ISL_455434 | Instituto de Diagnostico y Referencia Epidemiologicos (INDRE) | Instituto de Diagnostico y Referencia Epidemiologicos (INDRE) | Taboada Ramírez Blanca. Ramirez-Gonzalez Ernesto et al |
| EPI_ISL_1301458,EPI_ISL_1301512,EPI_ISL_1301514,EPI_ISL_1301519,EPI_ISL_1301685,EPI_ISL_1301691,EPI_ISL_1301696,EPI_ISL_1301703,EPI_ISL_1301709 | Instituto de Diagnostico y Referencia Epidemiologicos InDRE_RNLSP | Instituto de Biotecnología de la UNAM | Authors from IBT et al |
| EPI_ISL_1060683,EPI_ISL_1060708,EPI_ISL_1060704,EPI_ISL_1060720,EPI_ISL_1060733 | Instituto de Diagnostico y Referencia Epidemiologicos (INDRE)_RNLSP | Instituto de Diagnostico y Referencia Epidemiologicos (INDRE) | Claudia Wong-Arambula et al |
| EPI_ISL_913914,EPI_ISL_913924,EPI_ISL_913927,EPI_ISL_913934,EPI_ISL_913938,EPI_ISL_913948,EPI_ISL_913959,EPI_ISL_933669,EPI_ISL_1054948,EPI_ISL_1054962,EPI_ISL_1054970,EPI_ISL_1054978,EPI_ISL_1054986,EPI_ISL_1168464,EPI_ISL_1168489,EPI_ISL_1168521,EPI_ISL_1168540,EPI_ISL_1168542,EPI_ISL_1168598,EPI_ISL_1168607,EPI_ISL_1168613,EPI_ISL_1168615,EPI_ISL_1168621,EPI_ISL_1168624 | Instituto de Diagnostico y Referencia Epidemiologicos INDRE_RNLSP | Instituto de Diagnostico y Referencia Epidemiologicos (INDRE) | Claudia Wong-Arambula et al |
| EPI_ISL_748139,EPI_ISL_748138 | Instituto de Investigaciones Biológicas Clemente Estable | Institut Pasteur de Montevideo | Daiana Mir et al |

| | | | |
|---|---|---|---|
| EPI_ISL_1395785 | Instituto de Investigaciones Biomédicas en Retrovirus y SIDA (INBIRS) | Área de Secuenciación del Laboratorio de Virología del Hospital de Niños Dr. Ricardo Gutierrez on behalf of 'Proyecto Argentino Interinstitucional de genomica de SARS-CoV-2' (PAIS Consortium) | Vanesa Seery et al |
| EPI_ISL_1629797,EPI_ISL_1629795 | Instituto de Medicina Tropical Alexander Von Humboldt, Universidad Peruana Cayetano Heredia | Laboratorio de Genómica Microbiana, Universidad Peruana Cayetano Heredia | Lenin Maturrano et al |
| EPI_ISL_1378844 | Instituto de Medicina Tropical & Salud Global Universidad Iberoamericana | Grubaugh Lab - Yale School of Public Health | Joseph Fauver et al |
| EPI_ISL_491188,EPI_ISL_491191 | Instituto Gulbenkian de Ciência | Instituto Gulbenkian de Ciência | João Costa et al |
| EPI_ISL_491270,EPI_ISL_491297,EPI_ISL_491288 | Instituto Gulbenkian de Ciência | Instituto Gulbenkian de Ciência | Susana Ladeiro et al |
| EPI_ISL_412972 | Instituto Nacional de Enfermedades Respiratorias | Instituto de Diagnostico y Referencia Epidemiologicos (INDRE) | Ramirez-Gonzalez Ernesto et al |
| EPI_ISL_837781,EPI_ISL_837811 | Instituto Nacional de Enfermedades Respiratorias (INER) | Instituto Nacional de Enfermedades Respiratorias (INER) | Celia Boukadida et al |
| EPI_ISL_1347897,EPI_ISL_1347920,EPI_ISL_1347933,EPI_ISL_1347934,EPI_ISL_1347935,EPI_ISL_1545346,EPI_ISL_1545358 | Instituto Nacional de Investigação em Saúde | KRISP, KZN Research Innovation and Sequencing Platform | Morais J et al |
| EPI_ISL_491945,EPI_ISL_491946,EPI_ISL_491951,EPI_ISL_491953,EPI_ISL_491954 | Instituto Nacional de Investigación en Salud Pública - INSPI | INSPI - Charité | Alfredo Bruno Caicedo et al |
| EPI_ISL_2080661 | Instituto Nacional de Medicina Genomica | Instituto Nacional de Medicina Genomica | Hidalgo-Miranda A et al |

| | | | |
|---|---|---|---|
| EPI_ISL_522986 | Instituto Nacional de Medicina Genómica | Instituto Nacional de Medicina Genómica | Hidalgo-Miranda A et al |
| EPI_ISL_536486,EPI_ISL_536497,EPI_ISL_536518,EPI_ISL_536525 | Instituto Nacional de Salud | Laboratorio de Infecciones Respiratorias Agudas | Eduardo Juscamayta Lopez et al |
| EPI_ISL_941943,EPI_ISL_941955 | Instituto Nacional de Salud, Bogotá, Colombia | Centro de Investigaciones en Microbiología y Biotecnología-UR (CIMBIUR), Facultad de Ciencias Naturales, Universidad del Rosario, Bogotá, Colombia Instituto Nacional de Salud, Bogotá, Colombia Icahn School of Medicine at Mount Sinai, New York, USA | Luz Helena Patiño et al |
| EPI_ISL_447756,EPI_ISL_447794,EPI_ISL_447805 | Instituto Nacional de Salud, Bogotá, Colombia | Grupo de Investigaciones Microbiológicas-UR (GIMUR), Departamento de Biología, Facultad de Ciencias Naturales, Universidad del Rosario, Bogotá, Colombia Instituto Nacional de Salud, Bogotá, Colombia Icahn School of Medicine at Mount Sinai, New York, USA | Juan David Ramírez et al |

| | | | |
|---|---|---|---|
| EPI_ISL_498160,EPI_ISL_498167,EPI_ISL_526969,EPI_ISL_526967,EPI_ISL_526949,EPI_ISL_653745,EPI_ISL_653747,EPI_ISL_653754,EPI_ISL_653762,EPI_ISL_739663,EPI_ISL_739672,EPI_ISL_739673,EPI_ISL_739676,EPI_ISL_739677,EPI_ISL_739680 | Instituto Nacional de Salud, Bogotá, Colombia | Instituto Nacional de Salud, Bogotá, Colombia | Katherine Laiton-Donato et al |
| EPI_ISL_1424059 | Instituto Nacional de Salud- Dirección de Investigación en Salud Pública | Instituto Nacional de Salud- Dirección de Investigación en Salud Pública | Katherine Laiton-Donato et al |
| EPI_ISL_956289,EPI_ISL_956291,EPI_ISL_956294 | Instituto Nacional de Salud- Dirección de Redes de Laboratorios de Salud Pública | Instituto Nacional de Salud- Dirección de Investigación en Salud Pública | Katherine Laiton-Donato et al |
| EPI_ISL_456122,EPI_ISL_456140 | Instituto Nacional de Salud - Unidad de Secuenciación y Análisis Genómico | Instituto Nacional de Salud, Universidad Cooperativa de Colombia, Instituto Alexander von Humboldt, Imperial College-London, London School of Hygiene & Tropical Medicine | Katherine Laiton-Donato et al |
| EPI_ISL_510889,EPI_ISL_511532,EPI_ISL_511691,EPI_ISL_693553,EPI_ISL_1023463 | Instituto Nacional de Saude (INSA) | Instituto Nacional de Saude (INSA) | Borges et al |
| EPI_ISL_1854273 | Instituto Nacional de Saude (INSA) and BioSystems & Integrative Sciences Institute (BioISI) Genomics Unit, FCUL | Instituto Nacional de Saude (INSA) and BioSystems & Integrative Sciences Institute (BioISI) Genomics Unit, FCUL | Borges et al |

| | | | |
|---|---|---|---|
| EPI_ISL_2004296 | Instituto Nacional de Saude (INSA) and i3S - Instituto de Investigação e Inovação em Saúde | Instituto Nacional de Saude (INSA) | Borges et al |
| EPI_ISL_732227,EPI_ISL_732255,EPI_ISL_1854270 | Instituto Nacional de Saude (INSA) and Instituto Gulbenkian de Ciencia (IGC) | Instituto Nacional de Saude (INSA) and Instituto Gulbenkian de Ciencia (IGC) | Borges et al |
| EPI_ISL_887420,EPI_ISL_887421,EPI_ISL_887458,EPI_ISL_887464,EPI_ISL_887500,EPI_ISL_887502,EPI_ISL_964947 | Instituto Nacional de Saude (INS), Mozambique | KRISP, KZN Research Innovation and Sequencing Platform | Nalia Ismael et al |
| EPI_ISL_418206,EPI_ISL_481243 | Institut Pasteur Dakar | Institut Pasteur de Dakar | Ndongo Dia et al |
| EPI_ISL_498238,EPI_ISL_498239 | Institut Pasteur de Dakar | Institut Pasteur de Dakar | Ndongo Dia et al |
| EPI_ISL_999032 | Institut Pasteur de Guinée | Institut Pasteur de Dakar | Grayo Solene et al |
| EPI_ISL_613433,EPI_ISL_613430,EPI_ISL_613420,EPI_ISL_613443,EPI_ISL_613446,EPI_ISL_613425 | Institut Pasteur de la Guadeloupe | Institut Pasteur de la Guadeloupe | Marion Barbet et al |
| EPI_ISL_1992100 | Institut Pasteur de Montevideo | Institut Pasteur de Montevideo | Daiana Mir et al |
| EPI_ISL_2110643 | Institut Pasteur du Maroc | Functional Genomics platform/CNRST | Melloul Marouane et al |
| EPI_ISL_459969,EPI_ISL_459974,EPI_ISL_459978,EPI_ISL_459983,EPI_ISL_459984 | Institut Pasteur du Maroc | Institut Pasteur du Maroc | Marion Barbet et al |
| EPI_ISL_1652055,EPI_ISL_1652057,EPI_ISL_1652056 | Integrated Biorepository of H3Africa Uganda – IBRH3AU | Molecular Biology Laboratory | Savannah Mwesigwa et al |
| EPI_ISL_1968643 | INTERLAB | Omics Sciences Laboratory | Derly Andrade Molina et al |

| | | | |
|---|---|---|---|
| EPI_ISL_1919648 | Ipoh Public Health Laboratory (MKAI), Ministry of Health Malaysia | Institute for Medical Research, Infectious Disease Research Centre, National Institutes of Health, Ministry of Health Malaysia | Suppiah J et al |
| EPI_ISL_751447 | IRCCS Sacro Cuore Don Calabria Hospital, Department of Infectious, Tropical Diseases & Microbiology | University of Verona, Department of Biotechnology | Antonio Mori et al |
| EPI_ISL_1630270 | IRESSEF | Abbott | Souleymane Mboup et al |
| EPI_ISL_1910386,EPI_ISL_1910388,EPI_ISL_1910391,EPI_ISL_1910390 | Iressef Genomics lab | IRESSEF | Souleymane MBOUP et al |
| EPI_ISL_1167135,EPI_ISL_1167139,EPI_ISL_1167164,EPI_ISL_1167126 | Iressef Genomics lab | L'institut de Recherche en Santé, de Surveillance Épidémiologique et de Formation (IRESSEF) | Souleymane MBOUP et al |
| EPI_ISL_516907,EPI_ISL_516913,EPI_ISL_575334,EPI_ISL_649062 | Israel Central Virology laboratory | Israel Central Virology laboratory | Neta Zuckerman et al |
| EPI_ISL_1073586,EPI_ISL_1209905,EPI_ISL_1240651,EPI_ISL_1763181,EPI_ISL_2084096,EPI_ISL_2084952,EPI_ISL_2085869,EPI_ISL_2083957 | Israel Central Virology laboratory | Israel National Consortium for SARS-CoV-2 sequencing | Neta Zuckerman et al |
| EPI_ISL_1240647,EPI_ISL_1240649 | Israel Central Virology Laboratory | Israel National Consortium for SARS-CoV-2 sequencing | Neta Zuckerman et al |

| | | | |
|---|---|---|---|
| EPI_ISL_1254965 | Istituto Zooprofilattico Sperimentale del Mezzogiorno | Telethon Institute of Genetics and Medicine (TIGEM) | Antonio Grimaldi Patrizia Annunziata Francesco Panariello Biancamaria Pierri Claudia Tiberio Valentina Bouche Chiara Colantuono Maria Concetta Cuomo Denise Di Concilio Lucio Di Filippo Anna Manfredi Marcello Salvi Antonio Limone Luigi Atripaldi Pellegrino Cerino Andrea Ballabio Davide Cacchiarelli et al |
| EPI_ISL_736840,EPI_ISL_837310,EPI_ISL_1424168 | Istituto Zooprofilattico Sperimentale del Mezzogiorno | TIGEM | Antonio Grimaldi et al |
| EPI_ISL_2004073,EPI_ISL_2004075,EPI_ISL_2004069 | IU-Cerrahpasa, Cerrahpasa School of Medicine, COVID-19 Lab | IU-Cerrahpasa, Cerrahpasa School of Medicine, COVID-19 Lab | Mert Kuskucu et al |
| EPI_ISL_422424 | Jaber Al Ahmad Al Sabah Hospital | Dasman diabetes Institute | Fahd Al-Mulla et al |
| EPI_ISL_2151355,EPI_ISL_2151365 | Jamil-ur-Rahman Center for Genome Research, Dr. Panjwani Center for Molecular Medicine and Drug Research | Jamil-ur-Rahman Center for Genome Research, Dr. Panjwani Center for Molecular Medicine and Drug Research | Irfan et al |
| EPI_ISL_779290,EPI_ISL_779283 | Jamil-ur-Rahman Center for Genome Research, Dr. Panjwani Center for Molecular Medicine and Drug Research | Jamil-ur-Rahman Center for Genome Research, Dr. Panjwani Center for Molecular Medicine and Drug Research | Shakeel et al |
| EPI_ISL_416575 | Japanese Quarantine Stations | Pathogen Genomics Center, National Institute of Infectious Diseases | Tsuyoshi Sekizuka et al |
| EPI_ISL_1753676 | Jessa | Jessa | Cruys et al. on behalf of the Jessa_cmdLab et al |
| EPI_ISL_911881 | Johns Hopkins Hospital Department of Pathology | Johns Hopkins Hospital Department of Pathology | C. Paul Morris et al |

| | | | | | |
|---|---|---|---|---|---|
| Carlos Cortes et alEPI_ISL_953403 Laboratorio de Investigaciones de BaneySwiss Tropical and Public Health InstituteSalome Hosch et alEPI_ISL_1700675,EPI_ISL_1700676,EPI_ISL_1700685,EPI_ISL_1700687,EPI_ISL_2002669,EPI_ISL_2002670,EPI_ISL_2002674,EPI_ISL_2002676,EPI_ISL_2002686 Laboratorio de Investigaciones de BaneyUniversity Hospital Basel, Clinical BacteriologyCarlos Cortes et alEPI_ISL_648327,EPI_ISL_648334,EPI_ISL_648337,EPI_ISL_648339,EPI_ISL_648373,EPI_ISL_649157 Laboratorio de la Dirección de EpidemiologíaÁrea de Secuenciación del Laboratorio de Virología del Hospital de Niños Dr. Ricardo Gutierrez on | J.W. Ruby Memorial Hospital | WVU and Marshall University Combined Genomics Core Facilities | James Denvir et alEPI_ISL_1528686 J.W. Ruby Memorial HospitalWVU and Marshall University Combined Genomics Core FacilitiesJames Denvir et alEPI_ISL_1742526 Kafkas University, Faculty of Medicine, Department of Medical MicrobiologyKafkas University, Faculty of Medicine, Department of Medical MicrobiologyMurat Karamese et alEPI_ISL_495424 Kansas Health and Environmental LabKansas Health and Environmental LabMike Grose et alEPI_ISL_1167563,EPI_ISL_2007091 Kantonsspital Baden AGInstitute of Medical Virology, University of ZurichVerena Kufner et alEPI_ISL_1791065,EPI_ISL_1791066 Kantor Pengelola Sumbawa TechnoparkNational Institute of Health Research and DevelopmentSubangkit et alEPI_ISL_2047557 Karolinska UniversitetslaboratorietThe | Alexandr Shevtsov et al | EPI_ISL_435045,EPI_ISL_435046,EPI_ISL_435047 |
| EPI_ISL_1138520 | Laboratory of Communicable Diseases | 1. Laboratory of Communicable Diseases (Estonia); 2. Eurofins Genomics Europe Sequencing GmbH | Liidia Dotsenko et al | | |

| | | | |
|---|---|---|---|
| EPI_ISL_1716736 | Laboratory of Immunohematology, Division of Hematology | Greek Genome Center, Biomedical Research Foundation of the Academy of Athens (BRFAA) | Emmanouil Athanasiadis et al |
| EPI_ISL_434468,EPI_ISL_437890 | Laboratory of Microbiology, Medical School, National and Kapodistrian University of Athens | Laboratory of Biology, Department of Medicine, Democritus University of Thrace | Kassela K. et al |
| EPI_ISL_875997 | Laboratory of Molecular Biology, Diagnostyka sp. z o.o. | genXone SA, Research & Development Laboratory | Maciej Sykulski et al |
| EPI_ISL_801569,EPI_ISL_801589,EPI_ISL_801650,EPI_ISL_801722,EPI_ISL_801769 | Laboratory of Molecular Virology, Pontificia Universidad Católica de Chile | MSHS Pathogen Surveillance Program | Leonardo I. Almonacid et al |
| EPI_ISL_1181442,EPI_ISL_1181366 | Laboratory of Respiratory Viruses and Measles, Oswaldo Cruz Institute, FIOCRUZ | Laboratory of Respiratory Viruses and Measles, Oswaldo Cruz Institute, FIOCRUZ | Paola Resende et al |
| EPI_ISL_1495053 | Laboratory of Respiratory Viruses Teaching and Clinical Center of the Medical University of Lodz | Biobank Lab, Department of Molecular Biophysics, Faculty of Biology and Environmental Protection, University of Lodz | Strapagiel Dominik et al |
| EPI_ISL_945547 | Laboratory of Virology and Molecular Diagnostics | Institute of Public Health of Republic of North Macedonia Laboratory of Virology and Molecular Diagnostics | Maja Kuzmanovska et al |
| EPI_ISL_1669957 | Laboratory of virology and molecular diagnostics, Institute of Public Health | Laboratory of virology and molecular diagnostics, Institute of Public Health | Kuzmanovska M et al |
| EPI_ISL_1334570,EPI_ISL_1448020,EPI_ISL_1448018 | Laboratory of virology, National center of expertise | RSE "National Center for Biotechnology" and RSE "National Center of Expertise" | Shevtsov Alexandr et al |

| | | | |
|---|---|---|---|
| EPI_ISL_454599 | Laboratory of virology, National Center of Expertise | Laboratory of molecular-genetic research, National Center for Expertise, Kazakhstan National Center for Biotechnology, Kazakhstan | Abdaliyev Askar et al |
| EPI_ISL_1364612,EPI_ISL_1365697,EPI_ISL_1365742,EPI_ISL_1489929 | Laboratory of Virology, National center of expertise | RSE "National Center of Expertise" and RSE "National center for Biotechnology" | Abdaliyev Askar et al |
| EPI_ISL_576121,EPI_ISL_576120 | Laboratory, The Bio Arte Limited | Laboratory, The Bio Arte Limited | Biazzo et al |
| EPI_ISL_1904299 | Labor Doz DDr Stefan Mustafa | AGES IMED Vienna | Sara Meschini et al |
| EPI_ISL_1862442 | Labor Dr. Fenner und Kollegen | Heinrich Pette Institute, Leibniz Institute for Experimental Virology | Alexis Robitaille et al |
| EPI_ISL_1566762,EPI_ISL_1851414 | Labor Dr. Heidrich & Kollegen MVZ GmbH Hamburg | Robert Koch Institute | ? |
| EPI_ISL_1847409 | Labor Dr. Spranger | Robert Koch Institute | ? |
| EPI_ISL_1725188 | Labor Dr. Wisplinghoff - KÃ¶ln | Robert Koch Institute | ? |
| EPI_ISL_487429 | Labor Kneißler GmbH & Co. KG | Heinrich Pette Institute, Leibniz Institute for Experimental Virology | Thomas Günther et al |
| EPI_ISL_1233630,EPI_ISL_1233673 | Labormedizinisches Zentrum Dr Risch | University Hospital Basel, Clinical Bacteriology | Tim Roloff et al |
| EPI_ISL_1573247,EPI_ISL_1724314 | Labor Prof. Dr. G. Enders MVZ GbR | Robert Koch Institute | ? |
| EPI_ISL_1140257 | Labor ZOTZ\|KLIMAS; MVZ Düsseldorf-Centrum | Robert Koch Institute | ? |
| EPI_ISL_456157,EPI_ISL_456209,EPI_ISL_456389 | LabPLUS | Institute of Environmental Science and Research (ESR) | Matt Storey et al |

| | | | |
|---|---|---|---|
| EPI_ISL_1172031,EPI_ISL_1250708,EPI_ISL_1315322,EPI_ISL_1621304,EPI_ISL_1621305,EPI_ISL_1621296,EPI_ISL_1621306,EPI_ISL_1621315,EPI_ISL_1621319,EPI_ISL_1621314,EPI_ISL_1621307,EPI_ISL_1621324,EPI_ISL_1621325,EPI_ISL_1621323,EPI_ISL_1904852,EPI_ISL_1904857,EPI_ISL_1904859,EPI_ISL_1967902,EPI_ISL_1967901,EPI_ISL_2103201,EPI_ISL_2103202 | LabPLUS | Institute of Environmental Science and Research (ESR) | Rachel Boyle et al |
| EPI_ISL_548070,EPI_ISL_579105,EPI_ISL_579106,EPI_ISL_579116,EPI_ISL_579403,EPI_ISL_579425,EPI_ISL_622773,EPI_ISL_661258,EPI_ISL_877209,EPI_ISL_877212,EPI_ISL_877226,EPI_ISL_1016853,EPI_ISL_1016867 | LabPLUS | Institute of Environmental Science and Research (ESR) | Xiaoyun Ren et al |
| EPI_ISL_1969245 | Lab RSUP DR Mohammad Hoesin Palembang | National Institute of Health Research and Development | Subangkit et al |
| EPI_ISL_547986,EPI_ISL_547994,EPI_ISL_547995,EPI_ISL_548104,EPI_ISL_548105,EPI_ISL_579422,EPI_ISL_622778 | LabTests | Institute of Environmental Science and Research (ESR) | Xiaoyun Ren et al |
| EPI_ISL_1827216 | Lab voor klinische biologie | Lab voor klinische biologie | Marija Janevska et al |
| EPI_ISL_419265 | Lab voor klinische biologie | Onderzoeksgroep Virologie | Laurens Lambrechts et al |

| | | | |
|---|---|---|---|
| EPI_ISL_1628364 | LACEN do Estado de Goias | Instituto Adolfo Lutz, Interdiciplinary Procedures Center, Strategic Laboratory | Claudio Tavares Sacchi et al |
| EPI_ISL_943976 | LACEN do Estado de Tocantins | Instituto Adolfo Lutz, Interdiciplinary Procedures Center, Strategic Laboratory | Claudio Tavares Sacchi et al |
| EPI_ISL_1040835 | LACEN do Mato Grosso do Sul | Instituto Adolfo Lutz, Interdiciplinary Procedures Center, Strategic Laboratory | Claudio Tavares Sacchi et al |
| EPI_ISL_1213274 | LAFEM/UESC | Bioinformatics Laboratory / LNCC | Alessandra P Lamarca et al |
| EPI_ISL_1904967,EPI_ISL_1904969,EPI_ISL_1904971 | LAM ORIADE ABBAYE ST MARTIN D'HERES | CNR Virus des Infections Respiratoires - France SUD | Antonin Bal et al |
| EPI_ISL_2013037,EPI_ISL_2013040 | Lancet Laboratories | National Institute for Communicable Diseases of the National Health Laboratory Service | Amoako DG et al |
| EPI_ISL_738332,EPI_ISL_738336,EPI_ISL_812266 | Landstuhl Regional Medical Center | United States Air Force School of Aerospace Medicine | Anthony Fries et al |
| EPI_ISL_1941773 | LA SOURCE | Laboratory of genomics and metagenomics | Trestan Pillonel et al |
| EPI_ISL_884249 | LATE - Laboratório de Técnicas Especiais - Hospital Israelita Albert Einstein | LATE - Laboratório de Técnicas Especiais - Hospital Israelita Albert Einstein | Deyvid Amgarten et al |
| EPI_ISL_437096 | Latvijas Infektoloģijas centrs | Latvian Biomedical Research and Study Centre | Ivars Silamiķelis et al |
| EPI_ISL_1091784 | LDSP | Universidad Nacional de Colombia - Laboratorio Genómico One Health | Andres F. Cardona-Rios et al |
| EPI_ISL_2155048 | LDSP CALDAS | Instituto Nacional de Salud- Dirección de Investigación en Salud Pública | Katherine Laiton-Donato et al |

| | | | |
|---|---|---|---|
| EPI_ISL_1805632 | LDSP TOLIMA | Instituto Nacional de Salud- Dirección de Investigación en Salud Pública | Katherine Laiton-Donato et al |
| EPI_ISL_1582987 | LDSP VICHADA | Instituto Nacional de Salud- Dirección de Investigación en Salud Pública | Katherine Laiton-Donato et al |
| EPI_ISL_498552 | Lebanese American University | Lebanese American University | Abi Habib et al |
| EPI_ISL_447900 | Lednicky Laboratory at Emerging Pathogens Institute | University of Florida | Lednicky et al |
| EPI_ISL_1516777 | LESP Aguascalientes | Instituto de Diagnostico y Referencia Epidemiologicos (INDRE) | Claudia Wong-Arambula et al |
| EPI_ISL_1532244 | LESP Baja California Sur | Instituto de Diagnostico y Referencia Epidemiologicos (INDRE) | Claudia Wong-Arambula et al |
| EPI_ISL_1700816 | LESP Campeche | Instituto de Diagnostico y Referencia Epidemiologicos (INDRE) | Claudia Wong-Arambula et al |
| EPI_ISL_1494658,EPI_ISL_2158212,EPI_ISL_2158224 | LESP Guanajuato | Instituto de Diagnostico y Referencia Epidemiologicos (INDRE) | Claudia Wong-Arambula et al |
| EPI_ISL_1334381,EPI_ISL_2157329 | LESP Guerrero | Instituto de Diagnostico y Referencia Epidemiologicos (INDRE) | Claudia Wong-Arambula et al |
| EPI_ISL_1805491,EPI_ISL_2094505 | LESP Jalisco | Instituto de Diagnostico y Referencia Epidemiologicos (INDRE) | Claudia Wong-Arambula et al |
| EPI_ISL_1805495 | LESP Jalisco/Unidad de Patologia | Instituto de Diagnostico y Referencia Epidemiologicos (INDRE) | Claudia Wong-Arambula et al |

| | | | |
|---|---|---|---|
| EPI_ISL_1504072 | LESP Morelos | Instituto de Diagnostico y Referencia Epidemiologicos (INDRE) | Claudia Wong-Arambula et al |
| EPI_ISL_1504058 | LESP Nuevo Leon | Instituto de Diagnostico y Referencia Epidemiologicos (INDRE) | Claudia Wong-Arambula et al |
| EPI_ISL_1857255 | LESP Oaxaca | Instituto de Diagnostico y Referencia Epidemiologicos (INDRE) | Claudia Wong-Arambula et al |
| EPI_ISL_1424021 | LESP Queretaro | Instituto de Diagnostico y Referencia Epidemiologicos (INDRE) | Claudia Wong-Arambula et al |
| EPI_ISL_2157320 | LESP Quintana Roo | Instituto de Diagnostico y Referencia Epidemiologicos (INDRE) | Claudia Wong-Arambula et al |
| EPI_ISL_2094506,EPI_ISL_2094507,EPI_ISL_2094508 | LESP San Luis Potosi | Instituto de Diagnostico y Referencia Epidemiologicos (INDRE) | Claudia Wong-Arambula et al |
| EPI_ISL_1337389 | LESP Sinaloa | Instituto de Diagnostico y Referencia Epidemiologicos (INDRE) | Claudia Wong-Arambula et al |
| EPI_ISL_1821158 | LESP Tamaulipas | Instituto de Diagnostico y Referencia Epidemiologicos (INDRE) | Claudia Wong-Arambula et al |
| EPI_ISL_1219574,EPI_ISL_1613125 | LIC | Latvian Biomedical Research and Study Centre | Janis Pjalkovskis et al |

| | | | |
|---|---|---|---|
| EPI_ISL_834888,EPI_ISL_1486529,EPI_ISL_1484918,EPI_ISL_1759112,EPI_ISL_1806530,EPI_ISL_1806764,EPI_ISL_1829281,EPI_ISL_1829220,EPI_ISL_1830471,EPI_ISL_1857887,EPI_ISL_1912578,EPI_ISL_1986886,EPI_ISL_2091685,EPI_ISL_2091852,EPI_ISL_2117860,EPI_ISL_2119041,EPI_ISL_2138697 | Lighthouse Lab in Alderley Park | Wellcome Sanger Institute for the COVID-19 Genomics UK (COG-UK) Consortium | Jacquelyn Wynn et al |

| | | | |
|---|---|---|---|
| EPI_ISL_1246284,EPI_ISL_1246268,EPI_ISL_1264032,EPI_ISL_1294622,EPI_ISL_1316100,EPI_ISL_1330057,EPI_ISL_1326666,EPI_ISL_1327318,EPI_ISL_1327609,EPI_ISL_1327362,EPI_ISL_1344560,EPI_ISL_1374314,EPI_ISL_1376830,EPI_ISL_1409773,EPI_ISL_1454606,EPI_ISL_1454202,EPI_ISL_1473725,EPI_ISL_1473648,EPI_ISL_1484542,EPI_ISL_1505036,EPI_ISL_1504705,EPI_ISL_1504748,EPI_ISL_1519930,EPI_ISL_1534778,EPI_ISL_1537321,EPI_ISL_1534917,EPI_ISL_1583979,EPI_ISL_1595472,EPI_ISL_1615877,EPI_ISL_1631286,EPI_ISL_1632412,EPI_ISL_1632316,EPI_ISL_1653120,EPI_ISL_1697735,EPI_ISL_1697635,EPI_ISL_1698521,EPI_ISL_1719066,EPI_ISL_1719089,EPI_ISL_1719142,EPI_ISL_1831443,EPI_ISL_1831538 | Lighthouse Lab in Cambridge | Wellcome Sanger Institute for the COVID-19 Genomics UK (COG-UK) Consortium | Rob Howes et al |
| EPI_ISL_540349 | Lighthouse Lab in Glasgow | Wellcome Sanger Institute for the COVID-19 Genomics UK (COG-UK) consortium | Harper VanSteenhouse et al |

| | | | |
|---|---|---|---|
| EPI_ISL_1042374,EPI_ISL_1264130,EPI_ISL_1544580,EPI_ISL_1544515,EPI_ISL_1719573,EPI_ISL_1742109,EPI_ISL_2006127,EPI_ISL_2118067,EPI_ISL_2119748 | Lighthouse Lab in Glasgow | Wellcome Sanger Institute for the COVID-19 Genomics UK (COG-UK) Consortium | Harper VanSteenhouse et al |
| EPI_ISL_681636,EPI_ISL_1377698,EPI_ISL_1537056,EPI_ISL_1594669,EPI_ISL_1631645,EPI_ISL_1698093,EPI_ISL_1700157,EPI_ISL_1790496,EPI_ISL_1790928,EPI_ISL_1830656,EPI_ISL_1830403,EPI_ISL_1986243,EPI_ISL_1985465,EPI_ISL_2004751,EPI_ISL_2004702,EPI_ISL_2021832,EPI_ISL_2091582,EPI_ISL_2118947,EPI_ISL_2138658 | Lighthouse Lab in Milton Keynes | Wellcome Sanger Institute for the COVID-19 Genomics UK (COG-UK) Consortium | The Lighthouse Lab in Milton Keynes et al |
| EPI_ISL_2117856 | Limbach - MVZ Humangenetik Ulm | Robert Koch Institute | ? |
| EPI_ISL_1791067 | Limmattal Hospital | Institute of Medical Virology, University of Zurich | Verena Kufner et al |
| EPI_ISL_603119,EPI_ISL_636839,EPI_ISL_636846,EPI_ISL_636889 | Lithuanian University of Health Sciences Hospital, Department of Laboratory Medicine | Lithuanian University of Health Sciences, Molecular cardiology lab. | Lukas Zemaitis et al |
| EPI_ISL_526391,EPI_ISL_665129 | Liverpool Clinical Laboratories | COVID-19 Genomics UK (COG-UK) Consortium | Sam Haldenby et al |
| EPI_ISL_2142670 | LMS REUNILAB | CNR Virus des Infections Respiratoires - France SUD | Antonin Bal et al |
| EPI_ISL_469051,EPI_ISL_469054 | LNR National Reference Laboratory, Mohammed VI University of Health Sciences | Medical Biotechnology Laboratory, Rabat Medical and Pharmacy School, Mohammed The Vth University in Rabat | Meriem LAAMARTI et al |

| | | | |
|---|---|---|---|
| EPI_ISL_1827701,EPI_ISL_1827703 | Lobamba | National Institute for Communicable Diseases of the National Health Laboratory Service | Maphalala GP et al |
| EPI_ISL_451294,EPI_ISL_574337,EPI_ISL_574366,EPI_ISL_768481 | LSUHS Emerging Viral Threat Laboratory | Microbial Genome Sequencing Center | Jeremy P. Kamil et al |
| EPI_ISL_1936202,EPI_ISL_1936244,EPI_ISL_1936258,EPI_ISL_1936275,EPI_ISL_1936287 | Main Chemical Laboratories Egypt Army | Main Chemical Laboratories Egypt Army | Mohamed Seadawy et al |
| EPI_ISL_1936296,EPI_ISL_1936301,EPI_ISL_1936356 | Main Chemical Laboratories Egypt Army | Main Chemical Laboratories Egypt Army | Sherine Helmy et al |
| EPI_ISL_755307,EPI_ISL_755480 | Maine Health and Environmental Testing Laboratory | Tewhey Lab, The Jackson Laboratory | Matluk et al |
| EPI_ISL_807151 | Makmal Kesihatan Awam Kebangsaan, Kementerian Kesihatan Malaysia | Integrative Pharmacogenomics Institute (iPROMISE) | Mohd Zaki Salleh et al |
| EPI_ISL_582124,EPI_ISL_977593 | Malaysia Genome Institute | Malaysia Genome Institute | Mohd Noor Mat Isa et al |
| EPI_ISL_2156920 | Mapmygenome | CSIR-Centre for Cellular and Molecular Biology-INSACOG | Lamuk Zaveri et al |
| EPI_ISL_509464 | Maryland Department of Health | Maryland Department of Health | Keller et al |
| EPI_ISL_522854 | Maryland Department of Health | Maryland Department of Health | Maryland Department of Health Laboratories Administration et al |
| EPI_ISL_1910617 | Mashrek Medical Diagnostic Center | Microbial Pathogenomics Lab - LAU | Jad Koweyes et al |
| EPI_ISL_677023 | Masonic Medical Research Institute | Wadsworth Center, New York State Department.of Health | Nathan Tucker et al |
| EPI_ISL_460235,EPI_ISL_765803 | Massachusetts General Hospital | Infectious Disease Program, Broad Institute of Harvard and MIT | Lemieux et al |

| | | | |
|---|---|---|---|
| EPI_ISL_1752154,EPI_ISL_2006677,EPI_ISL_2095276 | Max von Pettenkofer Institute, Virology, National Reference Center for Retroviruses, LMU Munich | Laboratory for Functional Genome Analysis; Dept. Genomics; Gene Center of the LMU Munich | Max Muenchhoff et al |
| EPI_ISL_1196245 | Mayo Clinic & Mayo Clinic Laboratories | Minnesota Department of Health, Public Health Laboratory | Alexandra Lorentz et al |
| EPI_ISL_1258019,EPI_ISL_1258052,EPI_ISL_1594056 | MB-Cadham Provincial laboratory | National Microbiology Laboratory (NML) | Anna Majer et al |
| EPI_ISL_1847210,EPI_ISL_1848786 | MDI Limbach Berlin GmbH; MVZ Labor Berlin | Robert Koch Institute | ? |
| EPI_ISL_524892 | MD PHL | MD PHL | Maryland Department of Health Laboratories Administration et al |
| EPI_ISL_482772,EPI_ISL_1109484,EPI_ISL_1109628 | Medical Ain Shams Research Institute (MASRI), Ain Shams University | Medical Ain Shams Research Institute (MASRI), Ain Shams University | Hesham Elghazaly et al |
| EPI_ISL_1508843 | Medical Genetics Laboratory, Regional Centre of Medical Genetics, Emergency County Hospital Craiova | Medical Genetics Laboratory, Regional Centre of Medical Genetics, Emergency County Hospital Craiova | Anca-Lelia (Riza) Costache et al |
| EPI_ISL_1041001 | Medical Laboratory Bruss | Laboratory of Recombinant Vaccines | Lukasz Rabalski et al |
| EPI_ISL_2107524 | Medical Laboratory Sciences, Arab American University | Medical Laboratory Sciences, Arab American University | Al-Jawabreh et al |
| EPI_ISL_873164,EPI_ISL_873165 | Medical Laboratory Sciences, Arab American University | Medical Laboratory Sciences, Arab American University | Dumaidi et al |
| EPI_ISL_707692 | Medical Research Center, Faculty of Medicine, Syarif Hidayatullah State Islamic University Jakarta | Medical Research Center, Faculty of Medicine, Syarif Hidayatullah State Islamic University Jakarta | Chris Adhiyanto et al |
| EPI_ISL_2102111 | Medics Labor AG | Clinical Bacteriology | Tim Roloff et al |

| | | | |
|---|---|---|---|
| EPI_ISL_2129020,EPI_ISL_2129024,EPI_ISL_2129038 | Medizinische Laboratorien Dusseldorf | Robert Koch Institute | ? |
| EPI_ISL_456377,EPI_ISL_456378 | MedLab Central Ltd | Institute of Environmental Science and Research (ESR) | Matt Storey et al |
| EPI_ISL_2110379 | Med. Labor Prof. Schenk Dr. Ansorge & Kollegen | Robert Koch Institute | ? |
| EPI_ISL_451583 | Medlab Pathology | NSW Health Pathology - Institute of Clinical Pathology and Medical Research; Westmead Hospital; University of Sydney | CIDM-PH et al. |
| EPI_ISL_1791042 | Megalab, Molecular and Cytogenetics Diagnostics | Department for Virology, Molecular Biology and Genome Research, R. G. Lugar Center for Public Health Research, National Center for Disease Control and Public Health (NCDC) of Georgia. | Giorgi Tomashvili et al |
| EPI_ISL_568881,EPI_ISL_568910,EPI_ISL_569236,EPI_ISL_804503,EPI_ISL_896118,EPI_ISL_2036730 | MEPHI, Aix Marseille University | MEPHI, Aix Marseille University | Anthony LEVASSEUR et al |
| EPI_ISL_1663546,EPI_ISL_1663548,EPI_ISL_1663549 | MGM Medical College, Jamshedpur | Institute of Life Sciences - INSACOG | Sunil K. Raghav et al |
| EPI_ISL_529842,EPI_ISL_577565,EPI_ISL_1058856 | Michigan Department of Health and Human Services, Bureau of Laboratories | Michigan Department of Health and Human Services, Bureau of Laboratories | Blankenship HM et al |
| EPI_ISL_1069173 | Micrbiological Laboratory,Lu`an Center for Disease Control and Prevention | Micrbiological Laboratory,Lu`an Center for Disease Control and Prevention | Yang Wei et al |

| | | | |
|---|---|---|---|
| EPI_ISL_480331,EPI_ISL_480337,EPI_ISL_480341,EPI_ISL_480345 | Microbial Genomics Laboratory, Institut Pasteur de Montevideo | Microbial Genomics Laboratory, Institut Pasteur de Montevideo | Cecilia Salazar et al |
| EPI_ISL_419726 | Microbiological Diagnostic Unit Public Health Laboratory | Microbiological Diagnostic Unit Public Health Laboratory | Seemann T. et al |
| EPI_ISL_563530 | Microbiological Diagnostic Unit - Public Health Laboratory (MDU-PHL) | MDU-PHL | Seemann et al |
| EPI_ISL_480770,EPI_ISL_592447,EPI_ISL_593100,EPI_ISL_641177,EPI_ISL_812425,EPI_ISL_812428,EPI_ISL_854755,EPI_ISL_877571,EPI_ISL_979360,EPI_ISL_1033152,EPI_ISL_1055284,EPI_ISL_1249999,EPI_ISL_1250002,EPI_ISL_1250005,EPI_ISL_1913110 | Microbiological Diagnostic Unit - Public Health Laboratory (MDU-PHL) | MDU-PHL | Seemann T. et al |
| EPI_ISL_418267 | Microbiology and Immunology department, Pasteur institute in Ho Chi Minh city | Microbiology and Immunology department, Pasteur institute in Ho Chi Minh city | Nguyen et al |
| EPI_ISL_1181810 | Microbiology and Virology Unit,Azienda Ospedale Padova,Padova,Italy | Department of Molecular Medicine,Computational Medicine Group,Univeresity of Padova,Padova,Italy | Elisa Franchin et al |
| EPI_ISL_513925 | Microbiology & Bioinformatics and Biostatistics, Kohat University of Science and Technology (Pakistan) & Shanghai Jiao Tong University (China) | Microbiology & Bioinformatics and Biostatistics, Kohat University of Science and Technology (Pakistan) & Shanghai Jiao Tong University (China) | Khan et al |
| EPI_ISL_2086160 | Microbiology Department. Complexo Hospitalario Universitario de Vigo | Microbiology Department. Complexo Hospitalario Universitario de Vigo | Alfaya N et al |

| | | | |
|---|---|---|---|
| EPI_ISL_2136859,EPI_ISL_2136883,EPI_ISL_2140641 | Microbiology Department, Laboratori Clínic Metropolitana Nord. Hospital Universitari Germans Trias i Pujol | Can Ruti SARS-CoV-2 Sequencing Hub (HUGTiP/IrsiCaixa/IGTP) | Marc Noguera-Julian et al |
| EPI_ISL_1661929,EPI_ISL_1999791 | Microbiology Department, Laboratori Clínic Metropolitana Nord. Hospital Universitari Germans Trias i Pujol. | Can Ruti SARS-CoV-2 Sequencing Hub (HUGTiP/IrsiCaixa/IGTP) | Marc Noguera-Julian et al |
| EPI_ISL_848191 | Microbiology Lab | NBCC Sequencing Facility | Dr. Jeff Wrana et al |
| EPI_ISL_1633322 | Microbiology Lab, University Hospital ATTIKON | Central Public Health Lab, National Public Health Organization | Kyriaki Tryfinopoulou et al |
| EPI_ISL_981938 | Microbiology Service, Hospital Universitario Clinico San Cecilio, Granada | Microbiology Service, Hospital Universitario Clinico San Cecilio, Granada | Adolfo de Salazar et al |
| EPI_ISL_1820897 | Microvida | Microvida | Suzan D. Pas et al |
| EPI_ISL_456214 | Middlemore Hospital | Institute of Environmental Science and Research (ESR) | Matt Storey et al |
| EPI_ISL_1082258,EPI_ISL_1082267,EPI_ISL_1082259,EPI_ISL_1315319 | Middlemore Hospital | Institute of Environmental Science and Research (ESR) | Rachel Boyle et al |
| EPI_ISL_548002,EPI_ISL_548130,EPI_ISL_548145,EPI_ISL_579310,EPI_ISL_579312,EPI_ISL_622783,EPI_ISL_622787,EPI_ISL_622819,EPI_ISL_649122,EPI_ISL_755635,EPI_ISL_794626,EPI_ISL_1016845,EPI_ISL_1016874,EPI_ISL_1016872 | Middlemore Hospital | Institute of Environmental Science and Research (ESR) | Xiaoyun Ren et al |
| EPI_ISL_718136 | Ministry of Health Hospitals | Institute of Health and Community Medicine | David Perera et al |
| EPI_ISL_812769 | Ministry of Health Turkey | Ministry of Health Turkey | Fatma Bayrakdar et al |

| | | | |
|---|---|---|---|
| EPI_ISL_1712681,EPI_ISL_1712696,EPI_ISL_1712910,EPI_ISL_1712939,EPI_ISL_1713159,EPI_ISL_1713204,EPI_ISL_1713210,EPI_ISL_1713251,EPI_ISL_1713269,EPI_ISL_1713339,EPI_ISL_1713480,EPI_ISL_1713883,EPI_ISL_1713508,EPI_ISL_1713559,EPI_ISL_1713580,EPI_ISL_1713817 | Ministry of Public Health / Hamad Medical Corporation | Biomedical Research Center (BRC), Qatar University / Qatar Genome Project (QGP) | BRC: Fatiha M. Benslimane et al |
| EPI_ISL_1714179,EPI_ISL_1714185,EPI_ISL_1714195,EPI_ISL_1714210,EPI_ISL_1714284,EPI_ISL_1714309,EPI_ISL_1714340,EPI_ISL_1714343,EPI_ISL_1714380,EPI_ISL_1714383,EPI_ISL_1714404,EPI_ISL_1714420,EPI_ISL_1714444,EPI_ISL_1714477,EPI_ISL_1714521 | Ministry of Public Health / Hamad Medical Corporation | Weill Cornell Medical College - Qatar (WCM-Q), Genomics Core Laboratory / Qatar Genome Project (QGP) | WCMQ: Ayeda A. Ahmed et al |
| EPI_ISL_462909,EPI_ISL_527592 | Minnesota Department of Health, Public Health Laboratory | Minnesota Department of Health, Public Health Laboratory | Matt Plumb et al |
| EPI_ISL_1362835,EPI_ISL_1392837 | Missouri State Public Health Laboratory | Missouri State Public Health Laboratory | Matthew Sinn et al |
| EPI_ISL_435129,EPI_ISL_520680,EPI_ISL_520740 | Mohammed Bin Rashid University of Medicine and Health Sciences | Al Jalila Genomics Center | Ahmad Abou Tayoun et al |
| EPI_ISL_903378 | MOH - Jaber Al-Ahmad Hospital (Innovation Research Laboratory) | MOH - Jaber Al-Ahmad Hospital (Innovation Research Laboratory) | Salman Al-Sabah et al |
| EPI_ISL_904015 | Molecular Biology and Virology lab, Faculty of Veterinary Medicine, Jordan University of Science and Technology | Molecular Biology and Virology lab, Faculty of Veterinary Medicine, Jordan University of Science and Technology | Mohammad Hussien Alboom et al |

| | | | |
|---|---|---|---|
| EPI_ISL_2031922,EPI_ISL_2031976 | Molecular diagnostic laboratory of Federal Budget Institution of Science "Central Research Institute of Epidemiology" of The Federal Service on Customers' Rights Protection and Human Well-being Surveillance | Group of Genomics and Postgenomic Technologies of Central Research Institute of Epidemilology | Samojlov A.E. et al |
| EPI_ISL_2006673,EPI_ISL_2101347,EPI_ISL_2105953,EPI_ISL_2105942,EPI_ISL_2105944,EPI_ISL_2105881 | Molecular Diagnostics Pathology Department Mater Dei Hospital Malta | Molecular Diagnostics Pathology Department Mater Dei Hospital Malta | G Zahra et al |
| EPI_ISL_467448,EPI_ISL_482718 | Molecular Diagnostics Services (MDS) | KRISP, KZN Research Innovation and Sequencing Platform | Giandhari J et al |
| EPI_ISL_1365032,EPI_ISL_1365034,EPI_ISL_1365023,EPI_ISL_1663660,EPI_ISL_1663662,EPI_ISL_1663669,EPI_ISL_1663668,EPI_ISL_1663649 | Molecular diagnostic unit for viral haemorrhagic fevers and emerging viruses, Bouaké CHU Laboratory | Molecular diagnostic unit for viral haemorrhagic fevers and emerging viruses, Bouaké CHU Laboratory | Chantal Akoua-Koffi et al |
| EPI_ISL_614350,EPI_ISL_614394,EPI_ISL_614395,EPI_ISL_681833,EPI_ISL_681834,EPI_ISL_1662586,EPI_ISL_1662591 | Molecular diagnostic unit for viral haemorrhagic fevers and emerging viruses, Bouaké CHU Laboratory | Project group Epidemology of Highly Pathogenic Microorganisms, Robert Koch-Institute | Chantal Akoua-Koffi et al |
| EPI_ISL_681684 | Molecular Medicine Laboratory, University of Magallanes | Centro Asistencial Docente y de Investigacion, Universidad de Magallanes | Jorge González et al |
| EPI_ISL_802556,EPI_ISL_802563 | Molecular Microbiology and Food Research Laboratory (MMFRLAB) - Universidad San Sebastián | Facultad de Ciencias de la Vida, UNAB | Dayán Sanhueza et al |

| | | | |
|---|---|---|---|
| EPI_ISL_406844 | Monash Medical Centre | Collaboration between the University of Melbourne at The Peter Doherty Institute for Infection and Immunity, and the Victorian Infectious Disease Reference Laboratory | Caly et al |
| EPI_ISL_1577330 | Montana Public Health Laboratory | Montana Public Health Laboratory | Joy Ritter et al |
| EPI_ISL_889358,EPI_ISL_889368 | Motol University Hospital | Institute of Applied Biotechnologies a.s. | Petr Klempt et al |
| EPI_ISL_2162148 | MPK clinic LLP | Reference laboratory for the control of viral infections | Nazym Tleumbetova et al |
| EPI_ISL_561000,EPI_ISL_561038,EPI_ISL_561054,EPI_ISL_561030,EPI_ISL_561094,EPI_ISL_561164,EPI_ISL_561207,EPI_ISL_561214,EPI_ISL_561197,EPI_ISL_561285,EPI_ISL_561292,EPI_ISL_810982,EPI_ISL_810986,EPI_ISL_811031,EPI_ISL_915421,EPI_ISL_1216072,EPI_ISL_1216076,EPI_ISL_1216077,EPI_ISL_1234529,EPI_ISL_1234530,EPI_ISL_1234531,EPI_ISL_1620171 | MRCG at LSHTM Genomics lab | MRCG at LSHTM Genomics lab | Abdul Karim sesay et al |
| EPI_ISL_471159,EPI_ISL_471160 | MRCG at LSHTM Genomics lab | MRCG at LSHTM Genomics lab | Sesay et al |
| EPI_ISL_1970564,EPI_ISL_1970565,EPI_ISL_1970566,EPI_ISL_1970567,EPI_ISL_1970568 | MRC/UVRI & LSHTM Uganda Research Unit | MRC/UVRI & LSHTM Uganda Research Unit | Matthew Cotten et al |
| EPI_ISL_954299,EPI_ISL_1469322,EPI_ISL_1469387 | MRC/UVRI & LSHTM Uganda Research Unit | Where sequence data have been generated and submitted to GISAID | Matthew Cotten et al |

| | | | |
|---|---|---|---|
| EPI_ISL_802146,EPI_ISL_802392 | MSHS Clinical Microbiology Laboratories | MSHS Pathogen Surveillance Program | Ana S. Gonzalez-Reiche et al |
| EPI_ISL_414476 | MSHS Clinical Microbiology Laboratories | MSHS Pathogen Surveillance Program | Gopi Patel et al |
| EPI_ISL_903850 | MS Public Health Laboratory | Genomics and Discovery, Respiratory Viruses Branch, Division of Viral Diseases, Centers for Disease Control and Prevention | Krista Queen et al |
| EPI_ISL_648014 | MS Public Health Laboratory | Pathogen Discovery, Respiratory Viruses Branch, Division of Viral Diseases, Centers for Disease Control and Prevention | Yan Li et al |
| EPI_ISL_1482832 | MUSC Molecular Pathology Laboratory | MUSC Molecular Pathology Laboratory | Julie W. Hirschhorn et al |
| EPI_ISL_1847214 | MVZ fÃ¼r Laboratoriumsmedizin und Mikrobiologie Koblenz-Mittelrhein (Labor Koblenz) | Robert Koch Institute | ? |
| EPI_ISL_1847444,EPI_ISL_2120586 | MVZ Labor Dr. Limbach & Kollegen GbR | Robert Koch Institute | ? |
| EPI_ISL_860331,EPI_ISL_860499 | MVZ Labor Krone GbR | Center of Medical Microbiology, Virology, and Hospital Hygiene, University of Duesseldorf | Dennis Deschka et al |
| EPI_ISL_2115263 | MVZ Labor Krone GbR | Robert Koch Institute | ? |
| EPI_ISL_447918 | n/a | National Institute of Health. Department of medical Sciences, Ministry of Public Health, Thailand | Pilailuk et al |
| EPI_ISL_2082097 | Nacionalinis maisto ir veterinarijos rizikos vertinimo institutas | National Public Health Surveillance Laboratory | Lukas Zemaitis et al |

| | | | |
|---|---|---|---|
| EPI_ISL_1591098,EPI_ISL_1593728 | NAMRU-6 | Pathogen Discovery, Respiratory Viruses Branch, Division of Viral Diseases, Centers for Disease Control and Prevention | Yan Li et al |
| EPI_ISL_475539 | Narhalsan Sjobo vardcentral | The Public Health Agency of Sweden | Oskar Karlsson Lindsjo et al |
| EPI_ISL_1805741,EPI_ISL_1805787,EPI_ISL_1805933,EPI_ISL_1805932,EPI_ISL_1805959,EPI_ISL_1805962 | National Center for Communicable Diseases (NCCD) National Influenza Center | National Center for Communicable Diseases (NCCD) National Influenza Center | Naranzul Ts et al |
| EPI_ISL_1805697 | National Center for Communicable Diseases (NCCD) National Influenza Center | National Centre for Disease Control (NCDC) National Influenza Center | Naranzul Ts et al |
| EPI_ISL_1718291,EPI_ISL_1718301 | National Center of Disease Control and Prevention of the Republic of Armenia | Institute of Molecular Biology NAS RA, Republic of Armenia, Department of Bioengineering, BioinformaticsInstitute and Molecular Biology IBMPh RAU, Republic of Armenia | Arsen Arakelyan et al |
| EPI_ISL_1854615,EPI_ISL_1854618,EPI_ISL_1854620,EPI_ISL_1854623,EPI_ISL_1854627,EPI_ISL_1854632,EPI_ISL_1854634,EPI_ISL_1854642 | National Center of Disease Control and Prevention of the Republic of Armenia | UW Virology Lab | Pavitra Roychoudhury et al |
| EPI_ISL_454571 | National Center of Expertise | National Center for Expertise, National Center for Biotechnology, Kazakhstan | Abdaliyev Askar et al |

| | | | |
|---|---|---|---|
| EPI_ISL_1301983,EPI_ISL_1401417,EPI_ISL_1401450,EPI_ISL_1608481,EPI_ISL_2081899,EPI_ISL_2081931,EPI_ISL_2081935,EPI_ISL_2082029 | National Center of Infectious and Parasitic Diseases | National Center of Infectious and Parasitic Diseases | Alexiev et al |
| EPI_ISL_1415225,EPI_ISL_1415214,EPI_ISL_1415270,EPI_ISL_1415164,EPI_ISL_1415274,EPI_ISL_1415172,EPI_ISL_1415165,EPI_ISL_1415288,EPI_ISL_1415298,EPI_ISL_1415307,EPI_ISL_1415316,EPI_ISL_1415278,EPI_ISL_1415324,EPI_ISL_1415277,EPI_ISL_1415353,EPI_ISL_1415356,EPI_ISL_1415387,EPI_ISL_1533798,EPI_ISL_1544025,EPI_ISL_1544002,EPI_ISL_1544040,EPI_ISL_1544014 | National Centre For Cell Science | National Centre For Cell Science – INSACOG | Dhiraj Paul et al |
| EPI_ISL_2107058,EPI_ISL_2107080,EPI_ISL_2107063 | National Centre For Cell Science – INSACOG | National Centre For Cell Science | Dhiraj Paul et al |
| EPI_ISL_626569 | National Centre for Communicable Disease (NCCD) National Influenza Center | National Centre for Communicable Disease (NCCD) National Influenza Center | Naranzul Ts et al |
| EPI_ISL_626566 | National Centre for Communication Disease (NCCD) National Influenza Center | National Centre for Communication Disease (NCCD) National Influenza Center | Naranzul Ts et al |
| EPI_ISL_435068 | National Centre for Disease control (NCDC), CSIR-Institute of Genomics and Integrative Biology (CSIR-IGIB) | NCDC/CSIR-IGIB | Pramod Kumar et al |

| | | | |
|---|---|---|---|
| EPI_ISL_1252783 | National Food and Veterinary Risk Assessment Institute | Vilnius University Hospital Santaros Klinikos, Center of Laboratory Medicine | Gytis Dudas et al |
| EPI_ISL_1909863 | National Food and Veterinary Risk Assessment Institute (NMVRVI) | National Public Health Surveillance Laboratory | Lukas Zemaitis et al |
| EPI_ISL_560386 | National Health Laboratory | Botswana Institute for Technology Research and innovation | Kefentse Arnold Tumedi et al |
| EPI_ISL_560385,EPI_ISL_1677705,EPI_ISL_1677715,EPI_ISL_1677714,EPI_ISL_1677737,EPI_ISL_1677719 | National Health Laboratory | Botswana Institute for Technology Research and Innovation | Kefentse Arnold Tumedi et al |
| EPI_ISL_456600,EPI_ISL_456602 | National Health Laboratory, Timor-Leste | Microbiological Diagnostic Unit Public Health Laboratory, The Peter Doherty Institute for Infection and Immunity | Soares da Silva et al |
| EPI_ISL_1407116,EPI_ISL_1407253 | National HIV Reference Laboratory, Ministry of Health, Public Health Institute of Malawi | KRISP, KZN Research Innovation and Sequencing Platform | Mvula B et al |
| EPI_ISL_416429,EPI_ISL_416427 | National Influenza Center, National Institute of Hygiene and Epidemiology (NIHE) | National Influenza Center, National Institute of Hygiene and Epidemiology (NIHE) | Le Quynh Mai et al |
| EPI_ISL_862078,EPI_ISL_1993555 | National Influenza Center, Virology Department | National Influenza Center | A Nejati et al |
| EPI_ISL_862077,EPI_ISL_1014687,EPI_ISL_1993549 | National Influenza Center, Virology Department | National Influenza Center | J Yavarian et al |
| EPI_ISL_1014685,EPI_ISL_1993551 | National Influenza Center, Virology Department | National Influenza Center | K Sadeghi et al |
| EPI_ISL_1014677,EPI_ISL_1014686 | National Influenza Center, Virology Department | National Influenza Center | NZ Shafiei Jandaghi et al |
| EPI_ISL_959284,EPI_ISL_1993547 | National Influenza Center, Virology Department | National Influenza Center | V Salimi et al |

| | | | |
|---|---|---|---|
| EPI_ISL_410301 | National Influenza Centre, National Public Health Laboratory, Kathmandu, Nepal | The University of Hong Kong | Ranjit Sah et al |
| EPI_ISL_402125 | National Institute for Communicable Disease Control and Prevention (ICDC) Chinese Center for Disease Control and Prevention (China CDC) | National Institute for Communicable Disease Control and Prevention (ICDC) Chinese Center for Disease Control and Prevention (China CDC) | Zhang et al |
| EPI_ISL_450298,EPI_ISL_470878 | National Institute for Communicable Diseases of the National Health Laboratory Service | National Institute for Communicable Diseases of the National Health Laboratory Service | Allam M et al |
| EPI_ISL_1910275 | National Institute for Food and Veterinary Risk Assessment (NMVRVI) | National Public Health Surveillance Laboratory | Lukas Zemaitis et al |
| EPI_ISL_469255 | National Institute for Viral Disease Control and Prevention, China CDC | Institute of Viral Disease Control and Prevention, China CDC | Xiang Zhao et al |
| EPI_ISL_591274 | National Institute for Viral Disease Control and Prevention, China CDC | National Institute for Viral Disease Control and Prevention, China CDC | Huilai Ma et al |
| EPI_ISL_469256,EPI_ISL_498692,EPI_ISL_498694,EPI_ISL_850948,EPI_ISL_850949,EPI_ISL_850951 | National Institute for Viral Disease Control and Prevention, China CDC | National Institute for Viral Disease Control and Prevention, China CDC | Xiang Zhao et al |
| EPI_ISL_2107094,EPI_ISL_2107095,EPI_ISL_2107096,EPI_ISL_2107097,EPI_ISL_2107098,EPI_ISL_2107101 | National Institute of Health (NIH) - Federal Government of Somalia | African Centre of Excellence for Genomics of Infectious Diseases (ACEGID), Redeemer's University | Olawoye et al |
| EPI_ISL_574433,EPI_ISL_576386 | National Institute of Health Research and Development | National Institute of Health Research and Development | Pawestri et al |
| EPI_ISL_1915546 | National Institute of Health Research and Development | National Institute of Health Research and Development | Subangkit et al |

| | | | |
|---|---|---|---|
| EPI_ISL_1258440,EPI_ISL_2099853 | National Institute of Infectious Diseases-Prof. Dr. Matei Bals Molecular Diagnostics Laboratory | National Institute of Infectious Diseases-Prof. Dr. Matei Bals Molecular Diagnostics Laboratory | Corina Casangiu et al |
| EPI_ISL_979966,EPI_ISL_979968,EPI_ISL_1279947,EPI_ISL_1279952,EPI_ISL_1279955,EPI_ISL_1279958,EPI_ISL_1279965 | National Institute of Infectious Diseases-Prof. Dr. Matei Bals Molecular Diagnostics Laboratory | National Institute of Infectious Diseases-Prof. Dr. Matei Bals Molecular Diagnostics Laboratory | Leontina Banica et al |
| EPI_ISL_480427 | National Institute of Laboratory Medicine and Referral Center | Bangladesh Council of Scientific and Industrial Research | Shahina Akter et al |
| EPI_ISL_490168 | National Institute of Laboratory Medicine and Referral Center | Genomic Research Lab, BCSIR | Abu Sayeed Mohammad Mahmud et al |
| EPI_ISL_603223 | National Institute of Laboratory Medicine and Referral Center | Genomic Research Lab, BCSIR | Md. Murshed Hasan Sarkar et al |
| EPI_ISL_464160 | National Institute of Laboratory Medicine and Referral Center | Genomic Research Lab, BCSIR | Shahina Akter et al |
| EPI_ISL_1492318,EPI_ISL_1588549 | National Institute of Public Health | National Institute of Public Health | Helena Jirincova et al |
| EPI_ISL_1828716,EPI_ISL_1828724,EPI_ISL_1971079 | National Institute of Public Health | State Veterinary Institute Prague | Nagy et al |
| EPI_ISL_2015692 | National Institute of Public Health - National Institute of Hygiene | 1. National Institute of Public Health - National Institute of Hygiene, Warsaw, Poland 2. Biobank Lab, University of Lodz 3. Laboratory of Respiratory Viruses, Teaching and Clinical Center of the Medical University of Lodz | Dominik Strapagiel et al |
| EPI_ISL_434555 | National Institutes of Health, University of the Philippines Manila | Philippine Genome Center | Carlo M. Lapid et al |

| | | | |
|---|---|---|---|
| EPI_ISL_1266684 | National Laboratory for Health, Environment and Food, OMM, Koper | CISLD (Clinical Institute of Special Laboratory Diagnostics), University Children's Hospital, University Medical Center Ljubljana | Jernej Kovač et al |
| EPI_ISL_1111611 | National Laboratory for Health, Environment and Food, OMM, Kranj | CISLD (Clinical Institute of Special Laboratory Diagnostics), University Children's Hospital, University Medical Center Ljubljana | Jernej Kovač et al |
| EPI_ISL_1664798 | National Laboratory for Health, Environment and Food, OMM, Kranj | NLZOH (National Laboratory for Health, Environment and Food) / CISLD (Clinical Institute of Special Laboratory Diagnostics), University Children's Hospital, University Medical Center Ljubljana | Sandra Janezic et al |
| EPI_ISL_1111747 | National Laboratory for Health, Environment and Food, OMM, Maribor | CISLD (Clinical Institute of Special Laboratory Diagnostics), University Children's Hospital, University Medical Center Ljubljana | Jernej Kovač et al |
| EPI_ISL_512639 | National Laboratory for Influenza/Virology reference laboratory, Public Health Center of the Ministry of Health of Ukraine | Respiratory Virus Unit, Microbiology Services Colindale, Public Health England | PHE Covid Sequencing Team et al |
| EPI_ISL_1191790,EPI_ISL_1191819,EPI_ISL_1191830,EPI_ISL_1191845,EPI_ISL_1191821 | National Microbiology Reference Laboratory | Quadram Institute Bioscience | Tapfumanei Mashe et al |
| EPI_ISL_647971,EPI_ISL_647977,EPI_ISL_647978 | National Microbiology Reference Laboratory | Quadram Institute Bioscience | Thanh Le Viet et al |

| | | | |
|---|---|---|---|
| EPI_ISL_1039984,EPI_ISL_1095615,EPI_ISL_1095621,EPI_ISL_1195207,EPI_ISL_1195201 | National Public Health Center, COVID Laboratory | National Public Health Center, National Biosafety Laboratory | Bernadett Pályi et al |
| EPI_ISL_738086 | National Public Health Laboratory | Department of Medical Microbiology, Hospital Pengajar Universiti Putra Malaysia | Hui-Yee Chee et al |
| EPI_ISL_457835 | National Public Health Laboratory | KEMRI-Wellcome Trust Research Programme/KEMRI-CGMR-C Kilifi | Githinji G. et al 2020 et al |
| EPI_ISL_416866 | National Public Health Laboratory | Malaysia Genome Institute | Mohd Noor Mat Isa et al |
| EPI_ISL_845546,EPI_ISL_845548,EPI_ISL_845549,EPI_ISL_845550,EPI_ISL_845551,EPI_ISL_845554,EPI_ISL_845560 | National Public Health Laboratory, Cameroon | African Centre of Excellence for Genomics of Infectious Diseases (ACEGID), Redeemer's University | Oluniyi P.E. et al |
| EPI_ISL_422430,EPI_ISL_469136,EPI_ISL_475997,EPI_ISL_479588,EPI_ISL_490060,EPI_ISL_512840,EPI_ISL_527362,EPI_ISL_536447 | National Public Health Laboratory, National Centre for Infectious Diseases | National Public Health Laboratory, National Centre for Infectious Diseases | Mak TM et al |
| EPI_ISL_410713 | National Public Health Laboratory, National Centre for Infectious Diseases | National Public Health Laboratory, National Centre for Infectious Diseases | Octavia S et al |

| | | | |
|---|---|---|---|
| EPI_ISL_596460,EPI_ISL_596457,EPI_ISL_605824,EPI_ISL_626636,EPI_ISL_645126,EPI_ISL_728184,EPI_ISL_728194,EPI_ISL_728198,EPI_ISL_768623,EPI_ISL_825067,EPI_ISL_1164354,EPI_ISL_1367560,EPI_ISL_1315626,EPI_ISL_1442952,EPI_ISL_1476997,EPI_ISL_1477006,EPI_ISL_1524797,EPI_ISL_1524796,EPI_ISL_1524799,EPI_ISL_1524800,EPI_ISL_1543977,EPI_ISL_1543978,EPI_ISL_1543981,EPI_ISL_1543980,EPI_ISL_1620161,EPI_ISL_1620158,EPI_ISL_1620162,EPI_ISL_1620166,EPI_ISL_1620159,EPI_ISL_1634424,EPI_ISL_1634426,EPI_ISL_1634428,EPI_ISL_1652105,EPI_ISL_1704836,EPI_ISL_1704835,EPI_ISL_1704837,EPI_ISL_1704840,EPI_ISL_1704841,EPI_ISL_1719868,EPI_ISL_1719885,EPI_ISL_1752677,EPI_ISL_1816938,EPI_ISL_1816942,EPI_ISL_1816943,EPI_ISL_1816949,EPI ISL 181695 | National Public Health Laboratory, National Centre for Infectious Diseases | National Public Health Laboratory, National Centre for Infectious Diseases | Tze Minn Mak et al |
| EPI_ISL_1497491 | National Public Health Organization | National Public Health Organization | Kyriaki Tryfinopoulou et al |
| EPI_ISL_480298,EPI_ISL_480303,EPI_ISL_480309 | National Reference Laboratory "Influenza and acute respiratory diseases" | NRL-HIV | Ivan Ivanov et al |

| | | | |
|---|---|---|---|
| EPI_ISL_1273391,EPI_ISL_1273393,EPI_ISL_1273392,EPI_ISL_1273395,EPI_ISL_1273396,EPI_ISL_1273399,EPI_ISL_1273407,EPI_ISL_1273408 | National Reference Laboratory - Ministry of Health Maseru Lesotho | National Institute for Communicable Diseases of the National Health Laboratory Service | Gorova V et al |
| EPI_ISL_962878 | National Virology Reference Laboratory | National Public Health Laboratory, National Centre for Infectious Diseases | Tze Minn Mak et al |
| EPI_ISL_791326,EPI_ISL_791294 | National Virus Reference Laboratory | Irish Coronavirus Sequencing Consortium - Teagasc Moorepark | Alejandro Abner Garcia Leon et al |
| EPI_ISL_767724 | National Virus Reference Laboratory | Irish Coronavirus Sequencing Consortium - Teagasc Moorepark | Calum Walsh et al |
| EPI_ISL_681903 | National Virus Reference Laboratory | Irish Coronavirus Sequencing Consortium - Teagasc Moorepark | Paul Cotter et al |
| EPI_ISL_1552253,EPI_ISL_1696536 | National Virus Reference Laboratory | National Virus Reference Laboratory | Guerrino Macori et al |
| EPI_ISL_501260,EPI_ISL_525368,EPI_ISL_525376,EPI_ISL_528463 | National Virus Reference Laboratory | National Virus Reference Laboratory | Michael Carr et al |

| | | | |
|---|---|---|---|
| EPI_ISL_1092791,EPI_ISL_1623009,EPI_ISL_1623010,EPI_ISL_1731221,EPI_ISL_1731198,EPI_ISL_1731330,EPI_ISL_1785273,EPI_ISL_1890977,EPI_ISL_1890990,EPI_ISL_1904670,EPI_ISL_1911115,EPI_ISL_1911116,EPI_ISL_1911119,EPI_ISL_1960691,EPI_ISL_1960834,EPI_ISL_1972735,EPI_ISL_2029353,EPI_ISL_2029354,EPI_ISL_2088044,EPI_ISL_2087953,EPI_ISL_2088069,EPI_ISL_2088214,EPI_ISL_2088236,EPI_ISL_2088246,EPI_ISL_2131897,EPI_ISL_2131943,EPI_ISL_2132155,EPI_ISL_2132169,EPI_ISL_2132009,EPI_ISL_2132188,EPI_ISL_2158539,EPI_ISL_2158496 | National Virus Reference Laboratory | National Virus Reference Laboratory | Zoe Yandle et al |
| EPI_ISL_493166 | National Virus Resource Center, Chinese Academy of Sciences, Wuhan 430071, China | Computational Virology Group, Center for Bacteria and Viruses Resources and Bioinformation, Wuhan Institute of Virology, Chinese Academy of Sciences, Wuhan 430071, China | Jianjun Chen et al |
| EPI_ISL_444999,EPI_ISL_445000 | Naval Health Research Center | Naval Medical Research Center Biological Defense Research Directorate | Logan Voegtly et al |
| EPI_ISL_1624278 | Naval Infectious Diseases Diagnostic Laboratory | Naval Medical Research Center Biological Defense Research Directorate | Logan Voegtly et al |

| | | | |
|---|---|---|---|
| EPI_ISL_1939891,EPI_ISL_1939893,EPI_ISL_1939926 | NCCS | inStem NCBS | Uma Ramakrishnan Dasaradhi Palakodeti Aswin SaiNarain et al |
| EPI_ISL_1663375,EPI_ISL_1663376,EPI_ISL_1663409,EPI_ISL_1663418 | NCCS, Pune | Institute of Life Sciences - INSACOG | Sunil K. Raghav et al |
| EPI_ISL_428205,EPI_ISL_428204,EPI_ISL_732824 | Nebraska Public Health Laboratory | UNMC COVID-19 Response Team | UNMC COVID-19 Response Team et al |
| EPI_ISL_754068,EPI_ISL_754069,EPI_ISL_754071 | Nepal Korea Friendship Municipality Hospital | Nepal Health Research Council | Pradip Gyanwali et al |
| EPI_ISL_424875 | NE Public Health Laboratory | Pathogen Discovery, Respiratory Viruses Branch, Division of Viral Diseases, Centers for Disease Control and Prevention | Yan Li et al |
| EPI_ISL_1094737 | NE Public Health Laboratory | Respiratory Viruses Branch, Division of Viral Diseases, Centers for Disease Control and Prevention | Krista Queen et al |
| EPI_ISL_1250385 | Netcare/AMPATH | KRISP, KZN Research Innovation and Sequencing Platform | Giandhari J et al |
| EPI_ISL_1136244,EPI_ISL_1219888,EPI_ISL_1378666,EPI_ISL_1663905,EPI_ISL_1961039 | Nevada State Public Health Laboratory | Nevada State Public Health Laboratory | Andrew Gorzalski et al |
| EPI_ISL_515306,EPI_ISL_515446,EPI_ISL_515449 | Nevada State Public Health Laboratory | Nevada State Public Health Laboratory | Richard Tillett et al |
| EPI_ISL_1064216 | New Mexico Department of Health Scientific Laboratory | Center for Global Health, University of New Mexico Health Sciences Center | Daryl Domman et al |
| EPI_ISL_510879,EPI_ISL_535271 | New Mexico Department of Health Scientific Laboratory | New Mexico Department of Health Scientific Laboratory | Ellie Johnson et al |

| | | | |
|---|---|---|---|
| EPI_ISL_1010700,EPI_ISL_1121988,EPI_ISL_1315070,EPI_ISL_1660480,EPI_ISL_1660404,EPI_ISL_1756024,EPI_ISL_1756026,EPI_ISL_1756027,EPI_ISL_2107448,EPI_ISL_2107447,EPI_ISL_2107444,EPI_ISL_2107458,EPI_ISL_2107442,EPI_ISL_2107529 | New South Wales Health Pathology Royal Prince Alfred Hospital | Microbiology RPAH | Foster et al |
| EPI_ISL_467437,EPI_ISL_487282,EPI_ISL_498058,EPI_ISL_515675,EPI_ISL_515688,EPI_ISL_515738,EPI_ISL_529776,EPI_ISL_602644,EPI_ISL_602653,EPI_ISL_678614 | NHLS-IALCH | KRISP, KZN Research Innovation and Sequencing Platform | Giandhari J et al |
| EPI_ISL_2162308,EPI_ISL_2162324,EPI_ISL_2162342,EPI_ISL_2162347 | NHLS Universitas Academic | UFS Virology | PA Bester et al |
| EPI_ISL_418241,EPI_ISL_420037,EPI_ISL_766869 | NIC Viral Respiratory Unit - Institut Pasteur of Algeria | National Reference Center for Viruses of Respiratory Infections, Institut Pasteur, Paris | Mélanie Albert et al |
| EPI_ISL_872604,EPI_ISL_872610,EPI_ISL_941280 | Nigeria Centre for Disease Control (NCDC) | African Centre of Excellence for Genomics of Infectious Diseases (ACEGID), Redeemer's University | Oluniyi P.E. et al |
| EPI_ISL_455419,EPI_ISL_455431,EPI_ISL_487109,EPI_ISL_527878,EPI_ISL_527890,EPI_ISL_729925,EPI_ISL_729943,EPI_ISL_729945,EPI_ISL_729985,EPI_ISL_730000,EPI_ISL_729978,EPI_ISL_729980,EPI_ISL_730025,EPI_ISL_730040 | Nigeria Centre for Disease Control (NCDC) | African Centre of Excellence for Genomics of Infectious Diseases (ACEGID), Redeemer's University, Ede, Osun State, Nigeria | Oluniyi P.E. et al |

| | | | |
|---|---|---|---|
| EPI_ISL_1035811 | Nigerian Centre for Disease Control (NCDC) | African Centre of Excellence for Genomics of Infectious Diseases (ACEGID), Redeemer's University, Ede | Olawoye et al |
| EPI_ISL_479917 | Niigata City Public Health Research Institute | Pathogen Genomics Center, National Institute of Infectious Diseases | Tsuyoshi Sekizuka et al |
| EPI_ISL_2161784,EPI_ISL_2161801 | NL-Dr. Leonard A. Miller Centre for Health Services | National Microbiology Laboratory (NML) | Anna Majer et al |
| EPI_ISL_960306 | NLZOH, Laboratory for Virology | NLZOH, Laboratory for Virology | Katarina Prosenc (Laboratory for Virology) et al |
| EPI_ISL_422404 | NMIMR, Department of Virology | WACCBIP, University of Ghana | Joyce M. Ngoi et al |
| EPI_ISL_1657140 | NMVRVI | National Public Health Surveillance Laboratory | Lukas Zemaitis et al |
| EPI_ISL_848173,EPI_ISL_848546,EPI_ISL_1919762 | North Dakota Department of Health, Public Health Laboratory | North Dakota Department of Health, Public Health Laboratory | Lisa Wingerter et al |
| EPI_ISL_456395 | North Shore Hospital | Institute of Environmental Science and Research (ESR) | Matt Storey et al |
| EPI_ISL_579407,EPI_ISL_579406 | North Shore Hospital | Institute of Environmental Science and Research (ESR) | Xiaoyun Ren et al |
| EPI_ISL_1482556 | NORTHWELL HEALTH LABORATORIES | Wadsworth Center, New York State Department of Health | Kirsten St. George et al |
| EPI_ISL_1157551 | Northwestern Memorial Hospital | Northwestern University - Ozer Lab | Ramon Lorenzo-Redondo et al |
| EPI_ISL_1192291 | Norwegian Institute of Public Health, Department of Virology | Norwegian Institute of Public Health, Department of Virology | Kathrine Stene-Johansen et al |
| EPI_ISL_416742 | NRL for Influenza, Centrum Epidemiology and Microbiology of National Institute of Public Health, Czech Republic | Charite Universitaetsmedizin Berlin, Institute of Virology | Victor M Corman et al |
| EPI_ISL_940777 | NSPI-CRN de Influenza y otros virus respiratorios | INSPI-Centro de Investigación Multidisciplinaria de la DTIDI | Leandro Patiño et al |

| | | | |
|---|---|---|---|
| EPI_ISL_1272333,EPI_ISL_2162072 | NS-QEII Health Sciences Centre | National Microbiology Laboratory (NML) | Anna Majer et al |
| EPI_ISL_925901,EPI_ISL_960300,EPI_ISL_960301,EPI_ISL_960228,EPI_ISL_1301741,EPI_ISL_1301789,EPI_ISL_1302681,EPI_ISL_1301756,EPI_ISL_1301760,EPI_ISL_1302680 | Nucleic Acid Testing, National Reference Laboratory | GIGA Medical Genomics | Yvan Butera et al |
| EPI_ISL_1117393,EPI_ISL_1117426 | Nucleo de Pesquisa em Inovacao Terapeutica - UFPE | LABBE, Federal University of Pernambuco | Wilson Jose da Silva Junior et al |
| EPI_ISL_1966572 | NUCLEO DE SAUDE VILA FALCAO DE BAURU | Instituto Butantan / Mendelics | Instituto Butantan: Dimas Tadeu Covas et al |
| EPI_ISL_1678079 | NYC Department of Health and Mental Hygiene | Centers for Disease Control and Prevention Division of Viral Diseases, Pathogen Discovery | Mili Sheth et al |
| EPI_ISL_1423467 | NYU Langone Health | Departments of Pathology and Medicine, New York University School of Medicine | Adriana Heguy et al |
| EPI_ISL_1746696 | NZOZ Medyczne Laboratorium Diagnostyczne; | 1. National Institute of Public Health - National Institute of Hygiene; 2. Eurofins Genomics Europe Sequencing GmbH | Wołkowicz Tomasz et al |
| EPI_ISL_1073986,EPI_ISL_1074020 | Office of Diseases Prevention and Control Region 4 Saraburi | COVID-19 Network Investigations (CONI) Alliance | Elizabeth Batty et al |
| EPI_ISL_882772 | Office of Diseases Prevention and Control Region 4 Saraburi | COVID-19 Network Investigations (CONI) Alliance | Kamolthip Atsawawaranunt et al |
| EPI_ISL_526101,EPI_ISL_667418,EPI_ISL_667468,EPI_ISL_832505,EPI_ISL_2080768 | OHSU Lab Services Molecular Microbiology Lab | Oregon SARS-CoV-2 Genome Sequencing Center | Brendan L. O'Connell et al |

| | | | |
|---|---|---|---|
| EPI_ISL_535364 | Oklahoma Animal Disease Diagnostic Laboratory | Oklahoma Animal Disease Diagnostic Laboratory | Sai Narayanan et al |
| EPI_ISL_1094794 | OK Public Health Laboratory, Oklahoma State DOH | Respiratory Viruses Branch, Division of Viral Diseases, Centers for Disease Control and Prevention | Krista Queen et al |
| EPI_ISL_889437 | Olomouc University Hospital | Institute of Applied Biotechnologies a.s. | Petr Klempt et al |
| EPI_ISL_1904868,EPI_ISL_2017488,EPI_ISL_2017485 | OLVZ Aalst | OLVZ Aalst | Anne Vankeerberghen et al |
| EPI_ISL_1532304 | Oman-National Influenza Center | Biotechnology & OMICs Laboratory | Abdul Latif Khan et al |
| EPI_ISL_1532285,EPI_ISL_1517192 | Oman-National Influenza Center | Biotechnology & OMICs Laboratory | Ahmed Al Harrasi et al |
| EPI_ISL_1532286,EPI_ISL_1532289 | Oman-National Influenza Center | Biotechnology & OMICs Laboratory | Intisar Al-Shukri et al |
| EPI_ISL_491165 | Oman-National Influenza Center | Biotechnology & OMICs Laboratory | Sajjad Asaf et al |
| EPI_ISL_491140 | Oman-National Influenza Center | Biotechnology & OMICs Laboratory | Samiha Al-Kharusi et al |
| EPI_ISL_518830 | Oman-National Influenza Center | Biotechnology & OMICs Laboratory, Natural & Medical Sciences Research Center, University of Nizwa | Sajjad Asaf et al |
| EPI_ISL_766569 | Oman-National Influenza Center | Oman-National Influenza Center | Samiha Al-Kharusi et al |
| EPI_ISL_491968,EPI_ISL_491999,EPI_ISL_1795046 | Oman-NIC | Department of Microbiology and Immunology-SQUH | Fahad Zadjali et al |
| EPI_ISL_1819073,EPI_ISL_1819076,EPI_ISL_1819077 | Oman-NIC | Oman-National Influenza Center-Department of Microbiology and Immunology-SQUH | Samira Al-Maruqi et al |
| EPI_ISL_457704,EPI_ISL_457938,EPI_ISL_457995 | Oman-NIC | Oman-NIC | Samira Al-Maruqi et al |

| | | | |
|---|---|---|---|
| EPI_ISL_1381300,EPI_ISL_1443667,EPI_ISL_1443670,EPI_ISL_1443671,EPI_ISL_1968638 | Omics Sciences Laboratory | Omics Sciences Laboratory | Derly Andrade Molina et al |
| EPI_ISL_639912 | Omsk Research Institute of Natural Focal Infections | WHO National Influenza Centre Russian Federation | Artem Fadeev et al |
| EPI_ISL_1040028 | Original detection - Virology Unit, Institut Pasteur du Cambodge; Sequencing - US National Institute of Allergy and Infectious Diseases Cambodia | Virology Unit, Institut Pasteur du Cambodge | Jennifer Bohl et al |
| EPI_ISL_419558 | OR State PHL-Virology/Immunology Section | Pathogen Discovery, Respiratory Viruses Branch, Division of Viral Diseases, Centers for Disease Control and Prevention | Ying Tao et al |
| EPI_ISL_708118 | Oslo University Hospital, Department of Medical Microbiology | Norwegian Institute of Public Health, Department of Virology | Kathrine Stene-Johansen et al |
| EPI_ISL_458069 | Osmania Medical College | CSIR-Centre for Cellular and Molecular Biology | Shashikala Reddy et al |
| EPI_ISL_747462 | Ospedale San Bonifacio | Istituto Zooprofilattico Sperimentale delle Venezie | Adelaide Milani et al |
| EPI_ISL_1919755,EPI_ISL_1919757 | Ospedale Santa Caterina Novella | Istituto Zooprofilattico Sperimentale della Puglia e della Basilicata | Parisi A. et al |
| EPI_ISL_549152,EPI_ISL_2142765 | Ostfold Hospital Trust - Kalnes, Centre for Laboratory Medicine, Section for gene technology and infection serology | Norwegian Institute of Public Health, Department of Virology | Kathrine Stene-Johansen et al |
| EPI_ISL_1366744 | OUCRU | OUCRU | Nguyen Van Vinh Chau et al |
| EPI_ISL_2140726 | Oudtshoorn Hospital wc OUD | NHLS/UCT | Arash Iranzadeh et al |
| EPI_ISL_1416967,EPI_ISL_1416968,EPI_ISL_1416980,EPI_ISL_1416988 | Outre mer | National Reference Center for Viruses of Respiratory Infections, Institut Pasteur, Paris | Marion Barbet et al |
| EPI_ISL_1273214 | Oxford University Clinical Research Unit (OUCRU) | Oxford University Clinical Research Unit (OUCRU) | Nguyen Van Vinh Chau et al |

| | | | |
|---|---|---|---|
| EPI_ISL_424881 | PA Department of Health, Bureau of Laboratories | Pathogen Discovery, Respiratory Viruses Branch, Division of Viral Diseases, Centers for Disease Control and Prevention | Yan Li et al |
| EPI_ISL_596543,EPI_ISL_596552,EPI_ISL_596560 | Palestinian Ministry of Health | Molecular Genetics Lab | Nouar Qutob et al |
| EPI_ISL_708194 | Pamukkale University Hospital | Pamukkale University Department of Medical Genetics | Onur TOKGUN et al. |
| EPI_ISL_1470937,EPI_ISL_1470959,EPI_ISL_1828556 | Pandemic Response Lab - NYC | Pandemic Response Lab, R&D | Henry Lee et al |
| EPI_ISL_2155416 | Parkway Medical and Diagnostic Center | Philippine Genome Center | Francis A. Tablizo et al |
| EPI_ISL_2035988 | Pasteur Institute - Laboratory of Clinical Virology | Pasteur Institute - Laboratory of Clinical Virology | Wasfi Fares et al |
| EPI_ISL_2013036 | Pathcare-Vermaak Centurion | National Institute for Communicable Diseases of the National Health Laboratory Service | Amoako DG et al |
| EPI_ISL_456163,EPI_ISL_456194,EPI_ISL_456195,EPI_ISL_456215,EPI_ISL_456223,EPI_ISL_456224,EPI_ISL_456301 | PathLab Bay of Plenty | Institute of Environmental Science and Research (ESR) | Matt Storey et al |
| EPI_ISL_582019,EPI_ISL_622805,EPI_ISL_622806 | PathLab Bay of Plenty | Institute of Environmental Science and Research (ESR) | Xiaoyun Ren et al |

| | | | |
|---|---|---|---|
| EPI_ISL_687765,EPI_ISL_691811,EPI_ISL_692046,EPI_ISL_900781,EPI_ISL_902443,EPI_ISL_1127322,EPI_ISL_1127375,EPI_ISL_1128467,EPI_ISL_1427640,EPI_ISL_1430417,EPI_ISL_1928350,EPI_ISL_1932860,EPI_ISL_1933024,EPI_ISL_1933230,EPI_ISL_1934050 | Pathogen Genomics Center, National Institute of Infectious Diseases | Pathogen Genomics Center, National Institute of Infectious Diseases | Tsuyoshi Sekizuka et al |
| EPI_ISL_513178 | Pathogen Genomics Lab King Abdullah University of Science and Technology(KAUST) | Pathogen Genomics Lab King Abdullah University of Science and Technology(KAUST) | Amit Kumar Subudhi et al |
| EPI_ISL_512922,EPI_ISL_513151 | Pathogen Genomics Lab King Abdullah University of Science and Technology(KAUST) | Pathogen Genomics Lab King Abdullah University of Science and Technology(KAUST) | Fadwa Alofi et al |
| EPI_ISL_751207 | Pathogen Genomics Lab King Abdullah University of Science and Technology(KAUST) | Pathogen Genomics Lab King Abdullah University of Science and Technology(KAUST) | Fathia Ben Rached et al |
| EPI_ISL_678239 | Pathogen Genomics Lab King Abdullah University of Science and Technology(KAUST) | Pathogen Genomics Lab King Abdullah University of Science and Technology(KAUST) | Muhammad Shuaib et al |
| EPI_ISL_678221,EPI_ISL_751210 | Pathogen Genomics Lab King Abdullah University of Science and Technology(KAUST) | Pathogen Genomics Lab King Abdullah University of Science and Technology(KAUST) | Sara Mfarrej et al |
| EPI_ISL_526152 | Pathology North - Hunter - NSW Health Pathology | NSW Health Pathology - Institute of Clinical Pathology and Medical Research; Westmead Hospital; University of Sydney | CIDM-PH et al. |

| | | | |
|---|---|---|---|
| EPI_ISL_591493 | Pathology North - Royal North Shore Hospital - NSW Health Pathology | NSW Health Pathology - Institute of Clinical Pathology and Medical Research; Westmead Hospital; University of Sydney | CIDM-PH et al. |
| EPI_ISL_407894,EPI_ISL_407896,EPI_ISL_410717 | Pathology Queensland | Public Health Virology Laboratory | Ben Huang et al |
| EPI_ISL_414414 | Pathology Queensland | Public Health Virology Laboratory | Bixing Huang et al |
| EPI_ISL_451539,EPI_ISL_455061,EPI_ISL_455069,EPI_ISL_513332,EPI_ISL_526118,EPI_ISL_591499,EPI_ISL_591501,EPI_ISL_593682,EPI_ISL_767937 | Pathology West - NSW Health Pathology | NSW Health Pathology - Institute of Clinical Pathology and Medical Research; Westmead Hospital; University of Sydney | CIDM-PH et al. |
| EPI_ISL_420456,EPI_ISL_470851,EPI_ISL_470868 | PathWest Laboratory Medicine WA | PathWest Laboratory Medicine WA | Chisha Sikazwe et al |

| | | | |
|---|---|---|---|
| EPI_ISL_512715,EPI_ISL_512717,EPI_ISL_512718,EPI_ISL_512719,EPI_ISL_512723,EPI_ISL_512758,EPI_ISL_594171,EPI_ISL_596681,EPI_ISL_596694,EPI_ISL_596719,EPI_ISL_596743,EPI_ISL_596828,EPI_ISL_596834,EPI_ISL_596840,EPI_ISL_605841,EPI_ISL_605844,EPI_ISL_672638,EPI_ISL_708762,EPI_ISL_794673,EPI_ISL_794677,EPI_ISL_794683,EPI_ISL_794686,EPI_ISL_794690,EPI_ISL_794716,EPI_ISL_794711,EPI_ISL_794734,EPI_ISL_794726,EPI_ISL_810969,EPI_ISL_933798,EPI_ISL_933801,EPI_ISL_1017678,EPI_ISL_1069392,EPI_ISL_1069393,EPI_ISL_1184507,EPI_ISL_1198803,EPI_ISL_1295932,EPI_ISL_1295933,EPI_ISL_1366740,EPI_ISL_1416324,EPI_ISL_1508993,EPI_ISL_1669125,EPI_ISL_1750967 | PathWest Laboratory Medicine WA | PathWest Laboratory Medicine WA Microbial Surveillance Unit | PathWest Laboratory Medicine WA Microbial Surveillance Unit et al |
| EPI_ISL_1911192 | PathWest Laboratory Medicine WA | PathWest Laboratory Medicine WA Microbial Surveillance UNit | PathWest Laboratory Medicine WA Microbial Surveillance UNit et al |
| EPI_ISL_469814 | PHE South West Regional Laboratory, National Infection Service | Wellcome Sanger Institute for the COVID-19 Genomics UK (COG-UK) consortium | Stephanie Hutchings et al |
| EPI_ISL_2153841 | Philippine Genome Center - Biobank (NCR) | Philippine Genome Center | Francis A. Tablizo et al |
| EPI_ISL_2155110 | Philippine Red Cross - Port Area | Philippine Genome Center | Francis A. Tablizo et al |

| | | | |
|---|---|---|---|
| EPI_ISL_968212,EPI_ISL_1196414,EPI_ISL_1300531,EPI_ISL_1300528,EPI_ISL_1300527,EPI_ISL_1465881,EPI_ISL_1516887,EPI_ISL_1563665,EPI_ISL_1563670,EPI_ISL_1563660,EPI_ISL_1910859,EPI_ISL_2137036 | PHV-FSS | PHV-FSS | Son Nguyen et al |
| EPI_ISL_968213 | PHV-FSS | PHV-FSS | Son Nguyen et al. |
| EPI_ISL_1824607 | PKC Jagakarsa | National Institute of Health Research and Development | Vivi Setiawaty et al |
| EPI_ISL_1498376,EPI_ISL_1963398,EPI_ISL_2047647,EPI_ISL_2047648,EPI_ISL_2047650,EPI_ISL_2047659,EPI_ISL_2047662,EPI_ISL_2131296,EPI_ISL_2131297,EPI_ISL_2131298,EPI_ISL_2131303,EPI_ISL_2131310 | Platform BIS UZA/UAntwerpen | Labo Klinische Biologie, UZA | Jasmine Coppens et al |
| EPI_ISL_1591163,EPI_ISL_1745201,EPI_ISL_1745202,EPI_ISL_1745199,EPI_ISL_2016711,EPI_ISL_2016712,EPI_ISL_2016714,EPI_ISL_2016715 | Platform BIS UZA/UAntwerpen | Labo Klinische Biologie, UZA | Marie Le Mercier et al |
| EPI_ISL_1402426,EPI_ISL_1439584 | Private clinic of Biogen Med, Tashkent, Uzbekistan | Center of Genomics and bioinformatics, Bioinformatics laboratory | Mirzakamol S Ayubov et al |
| EPI_ISL_513530 | Programa de Oncovirologia, Instituto Nacional de Câncer | Programa de Oncovirologia, Instituto Nacional de Câncer | Juliana D. Siqueira et al |
| EPI_ISL_1660340,EPI_ISL_1789673 | Pro-Vitam Diagnostics and Research Laboratory | Pro-Vitam Diagnostics and Research Laboratory | Szilard N. Fejer et al |
| EPI_ISL_1824604 | PRVKP FK UI | National Institute of Health Research and Development | Vivi Setiawaty et al |

| | | | |
|---|---|---|---|
| EPI_ISL_1209253 | Public Health Authority of the Slovak Republic | Bergthaler laboratory, CeMM Research Center for Molecular Medicine of the Austrian Academy of Sciences | Lukas Endler et al |
| EPI_ISL_1749544 | Public Health Authority of the Slovak Republic | Laboratory of Genomics and Bioinformatics, Comenius University Science Park | Tatiana Sedláčková et al |
| EPI_ISL_1112753,EPI_ISL_1112316 | Public Health Center of Ukraine | Charité Universitätsmedizin Berlin, Institute of Virology | Victor M Corman et al |
| EPI_ISL_1312018 | Public Health Institute of Zagreb County | Croatian Institute of Public Health | Irena Tabain et al |
| EPI_ISL_636973 | Public Health Lab | Public Health Lab | Alwasti et al |
| EPI_ISL_425177 | Public Health Ontario | Public Health Agency of Canada - National Microbiology Laboratory | Amrit S. Boese et al |
| EPI_ISL_513313 | Public Health, United States Air Force School of Aerospace Medicine | Public Health, United States Air Force School of Aerospace Medicine | Fries et al |
| EPI_ISL_985235,EPI_ISL_1061310,EPI_ISL_1061309,EPI_ISL_1121039 | Public Health Virology-Forensic and Scientific Services | Public Health Virology-Forensic and Scientific Services | Son Nguyen et al |
| EPI_ISL_944744,EPI_ISL_944748,EPI_ISL_1300533,EPI_ISL_1306127,EPI_ISL_1300526,EPI_ISL_1306132,EPI_ISL_1306135 | Public Health Virology-Forensic and Scientific Services (PHV-FSS) | Public Health Virology-Forensic and Scientific Services (PHV-FSS) | Son Nguyen et al |

| | | | |
|---|---|---|---|
| EPI_ISL_576135,EPI_ISL_593613,EPI_ISL_593634,EPI_ISL_593641,EPI_ISL_593639,EPI_ISL_639813,EPI_ISL_639818,EPI_ISL_639819,EPI_ISL_639750,EPI_ISL_641310,EPI_ISL_693276,EPI_ISL_693290,EPI_ISL_849665,EPI_ISL_849682,EPI_ISL_849683,EPI_ISL_849684,EPI_ISL_849691,EPI_ISL_849692,EPI_ISL_849694,EPI_ISL_849761,EPI_ISL_849751,EPI_ISL_849759 | Public Health Virology Laboratory, Forensic and Scientific Services (PHV-FSS) | Public Health Virology Laboratory, Forensic and Scientific Services (PHV-FSS) | Son Nguyen et al |
| EPI_ISL_579655,EPI_ISL_1055661,EPI_ISL_1055675,EPI_ISL_1055676 | QEII Health Sciences Centre | National Microbiology Laboratory (NML) | Anna Majer et al |
| EPI_ISL_530250,EPI_ISL_530276,EPI_ISL_530227,EPI_ISL_530229,EPI_ISL_530232,EPI_ISL_530237,EPI_ISL_530258,EPI_ISL_530239 | Queensland Health Forensic and Scientific Services, Public Health Virology | Public Health Virology Laboratory, Forensic and Scientific Services, Queensland Health | Son Nguyen et al |
| EPI_ISL_1424680,EPI_ISL_1424786 | Queensland Medical Laboratories | Melbourne Diagnostic Unit Public Health Laboratory (MDU-PHL) | Palou et al |
| EPI_ISL_1424520,EPI_ISL_1424521,EPI_ISL_1424528,EPI_ISL_1424546,EPI_ISL_1424582,EPI_ISL_1424621,EPI_ISL_1424624,EPI_ISL_1424632,EPI_ISL_1424635 | Queensland Medical Laboratories | Victorian Infectious Diseases Reference Laboratory (VIDRL) and the Melbourne Diagnostic Unit Public Health Laboratory (MDU-PHL) | Palou et al |
| EPI_ISL_498696,EPI_ISL_571018,EPI_ISL_571938,EPI_ISL_572053,EPI_ISL_845666,EPI_ISL_876887,EPI_ISL_937064 | Quest Diagnostics | Quest Diagnostics | Rosenthal et al |

| | | | |
|---|---|---|---|
| EPI_ISL_1552968,EPI_ISL_1648249,EPI_ISL_1648208,EPI_ISL_1738201,EPI_ISL_1753145,EPI_ISL_1762028,EPI_ISL_1798071,EPI_ISL_1840480,EPI_ISL_2047060,EPI_ISL_2133436,EPI_ISL_2143130 | Quest Diagnostics Incorporated | Centers for Disease Control and Prevention Division of Viral Diseases, Pathogen Discovery | Dakota Howard et al |
| EPI_ISL_1220377,EPI_ISL_1220464 | Quest Diagnostics Incorporated | Centers for Disease Control and Prevention Division of Viral Diseases, Pathogen Discovery | Peter W. Cook et al |
| EPI_ISL_1139353 | Quest Diagnostics Incorporated | Respiratory Viruses Branch, Division of Viral Diseases, Centers for Disease Control and Prevention | Peter W. Cook et al |
| EPI_ISL_569615 | Quick Care Huron | South Dakota Public Health Laboratory | Matt Plumb et al |
| EPI_ISL_878700 | Rady's Childrens Hospital | Andersen lab at Scripps Research | SEARCH Alliance San Diego with Nanda Radamchar et al |
| EPI_ISL_637110,EPI_ISL_637112,EPI_ISL_637113 | Rafik Hariri University Hospital | Microbial Pathogenomics Lab | Georgi Merhi et al |
| EPI_ISL_450512,EPI_ISL_450514 | Rafik Hariri University Hospital | Rafik Hariri University Hospital | Rita Feghali et al |
| EPI_ISL_447021,EPI_ISL_1073976,EPI_ISL_2104750 | Ramathibodi Hospital | COVID-19 Network Investigations (CONI) Alliance | Elizabeth Batty et al |
| EPI_ISL_2000573,EPI_ISL_2000595,EPI_ISL_2000597 | Rami Kantor lab | Rami Kantor lab | Josephine Darpolor et al |
| EPI_ISL_1467170,EPI_ISL_1631914,EPI_ISL_1740633,EPI_ISL_1790372,EPI_ISL_1829999,EPI_ISL_2006040,EPI_ISL_2005618,EPI_ISL_2092214 | Randox Laboratories | Wellcome Sanger Institute for the COVID-19 Genomics UK (COG-UK) Consortium | Randox Laboratories et al |

| | | | |
|---|---|---|---|
| EPI_ISL_2157347 | Red - Regional de Vigilancia Genómica del COVID-19 | Laboratory of Respiratory Viruses and Measles, Oswaldo Cruz Institute, FIOCRUZ | Paola Resende et al |
| EPI_ISL_1415407,EPI_ISL_1415408 | Reference Laboratory of the Ministry of Health | Laboratory of Respiratory Viruses and Measles, Oswaldo Cruz Institute, FIOCRUZ | Paola Resende et al |
| EPI_ISL_768533 | Regional Medical Sciences Center 5 Samut Songkhram | National Institute of Health, Department of Medical Sciences, Ministry of Public Health, Thailand | Pilailuk Okada et al |
| EPI_ISL_708800,EPI_ISL_708802 | Regional medical sciences center 6 chonburi | National Institute of Health, Department of Medical Sciences, Ministry of Public Health, Thailand | Pilailuk Okada et al |
| EPI_ISL_2110840 | Reitor da Universidade Jean Piaget Guine-Bissau | MRCG at LSHTM, Genomics lab | Aladje Balde et al |
| EPI_ISL_491474 | Research Institute for Tropical Medicine | Research Institute for Tropical Medicine | Ma. Angelica Tujan et al |
| EPI_ISL_2155097 | Research Institute for Tropical Medicine, Inc. (RITM) | Philippine Genome Center | Francis A. Tablizo et al |
| EPI_ISL_2105985 | Respiratory Viruses Branch, Centers for Disease Control and Prevention | Respiratory Viruses Branch, Centers for Disease Control and Prevention | Howard et al |
| EPI_ISL_747086,EPI_ISL_747171 | Respiratory Viruses Branch, Centers for Disease Control and Prevention | Respiratory Viruses Branch, Centers for Disease Control and Prevention | Queen et al |
| EPI_ISL_414523,EPI_ISL_407073 | Respiratory Virus Unit, Microbiology Services Colindale, Public Health England | Respiratory Virus Unit, Microbiology Services Colindale, Public Health England | Monica Galiano et al |

| | | | |
|---|---|---|---|
| EPI_ISL_416480 | R. G. Lugar Center for Public Health Research, National Center for Disease Control and Public Health (NCDC) of Georgia. | R. G. Lugar Center for Public Health Research, National Center for Disease Control and Public Health (NCDC) of Georgia. | Ann Machablishvili et al |
| EPI_ISL_415641,EPI_ISL_415643 | R. G. Lugar Center for Public Health Research, National Center for Disease Control and Public Health (NCDC) of Georgia. | R. G. Lugar Center for Public Health Research, National Center for Disease Control and Public Health (NCDC) of Georgia. | Nato Kotaria et al |
| EPI_ISL_872690,EPI_ISL_872727 | Rhode Island Department of Health | Infectious Disease Program, Broad Institute of Harvard and MIT | Lemieux et al |
| EPI_ISL_1970713 | "Riga East Clinical University Hospital, National Microbiology Reference Laboratory" | "Riga East Clinical University Hospital, National Microbiology Reference Laboratory; Eurofins Genomics Europe Sequencing GmbH" | Ģirts Šķenders et al |
| EPI_ISL_1590583 | Riga East University Hospital-National Microbiology Reference Laboratory; Eurofins Genomics Europe Sequencing GmbH | Riga East University Hospital-National Microbiology Reference Laboratory; Eurofins Genomics Europe Sequencing GmbH | Ģirts Šķenders et al |
| EPI_ISL_485201,EPI_ISL_485247 | River Road Testing Lab | Ginkgo Bioworks Clinical Laboratory | Rebecca C. Christofferson et al |
| EPI_ISL_468143,EPI_ISL_468145 | [Romania, Bucharest] National Institute for Infectious Diseases "Prof. Dr. Matei Balș" | [Romania, Bucharest] National Institute for Infectious Diseases "Prof. Dr. Matei Balș" | Leontina Banica et al |
| EPI_ISL_514136 | Rondônia Central Public Health Laboratory (LACEN/RO), vinculated to State Health Secretariat of Rondônia (SESAU/RO) | Molecular Virology Laboratory of Oswaldo Cruz Foundation of Rondônia | Luan Felipo Botelho-Souza et al |

| | | | |
|---|---|---|---|
| EPI_ISL_521860,EPI_ISL_521862,EPI_ISL_521861,EPI_ISL_779406,EPI_ISL_779410,EPI_ISL_812423,EPI_ISL_854745 | Royal Darwin Hospital Pathology | MDU-PHL | Meumann et al |
| EPI_ISL_577607 | Royal Hobart Hospital | Royal Hobart Hospital | Cooley L. et al |
| EPI_ISL_522565,EPI_ISL_522569,EPI_ISL_522608,EPI_ISL_522679,EPI_ISL_522686,EPI_ISL_522693,EPI_ISL_522705,EPI_ISL_522756,EPI_ISL_522762,EPI_ISL_522764 | Royal Hobart Hospital Microbiology Department | MDU-PHL | Cooley L. et al |
| EPI_ISL_454509 | RSE "National Center for Biotechnology" | RSE "National Center for Biotechnology" | Alexandr Shevtsov et al |
| EPI_ISL_574607 | RS Husada | Eijkman Institute for Molecular Biology, Ministry of Research and Technology/National Agency for Research and Innovation | Frilasita A Yudhaputri et al |
| EPI_ISL_1622430 | RS Mitra Keluarga Gading Serpong | Eijkman Institute for Molecular Biology, Ministry of Research and Technology/National Agency for Research and Innovation | Muhammad Rezki Rasyak et al |
| EPI_ISL_1622432 | RS Sentra Medika Cikarang | Eijkman Institute for Molecular Biology, Ministry of Research and Technology/National Agency for Research and Innovation | Muhammad Rezki Rasyak et al |
| EPI_ISL_529964 | RSUD Bangil Pasuruan | Institute of Tropical Disease, Universitas Airlangga | Aldise M Nastri et al |
| EPI_ISL_458081 | RSUD Bangil Pasuruan | Institute of Tropical Disease, Universitas Airlangga | Jezzy R Dewantari et al |

| | | | |
|---|---|---|---|
| EPI_ISL_2047572 | RSUD Siti Fatimah Prov Sumatera Selatan | National Institute of Health Research and Development | Hana Apsari Pawestri et al |
| EPI_ISL_2047517 | RSUP H Adam Malik Medan | National Institute of Health Research and Development | Kartika Dewi Puspa et al |
| EPI_ISL_1969246,EPI_ISL_1969247 | Rumah Sakit Umum Daerah Palangkaraya | National Institute of Health Research and Development | Subangkit et al |
| EPI_ISL_491298 | Rural Health Unit - Calauan, Laguna | Research Institute for Tropical Medicine | Tujan et al |
| EPI_ISL_1427492 | Saitama Prefectural Institute of Public Health | Pathogen Genomics Center, National Institute of Infectious Diseases | Tsuyoshi Sekizuka et al |
| EPI_ISL_1933482 | Sakai City Institute of Public Health | Pathogen Genomics Center, National Institute of Infectious Diseases | Tsuyoshi Sekizuka et al |
| EPI_ISL_749238,EPI_ISL_749474,EPI_ISL_749706,EPI_ISL_751011,EPI_ISL_750172,EPI_ISL_750173 | Sanatorio Americano | Institut Pasteur de Montevideo | Daiana Mir et al |
| EPI_ISL_667032 | San Diego County Public Health Laboratory | Andersen lab at Scripps Research | SEARCH Alliance San Diego with Tracy Basler et al |
| EPI_ISL_2035746 | San Gallicano Dermatological Institute I.F.O. | INMI Lazzaro Spallanzani IRCCS | F Messina et al |
| EPI_ISL_2035749 | San Gallicano Dermatological Institute I.F.O. | INMI Lazzaro Spallanzani IRCCS | O Butera et al |
| EPI_ISL_1477049 | Sanitary-Epidemiological And Public Health Department Of Tashkent Region, Uzbekistan | Center of Genomics and bioinformatics, Bioinformatics laboratory | Mirzakamol S Ayubov et al |
| EPI_ISL_1821470 | Santa Clara County Public Health Laboratory | Chan-Zuckerberg Biohub | CZB Cliahub Consortium et al |

| | | | |
|---|---|---|---|
| EPI_ISL_451088,EPI_ISL_451127,EPI_ISL_467987,EPI_ISL_467989,EPI_ISL_467994,EPI_ISL_468024,EPI_ISL_468037,EPI_ISL_468039,EPI_ISL_468041,EPI_ISL_483075,EPI_ISL_483110,EPI_ISL_492132,EPI_ISL_508606,EPI_ISL_510544,EPI_ISL_513912,EPI_ISL_513913,EPI_ISL_602577,EPI_ISL_622762,EPI_ISL_654806,EPI_ISL_732961,EPI_ISL_755570,EPI_ISL_755571,EPI_ISL_812358,EPI_ISL_1029956,EPI_ISL_1704786,EPI_ISL_1704675,EPI_ISL_1704803,EPI_ISL_1704804,EPI_ISL_1704806,EPI_ISL_1706367 | SA Pathology | SA Pathology | Lex Leong et al |
| EPI_ISL_455603 | SA Pathology | VPRL | Beard et al |
| EPI_ISL_1214244 | SARS-CoV-2 Sequencing Castilla y Leon-Spain Consortium | SARS-CoV-2 Sequencing Castilla y Leon-Spain Consortium | Antonio Orduña-Domingo et al |

| | | | |
|---|---|---|---|
| EPI_ISL_1427800,EPI_ISL_1927400,EPI_ISL_1927401,EPI_ISL_1927403,EPI_ISL_1927404,EPI_ISL_1927405,EPI_ISL_1927407,EPI_ISL_1927408,EPI_ISL_1927410,EPI_ISL_1927411,EPI_ISL_1927422,EPI_ISL_1927423,EPI_ISL_1927433,EPI_ISL_1927424,EPI_ISL_1927436,EPI_ISL_1927426,EPI_ISL_1927438,EPI_ISL_1927439,EPI_ISL_1927441,EPI_ISL_1927442,EPI_ISL_1927444,EPI_ISL_1927445,EPI_ISL_2131670,EPI_ISL_2131691,EPI_ISL_2131693,EPI_ISL_2131684,EPI_ISL_2131695,EPI_ISL_2131697,EPI_ISL_2131707,EPI_ISL_2131723,EPI_ISL_2131726,EPI_ISL_2131731,EPI_ISL_2131732,EPI_ISL_2131734,EPI_ISL_2131735,EPI_ISL_2131737,EPI_ISL_2131750,EPI_ISL_2131743,EPI_ISL_2131772,EPI_ISL_2131775,EPI_ISL_2131778,EPI_ISL_2131781,EPI_ISL_2131782,EPI_ISL_2131783,EPI_ISL_2131790,EPI_IS | SARS-CoV-2 testing team, National Institute of Infectious Diseases | Pathogen Genomics Center, National Institute of Infectious Diseases | Tsuyoshi Sekizuka et al |
| EPI_ISL_491108 | SC Department of Health and Environmental Control | SC Department of Health and Environmental Control | Flores et al |
| EPI_ISL_1678135 | SC Dept of Health and Env. Control-Bureau of Laboratories | Centers for Disease Control and Prevention Division of Viral Diseases, Pathogen Discovery | Mili Sheth et al |

| | | | |
|---|---|---|---|
| EPI_ISL_1009128,EPI_ISL_1009160 | School of Pharmacy | School of Pharmacy | Ahmed Kandeil et al |
| EPI_ISL_661189,EPI_ISL_661190,EPI_ISL_904009 | Scientific Veterinary Institute Novi Sad | Veterinary Specialized Institute "Kraljevo", Serbia | Vidanovic et al |
| EPI_ISL_1647029 | SC (UCO) Igiene e Sanità Pubblica (funzione integrata con SC Microbiologia e Virologia) | ARGO Laboratorio Genomica ed Epigenomica | Licastro D et al |
| EPI_ISL_420121 | Servicio de Microbiología. Consorcio Hospital General Universitario de Valencia | Sequencing and Bioinformatics Service and Molecular Epidemiology Research Group. FISABIO-Public Health | Vicente Soriano Chirona et al |
| EPI_ISL_468934,EPI_ISL_468938 | Servicio de Microbiología. Hospital Universitario Donostia. OSI Donostialdea. Área de Enfermedades Infecciosas, Grupo de Infección Respiratoria y Resistencia Antimicrobiana. Instituto de Investigación Sanitaria Biodonostia. | SeqCOVID-SPAIN consortium/IBV(CSIC) | Gustavo Cilla et al |
| EPI_ISL_510058 | Servicio de Microbiología. HRU de Málaga. Servicio Andaluz de Salud | SeqCOVID-SPAIN consortium/IBV(CSIC) | Inmaculada de Toro Peinado. MªConcepción Mediavilla Gradolph. Begoña Palop Borrás et al |
| EPI_ISL_420599,EPI_ISL_849299,EPI_ISL_849301,EPI_ISL_849317,EPI_ISL_849321,EPI_ISL_849358,EPI_ISL_856756,EPI_ISL_856781,EPI_ISL_2104757,EPI_ISL_2104770,EPI_ISL_2104777,EPI_ISL_2105572,EPI_ISL_2133401,EPI_ISL_2136073,EPI_ISL_2140071,EPI_ISL_2158778,EPI_ISL_2158846,EPI_ISL_2158739,EPI_ISL_2158768 | Servicio Virosis Respiratorias-Departamento Virología-INEI | Instituto Nacional Enfermedades Infecciosas C.G.Malbran | Baumeister E. et al |

| | | | |
|---|---|---|---|
| EPI_ISL_436728 | SERVIZIO DI IGIENE E SANITÀ PUBBLICA ASL Teramo | Istituto Zooprofilattico Sperimentale dell'Abruzzo e Molise "G.Caporale" | Lorusso A et al |
| EPI_ISL_849659 | Servizio Igiene Epidemiologia e Sanità Pubblica (SIESP)-L'Aquila | Istitituto Zooprofilattico Sperimentale dell'Abruzzo e Molise "G.Caporale" | Lorusso A et al |
| EPI_ISL_497950 | Shaoxing CDC | Zhejiang Provincial Center for Disease Control and Prevention | Yin Chen et al |
| EPI_ISL_463902 | Shaoxing Center for Disease Control and Prevention | Department of Pathology and Laboratory Medicine, University of California Los Angeles | Jinkun Chen et al |
| EPI_ISL_636120 | Sharp HealthCare Laboratory | Andersen lab at Scripps Research | SEARCH Alliance San Diego with Aaron Harding et al |
| EPI_ISL_582125 | Sheikh Khalifa Medical City | Molecular Surveillance lab Sheikh Khalifa Medical City | Amirtharaj Francis et al |
| EPI_ISL_582608,EPI_ISL_582609,EPI_ISL_582611,EPI_ISL_582616 | Sheikh Khalifa Medical City | Molecular/Surveillance lab Sheikh Khalifa Medical City | Amirtharaj Francis et al |
| EPI_ISL_1533610 | Shiraz University | Shiraz University | Abozar Ghorbani et al |
| EPI_ISL_1391211 | SIESP DIPARTIMENTO DI PREVENZIONE TERAMO TERAMO(TERAMO) | Istituto Zooprofilattico Sperimentale dell’Abruzzo e Molise “G. Caporale” | Lorusso A et al |
| EPI_ISL_470592 | Simile | Bioinformatics Laboratory / LNCC | Alexandra Gerber et al |
| EPI_ISL_1749436,EPI_ISL_1969690 | Singapore General Hospital | Department of Microbiology | Nurdyana Abdul Rahman et al |
| EPI_ISL_406973 | Singapore General Hospital | National Public Health Laboratory | Mak et al |
| EPI_ISL_956278,EPI_ISL_1159383 | Siti Khodijah Hospital | Institute of Tropical Disease, Universitas Airlangga | Aldise M Nastri et al |
| EPI_ISL_1629723 | SOMER | Universidad Nacional de Colombia - Laboratorio Genómico One Health | Simón Villegas Velásquez et al |

| | | | | |
|---|---|---|---|---|
| Lobiuc Andrei et al | Sonic Reference Laboratory | Sonic Reference Laboratory | Robert Moller et alEPI_ISL_1941720<br>South Dakota Public Health LaboratoryUniversity of Minnesota Genomics CenterDaryl M. Gohl et alEPI_ISL_1017003<br>South Eastern Area Laboratory ServicesNSW Health Pathology - Institute of Clinical Pathology and Medical Research; Westmead Hospital; University of SydneyCIDM-PH et al.EPI_ISL_455091,EPI_ISL_455099<br>South Eastern Area Laboratory Services (SEALS)NSW Health Pathology - Institute of Clinical Pathology and Medical Research; Westmead Hospital; University of SydneyCIDM-PH et al.EPI_ISL_490021,EPI_ISL_490030,EPI_ISL_526144,EPI_ISL_593712,EPI_ISL_667782,EPI_ISL_768600,EPI_ISL_1904455,EPI_ISL_1904449,EPI_ISL_1904458,EPI_ISL_1904454,EPI_ISL_1904457,EPI_ISL_1904462,EPI_ISL_1904465,EPI_ISL_1904469<br>Southern Community Labs | EPI_ISL_491047,EPI_ISL_491088,EPI_ISL_491090 |
| EPI_ISL_2091024 | Sultanah Aminah Hospital, Johor Bahru | Institute for Medical Research, Infectious Disease Research Centre, National Institutes of Health, Ministry of Health Malaysia | Suppiah J et al | |

| | | | |
|---|---|---|---|
| EPI_ISL_1972356 | Sungai Buloh Hospital | Institute for Medical Research, Infectious Disease Research Centre, National Institutes of Health, Ministry of Health Malaysia | Suppiah J et al |
| EPI_ISL_1544125 | Supratech Micropath Laboratory Research Institute Pvt Ltd, Ahmedabad | Gujarat Biotechnology Research Centre | Sonal Sharma et al |
| EPI_ISL_1629737 | SURA | Universidad Nacional de Colombia - Laboratorio Genómico One Health | Karl A Ciuoderis et al |
| EPI_ISL_475543 | Surbrunns VC | The Public Health Agency of Sweden | Oskar Karlsson Lindsjo et al |
| EPI_ISL_1659751,EPI_ISL_1808654,EPI_ISL_1808658,EPI_ISL_1808686,EPI_ISL_1808683,EPI_ISL_1902079,EPI_ISL_1899279,EPI_ISL_1899297,EPI_ISL_2032727,EPI_ISL_2034589,EPI_ISL_2033405,EPI_ISL_2034582,EPI_ISL_2033751 | Swedish national genomic surveillance program of SARS-CoV-2 | The Public Health Agency of Sweden | Maximilian Riess et al |
| EPI_ISL_1192957 | Swedish national genomic surveillance program of SARS-CoV-2 | The Public Health Agency of Sweden | Swedish national genomic surveillance program of SARS-CoV-2 et al |
| EPI_ISL_1969243 | Swissbel Hotel Airport | National Institute of Health Research and Development | Vivi Setiawaty et al |
| EPI_ISL_526206 | Sydney South West Pathology Service (SSWPS) - Liverpool Hospital - NSW Health Pathology | NSW Health Pathology - Institute of Clinical Pathology and Medical Research; Westmead Hospital; University of Sydney | CIDM-PH et al. |
| EPI_ISL_1919655,EPI_ISL_1970794 | SYNLAB | GIGA Medical Genomics | Keith Durkin et al |

| | | | |
|---|---|---|---|
| EPI_ISL_845627 | SYNLAB COLOMBIA S.A.S | Instituto Nacional de Salud - Dirección de Investigación en Salud Pública | Katherine Laiton-Donato et al |
| EPI_ISL_1190907,EPI_ISL_1319300,EPI_ISL_2107118 | Synlab Eesti OÜ | 1. Laboratory of Communicable Diseases (Estonia); 2. Eurofins Genomics Europe Sequencing GmbH | Liidia Dotsenko et al. |
| EPI_ISL_2095553 | Synlab Haut de France | UMR 8199/1283 EGID | Derhourhi Mehdi et al |
| EPI_ISL_2117300 | SYNLAB Jena Oncoscreen | Robert Koch Institute | ? |
| EPI_ISL_2123440,EPI_ISL_2123616 | SYNLAB Labor MÃ¼nchen Zentrum LMZ | Robert Koch Institute | ? |
| EPI_ISL_1571754,EPI_ISL_2129841 | SYNLAB MVZ Leinfelden-Echterdingen | Robert Koch Institute | ? |
| EPI_ISL_1728331,EPI_ISL_2124794 | SYNLAB MVZ Leverkusen | Robert Koch Institute | ? |
| EPI_ISL_1641831,EPI_ISL_2126465 | SYNLAB MVZ Weiden | Robert Koch Institute | ? |
| EPI_ISL_412968 | Takayuki Hishiki Kanagawa Prefectural Institute of Public Health | Takayuki Hishiki Kanagawa Prefectural Institute of Public Health | Hishiki et al |
| EPI_ISL_2162143 | Taldykorgan Anti-Plague Station | Reference laboratory for the control of viral infections | Nazym Tleumbetova et al |
| EPI_ISL_914796 | TAMIZAJE COMUNITARIO - PASO CANOAS | Inciensa, Instituto Costarricense de Investigación y Enseñanza en Nutrición y Salud | Francisco Duarte et al |
| EPI_ISL_682261 | TAMIZAJE COMUNITARIO- PASO CANOAS | Inciensa, Instituto Costarricense de Investigación y Enseñanza en Nutrición y Salud | Francisco Duarte et al |
| EPI_ISL_1379449 | Tampa General Hospital Esoteric Lab | Tampa General Hospital Esoteric Research & Development Lab | Grant Vestal et al |

| | | | |
|---|---|---|---|
| EPI_ISL_966940 | Technical Support Units for Scientific Research (UATRS), National Centre for Scientific and Technical Research (CNRST) | Technical Support Units for Scientific Research (UATRS), National Centre for Scientific and Technical Research (CNRST) | Touil et al |
| EPI_ISL_733572 | Temporary Specimen Collection Centre | Hong Kong Department of Health | Alan K.L. Tsang et al |
| EPI_ISL_937538,EPI_ISL_1296450 | Thai Red Cross Emerging Infectious Diseases Health Science Centre, Chulalongkorn Hospital, Faculty of Medicine, Chulalongkorn University | Thai Red Cross Emerging Infectious Diseases Center and Faculty of Medicine, Chulalongkorn University | Opass Putcharoen et al |
| EPI_ISL_756311,EPI_ISL_756357,EPI_ISL_1490226 | The Caribbean Public Health Agency | Carrington Lab, Department of PreClinical Sciences, Faculty of Medical Sciences, The University of the West Indies | Nikita S. D. Sahadeo et al |
| EPI_ISL_696519 | Thembalethu CDC wc THC & NHLS/UCT | KRISP, KZN Research Innovation and Sequencing Platform | Arash Iranzadeh et al |
| EPI_ISL_2155984 | The Medical City -Â Ortigas | Philippine Genome Center | Francis A. Tablizo et al |
| EPI_ISL_1838050,EPI_ISL_1838052,EPI_ISL_1838097,EPI_ISL_1838131,EPI_ISL_1838134,EPI_ISL_1838156,EPI_ISL_1838158,EPI_ISL_1838292,EPI_ISL_1838345 | The National Centre for Cell Science | CSIR-Centre for Cellular and Molecular Biology-INSACOG | Dhiraj Paul et al |
| EPI_ISL_577626,EPI_ISL_577630,EPI_ISL_584076,EPI_ISL_584081,EPI_ISL_626589,EPI_ISL_850664 | The National Institute of Public Health | State Veterinary Institute Prague | Nagy et al |
| EPI_ISL_827291,EPI_ISL_830235,EPI_ISL_829948 | The National University Hospital of Iceland | deCODE genetics | Daniel F Gudbjartsson et al |

| | | | |
|---|---|---|---|
| EPI_ISL_891225 | The Oncology Institute "Prof. Dr. Ion Chiricuta" Cluj Napoca | Stefan cel Mare University Metagenomics Lab | Lobiuc Andrei et al |
| EPI_ISL_754233,EPI_ISL_754235 | The Republican Research and Practical Center for Epidemiology and Microbiology (RRPCEM) | WHO National Influenza Centre Russian Federation | Elena Gasich et al |
| EPI_ISL_1196006 | TLC Clinic | National Institute for Communicable Diseases of the National Health Laboratory Service | Maphalala GP et al |
| EPI_ISL_480038 | Tochigi Prefectural Institute of Public Health and Environmental Science | Pathogen Genomics Center, National Institute of Infectious Diseases | Tsuyoshi Sekizuka et al |
| EPI_ISL_538254,EPI_ISL_560358,EPI_ISL_1225259 | TriCore Reference Laboratories | Center for Global Health, University of New Mexico Health Sciences Center | Daryl Domman et al |
| EPI_ISL_717692,EPI_ISL_717695,EPI_ISL_717694,EPI_ISL_717700,EPI_ISL_717697,EPI_ISL_756363 | Trinidad Public Health Laboratory | Carrington Lab, Department of PreClinical Sciences, Faculty of Medical Sciences, The University of the West Indies | Nikita S. D. Sahadeo et al |
| EPI_ISL_436099,EPI_ISL_436108,EPI_ISL_447252 | TSGH-CP molecular lab | TSGH-CP molecular lab | Cherng-Lih Perng et al |
| EPI_ISL_437336 | TSGH-CP molecular lab, Division of Clinical Pathology, Department of Pathology | TSGH-CP molecular lab, Division of Clinical Pathology, Department of Pathology | Cherng-Lih Perng et al |
| EPI_ISL_1960202 | UAB Medicina practica laboratorija | National Public Health Surveillance Laboratory | Lukas Zemaitis et al |
| EPI_ISL_737956,EPI_ISL_737942,EPI_ISL_737965,EPI_ISL_737969,EPI_ISL_737976,EPI_ISL_738008,EPI_ISL_738023,EPI_ISL_738028,EPI_ISL_738038 | Uganda Central Public Health Lab and Uganda Virus Research Institute | MRC/UVRI & LSHTM Uganda Research Unit | Matthew Cotten et al |

| | | | |
|---|---|---|---|
| EPI_ISL_451188,EPI_ISL_451199,EPI_ISL_451201 | Uganda Virus Research Institute | MRC/UVRI & LSHTM Uganda Research Unit | Dan Lule Bugembe et al |
| EPI_ISL_628761,EPI_ISL_648124,EPI_ISL_648125,EPI_ISL_861460,EPI_ISL_861459 | UHAS COVID-19 Lab | UHAS COVID-19 Lab | Kwabena O. Duedu et al |
| EPI_ISL_1745236 | ULSS 7 Pedemontana - Distretto 2 | Istituto Zooprofilattico Sperimentale delle Venezie | Adelaide Milani et al |
| EPI_ISL_1993936,EPI_ISL_2099526,EPI_ISL_2132653 | UMC Groningen, Clinical Virology, Department of Medical Microbiology and Infection Prevention | UMC Groningen, Clinical Virology, Department of Medical Microbiology and Infection Prevention | Hubert Niesters et al |
| EPI_ISL_1287766 | Unidad de Investigación Biomédica de Zacatecas (UIBZ) | Instituto Nacional de Enfermedades Respiratorias (INER): Centro de Investigación en Enfermedades Infecciosas (CIENI) | Consorcio Mexicano de Vigilancia Genómica (CoViGen-Mex). Authors (in alphabetical order): Julio Elias Alvarado-Yaah et al |
| EPI_ISL_2091233 | Unidad de Investigación Médica de Yucatán (UIMY) | Centro de Investigación en Enfermedades Infecciosas (CIENI), Instituto Nacional de Enfermedades Respiratorias (INER) | Consorcio Mexicano de Vigilancia Genómica (CoViGen-Mex). Authors (in alphabetical order): Julio Elias Alvarado-Yaah et al |
| EPI_ISL_1585433,EPI_ISL_1585434 | Unidad de Investigación Médica de Yucatán (UIMY) | Instituto Nacional de Enfermedades Respiratorias (INER): Centro de Investigación en Enfermedades Infecciosas (CIENI) | Consorcio Mexicano de Vigilancia Genómica (CoViGen-Mex). Authors (in alphabetical order): Julio Elias Alvarado-Yaah et al |
| EPI_ISL_2101685 | Unidade de apoio ao diagnóstico da COVID – UNADIG | Bioinformatics Laboratory / LNCC | Luiz G P de Almeida et al |
| EPI_ISL_693226 | Unidade de Pronto Atendimento Sao José | Instituto Adolfo Lutz, Interdiciplinary Procedures Center, Strategic Laboratory | Claudio Tavares Sacchi et al |

| | | | |
|---|---|---|---|
| EPI_ISL_794656 | UNIDAD HEMATOLOGICA ESPECIALIZADA | Instituto Nacional de Salud - Dirección de Investigación en Salud Pública | Katherine Laiton-Donato et al |
| EPI_ISL_590917 | Unilabs Laboratory Medicine | Norwegian Institute of Public Health, Department of Virology | Kathrine Stene-Johansen et al |
| EPI_ISL_812563,EPI_ISL_812569,EPI_ISL_812575 | United States Air Force School of Aerospace Medicine | United States Air Force School of Aerospace Medicine | Anthony Fries et al |
| EPI_ISL_411951 | Unit for Laboratory Development and Technology Transfer, Public Health Agency of Sweden | Unit for Laboratory Development and Technology Transfer, Public Health Agency of Sweden | Bengner et al |
| EPI_ISL_1299863 | Unit of lab surveillance of viral emerging diseases, National Lab of Influenza | Respiratory Virus Unit, National Infection Service, Public Health England | PHE Covid Sequencing Team et al |
| EPI_ISL_526268 | Unity Health Toronto | Ontario Institute for Cancer Research | Ramzi Fattouh et al |
| EPI_ISL_2157423 | UNIVERSIDADE FEDERAL DE VIÇOSA | Laboratory of Respiratory Viruses and Measles, Oswaldo Cruz Institute, FIOCRUZ | Paola Resende et al |
| EPI_ISL_523812 | Universidad Iberoamericana, Instituto de Medicina Tropical & Salud Global | International Centre for Genetic Engineering and Biotechnology (ICGEB) and ARGO Open Lab Platform | Robert Paulino-Ramirez et al |
| EPI_ISL_1692748 | Università Federico II - Dipartimento di scienze mediche traslazionali - Napoli | Telethon Institute of Genetics and Medicine (TIGEM) | Antonio Grimaldi Patrizia Annunziata Francesco Panariello Teresa Giuliano Michele Cennamo Valentina Bouche Chiara Colantuono Lucio Di Filippo Mariano Fiorenza Anna Manfredi Marcello Salvi Giuseppe Portella Andrea Ballabio Davide Cacchiarelli et al |

| | | | |
|---|---|---|---|
| EPI_ISL_1939228 | UniversitätsSpital Zürich | Institute of Medical Virology | Verena Kufner et al |
| EPI_ISL_812967 | University Clinical Research Center, University of Sciences | University Clinical Research Center, University of Sciences | Diarra et al |
| EPI_ISL_581980 | University Hospital Basel, Clinical Virology | University Hospital Basel, Clinical Bacteriology | Madlen Stange et al |
| EPI_ISL_931063 | University Hospital Basel, Clinical Virology | University Hospital Basel, Clinical Bacteriology | Tim Roloff et al |
| EPI_ISL_631388 | University Hospital Cologne | Center of Medical Microbiology, Virology, and Hospital Hygiene, University of Duesseldorf | Maximilian Damagnez et al |
| EPI_ISL_710565,EPI_ISL_710571 | University Hospital Dubrava | Ruđer Boškovic Institute; Forensic Science Centre Ivan Vučetić; University of Zagreb Faculty of Science | Robert Belužić et al |
| EPI_ISL_415457,EPI_ISL_415699 | University Hospitals of Geneva Laboratory of Virology | University Hospitals of Geneva Laboratory of Virology | Laubscher F. et al |
| EPI_ISL_1296150,EPI_ISL_1369531,EPI_ISL_1811202,EPI_ISL_1963298,EPI_ISL_1963299,EPI_ISL_1963300 | University Hospitals of Geneva, Laboratory of Virology | HUG, Laboratory of Virology and the Health2030 Genome Center | Samuel Cordey et al |
| EPI_ISL_776042,EPI_ISL_959973 | University Medical Center Hamburg Eppendorf | Heinrich Pette Institute, Leibniz Institute for Experimental Virology | Alexis Robitaille et al |
| EPI_ISL_1072985 | University of Balamand | Microbial Genomics Lab, Lebanese American University, Byblos | Mira El Chaar et al |
| EPI_ISL_1922125 | University of Bari Biomedical Sciences and Human Oncology | University of Bari Biomedical Sciences and Human Oncology | Chironna M. et al |

| | | | |
|---|---|---|---|
| EPI_ISL_1133254 | University of Chittagong | Central Biological Research Laboratory and Department of Biochemistry and Molecular Biology | H. M. Abdullah Al Masud et al |
| EPI_ISL_671423,EPI_ISL_671453 | University of Debrecen, Department of Medical Microbiology | National Laboratory of Virology, Szentágothai Research Centre | Endre Gábor Tóth et al |
| EPI_ISL_2100508 | University of Health Sciences | Quadram Institute Bioscience | Muhammad Bilal Sarwar et al |
| EPI_ISL_1016970 | University of Sarajevo, Veterinary Faculty, Laboratory for Molecular Diagnostic and Research Laboratory | University of Sarajevo, Veterinary Faculty, Laboratory for Molecular Diagnostic and Research Laboratory | Goletic S. et al |
| EPI_ISL_955155,EPI_ISL_955144,EPI_ISL_955142,EPI_ISL_955148,EPI_ISL_955146 | University of Sarajevo, Veterinary Faculty, Laboratory for Molecular Diagnostic and Research Laboratory | University of Sarajevo, Veterinary Faculty, Laboratory for Molecular Diagnostic and Research Laboratory | Goletić T. et al |
| EPI_ISL_677748,EPI_ISL_677759 | University of Szeged, Institute of Clinical Microbiology | National Laboratory of Virology, Szentágothai Research Centre | Endre Gábor Tóth et al |
| EPI_ISL_484835,EPI_ISL_509829,EPI_ISL_509953,EPI_ISL_541544,EPI_ISL_788901 | University of Wisconsin-Madison AIDS Vaccine Research Laboratories | University of Wisconsin-Madison AIDS Vaccine Research Laboratories | Gage Moreno et al |
| EPI_ISL_977256,EPI_ISL_977272,EPI_ISL_977325,EPI_ISL_977287,EPI_ISL_977421,EPI_ISL_977336,EPI_ISL_977339,EPI_ISL_977355 | University of Zambia, School of Veterinary Medicine | UNZAVET and PATH | Mulenga Mwenda-Chimfwembe et al |
| EPI_ISL_463001 | unknown | Clinical virology | Fares et al |
| EPI_ISL_468160 | unknown | Department of Virology, Public Health Laboratories Division, National Institute of Health | Massab Umair et al |

| | | | |
|---|---|---|---|
| EPI_ISL_454216,EPI_ISL_1739160,EPI_ISL_1739170,EPI_ISL_1739180,EPI_ISL_1739182 | unknown | Instituto Nacional de Saude (INSA) | Borges et al |
| EPI_ISL_476559 | unknown | Laboratoire Sciences et Technologies de la Santé (STS) Institut Supérieur des Sciences de la Santé Université Hassan 1er, Settat, Morocco | Hajar Lemriss et al |
| EPI_ISL_483060 | unknown | Microbiology, Canterbury Health Laboratories | Dilcher et al |
| EPI_ISL_1938308 | unknown | PHV-FSS | Son Nguyen et al |
| EPI_ISL_1373715,EPI_ISL_1732346 | UPMC Clinical Microbiology Laboratory | Microbial Genome Sequencing Center; Microbial Genomic Epidemiology Laboratory, University of Pittsburgh | Lee H. Harrison et al |
| EPI_ISL_708817 | Urban Institute for Disease Prevention and Control | National Institute of Health, Department of Medical Sciences, Ministry of Public Health, Thailand | Pilailuk Okada et al |
| EPI_ISL_524361,EPI_ISL_930722,EPI_ISL_1300601,EPI_ISL_1300544,EPI_ISL_1403864 | Utah Public Health Laboratory | Utah Public Health Laboratory | Erin L. Young et al |
| EPI_ISL_746996 | Utah Public Health Laboratory | Utah Public Health Laboratory | Erin Young et al |
| EPI_ISL_570461,EPI_ISL_570515,EPI_ISL_570757,EPI_ISL_570954,EPI_ISL_1324138,EPI_ISL_1324141,EPI_ISL_1498061,EPI_ISL_1616618,EPI_ISL_1789413,EPI_ISL_2082524,EPI_ISL_2082694,EPI_ISL_2104866 | UW Virology Lab | UW Virology Lab | Pavitra Roychoudhury et al |

| | | | |
|---|---|---|---|
| EPI_ISL_734583 | UZ Leuven, National Reference Laboratory for Coronaviruses, Laboratory Medicine, Leuven, Belgium | KU Leuven, Rega Institute, Clinical and Epidemiological Virology | Tony Wawina-Bokalanga et al |
| EPI_ISL_1168406,EPI_ISL_1168421 | VA Connecticut Healthcare System | Grubaugh Lab - Yale School of Public Health | Mary Petrone et al |
| EPI_ISL_1373885,EPI_ISL_1373949,EPI_ISL_1373983 | Vanderbilt University Medical Center | Pathogen Discovery, Respiratory Viruses Branch, Division of Viral Diseases, Centers for Disease Control and Prevention | Jing Zhang et al |
| EPI_ISL_452236 | VC Sorgenfrimottagningen | The Public Health Agency of Sweden | Lisa Esbjornsson Klemendz et al |
| EPI_ISL_1662284,EPI_ISL_1663498 | Veer Surendra Sai Institute of Medical Sciences and Research, Burla, Sambalpur | Institute of Life Sciences - INSACOG | Sunil K. Raghav et al |
| EPI_ISL_1118534 | Vestfold Hospital, Toensberg Department of Microbiology | Norwegian Institute of Public Health, Department of Virology | Kathrine Stene-Johansen et al |
| EPI_ISL_904010 | Veterinary Specialized Institute Kraljevo | Veterinary Specialized Institute "Kraljevo", Serbia | Vidanovic et al |
| EPI_ISL_582532 | Veterinary Specialized Institute "Kraljevo", Serbia | Veterinary Specialized Institute "Kraljevo", Serbia | Vidanovic et al |
| EPI_ISL_833511 | Veterinary Specialized Instute "Nis" | Veterinary Specialized Institute "Kraljevo", Serbia | Vidanovic et al |
| EPI_ISL_833573 | Veterinary Specialized Instute "Nis" | Veterinary Specialized Institute "Sabac", Serbia | Vidanovic et al |
| EPI_ISL_640141,EPI_ISL_1534295,EPI_ISL_2140692 | Victoria Hospital wc VHW | NHLS/UCT | Arash Iranzadeh et al |
| EPI_ISL_426673,EPI_ISL_436117,EPI_ISL_456488,EPI_ISL_456587 | Victorian Infectious Diseases Reference Laboratory (VIDRL) | Microbiological Diagnostic Unit Public Health Laboratory and Victorian Infectious Diseases Reference Laboratory, Doherty Institute | Caly L. et al |

| | | | |
|---|---|---|---|
| EPI_ISL_430709,EPI_ISL_430524 | Victorian Infectious Diseases Reference Laboratory (VIDRL) | Microbiological Diagnostic Unit Public Health Laboratory and Victorian Infectious Diseases Reference Laboratory, The Peter Doherty Institute for Infection and Immunity | Caly L. et al |
| EPI_ISL_416411,EPI_ISL_416410,EPI_ISL_419734 | Victorian Infectious Diseases Reference Laboratory (VIDRL) | Victorian Infectious Diseases Reference Laboratory and Microbiological Diagnostic Unit Public Health Laboratory, Doherty Institute | Caly L. et al |
| EPI_ISL_480650,EPI_ISL_480651,EPI_ISL_480702,EPI_ISL_480742,EPI_ISL_521277,EPI_ISL_521880,EPI_ISL_522085,EPI_ISL_522098,EPI_ISL_593411,EPI_ISL_593412,EPI_ISL_640337,EPI_ISL_1033154,EPI_ISL_1249993,EPI_ISL_1913202,EPI_ISL_1913212 | Victorian Infectious Diseases Reference Laboratory (VIDRL) | VIDRL and MDU-PHL | Caly L. et al |
| EPI_ISL_1005421 | Vilnius university hospital Santaros Klinikos, Center of Laboratory Medicine | Vilnius university hospital Santaros Klinikos, Center of Laboratory Medicine | Gytis Dudas et al |
| EPI_ISL_802860,EPI_ISL_904939 | Vilnius University Hospital Santaros Klinikos, Vilnius University | Institute of Biotechnology, Life Sciences Center, Vilnius University | Emilija Vasiliunaite et al |
| EPI_ISL_560394,EPI_ISL_560397,EPI_ISL_560395 | Vilnius University Hospital Santaros Klinikos, Vilnius University | Institute of Biotechnology, Life Sciences Center, Vilnius University and Thermo Fisher Scientific | Justinas Slikas et al |

| | | | |
|---|---|---|---|
| EPI_ISL_1914236,EPI_ISL_1914238,EPI_ISL_1914239 | Viollier AG | Department of Biosystems Science and Engineering, ETH Zurich | Chaoran Chen et al |
| EPI_ISL_1658295 | Viollier AG | Department of Biosystems Science and Engineering, ETH Zurich | Christian Beisel et al |
| EPI_ISL_728931,EPI_ISL_796566,EPI_ISL_1001690,EPI_ISL_1119544,EPI_ISL_1195176,EPI_ISL_1361039,EPI_ISL_1750651,EPI_ISL_1750690,EPI_ISL_2018995,EPI_ISL_2019338 | Viollier AG | Department of Biosystems Science and Engineering, ETH Zürich | Chaoran Chen et al |
| EPI_ISL_483656,EPI_ISL_486495,EPI_ISL_539469,EPI_ISL_560449,EPI_ISL_693898,EPI_ISL_1003119,EPI_ISL_1259729,EPI_ISL_1259649,EPI_ISL_1598117,EPI_ISL_2017208,EPI_ISL_2017225 | Viollier AG | Department of Biosystems Science and Engineering, ETH Zürich | Christian Beisel et al |
| EPI_ISL_2017815 | Viral Respiratory Infections Laboratory, Cantacuzino National Military-Medical Institute | Cantacuzino Institute Virology | Luiza Ustea et al |
| EPI_ISL_2100242 | Viral Respiratory Infections Laboratory, Cantacuzino National Military-Medical Institute | Cantacuzino Institute Virology | Sorin Dinu et al |

| | | | |
|---|---|---|---|
| EPI_ISL_437348,EPI_ISL_471400,EPI_ISL_471403,EPI_ISL_527527,EPI_ISL_591083,EPI_ISL_591084,EPI_ISL_591085,EPI_ISL_591086,EPI_ISL_965124,EPI_ISL_1623756,EPI_ISL_1623757,EPI_ISL_1972308,EPI_ISL_1972323,EPI_ISL_2135841,EPI_ISL_2135846,EPI_ISL_2135844 | Viral Respiratory Lab, National Institute for Biomedical Research (INRB) | Pathogen Sequencing Lab, National Institute for Biomedical Research (INRB) | Placide Mbala-Kingebeni et al |
| EPI_ISL_429972,EPI_ISL_485861,EPI_ISL_572280,EPI_ISL_614052 | Virginia DCLS | Virginia DCLS | Virginia DCLS et al |
| EPI_ISL_2090606 | VIROLOGY, AFRIMS | VIROLOGY, AFRIMS | Velasco et al |
| EPI_ISL_1191597,EPI_ISL_1191602,EPI_ISL_1191603,EPI_ISL_1191604,EPI_ISL_1827670 | Virology Department, Victoria Hospital, Plaine-Wilhems, Mauritius | National Institute for Communicable Diseases of the National Health Laboratory Service | Ramuth M et al |
| EPI_ISL_890237 | Virology, International Centre for Diarrhoeal Disease Research, Bangladesh (ICDDR,B) | International Centre for Diarrhoeal Disease Research (ICDDR,B) | Hossain et al |
| EPI_ISL_493203 | Virology Lab,Department of Pathology, National Cheng Kung University Hospital | Virology Lab,Department of Pathology, National Cheng Kung University Hospital | Huey-Pin Tsai et al |
| EPI_ISL_416458 | Virology laboratory Ministry of Health Kuwait sequenced at Dasman Diabetes Institute | Dasman Diabetes Institute | Fahd Al-Mulla et al |
| EPI_ISL_1970570 | Virology Laboratory, Scientific Department, Army Medical Center | Virology Laboratory, Scientific Department, Army Medical Center | Silvia Fillo et al |

| | | | |
|---|---|---|---|
| EPI_ISL_508863,EPI_ISL_625456,EPI_ISL_677636,EPI_ISL_1660225,EPI_ISL_1660234,EPI_ISL_1660238,EPI_ISL_1660241,EPI_ISL_1660282,EPI_ISL_1660285,EPI_ISL_1660291,EPI_ISL_1660288,EPI_ISL_1660289,EPI_ISL_1660290,EPI_ISL_1672390 | Virology Unit, Institut Pasteur de Madagascar | Virology Unit, Institut Pasteur de Madagascar | Christian Ranaivoson et al |
| EPI_ISL_1660310,EPI_ISL_1660323,EPI_ISL_1660324,EPI_ISL_1660328,EPI_ISL_1660331 | Virology Unit, Institut Pasteur de Madagascar | Virology Unit, Institut Pasteur de Madagascar | Marion Barbet et al |
| EPI_ISL_1711996,EPI_ISL_1818965,EPI_ISL_2106254,EPI_ISL_2106276 | Virology Unit, Institut Pasteur du Cambodge | Virology Unit, Institut Pasteur du Cambodge | Jurre Y Siegers et al |
| EPI_ISL_918363,EPI_ISL_918364,EPI_ISL_918367,EPI_ISL_918369,EPI_ISL_933783,EPI_ISL_933785,EPI_ISL_1532809 | Virology Unit, Institut Pasteur du Cambodge | Virology Unit, Institut Pasteur du Cambodge | Sokhoun Yann et al |
| EPI_ISL_411902 | Virology Unit, Institut Pasteur du Cambodge. | Virology Unit, Institut Pasteur du Cambodge (Sequencing done by: Jessica E Manning/Jennifer A Bohl at Malaria and Vector Research Research Laboratory, National Institute of Allergy and Infectious Diseases and Vida Ahyong from Chan-Zuckerberg Biohub) | Erik A Karlsson et al |
| EPI_ISL_452138 | VI-US Virgin Islands Department of Health | Pathogen Discovery, Respiratory Viruses Branch, Division of Viral Diseases, Centers for Disease Control and Prevention | Anna Uehara et al |

| | | | |
|---|---|---|---|
| EPI_ISL_454647 | VI-US Virgin Islands Department of Health | Pathogen Discovery, Respiratory Viruses Branch, Division of Viral Diseases, Centers for Disease Control and Prevention | Ying Tao et al |
| EPI_ISL_2017734 | VRDL, Goverment Medical College (GMC), Surat | Gujarat Biotechnology Research Centre | Janvi Raval et al |
| EPI_ISL_903650,EPI_ISL_903694 | VT Dept. of Health Laboratory | Genomics and Discovery, Respiratory Viruses Branch, Division of Viral Diseases, Centers for Disease Control and Prevention | Krista Queen et al |
| EPI_ISL_676928,EPI_ISL_676721,EPI_ISL_754724 | Wadsworth Center, New York State Department.of Health | Wadsworth Center, New York State Department.of Health | Kirsten St. George et al |
| EPI_ISL_456168,EPI_ISL_456259 | Waikato Hospital | Institute of Environmental Science and Research (ESR) | Matt Storey et al |
| EPI_ISL_707794 | Waikato Hospital | Institute of Environmental Science and Research (ESR) | Xiaoyun Ren et al |
| EPI_ISL_434069 | Washington State Department of Health | Seattle Flu Study | Chu et al |
| EPI_ISL_417143 | Washington State Department of Health | Seattle Flu Study | Chu etl al et al |
| EPI_ISL_497484 | Washington State Department of Health | Seattle Flu Study | Deborah A. Nickerson et al |
| EPI_ISL_456182,EPI_ISL_456187,EPI_ISL_456188,EPI_ISL_456319,EPI_ISL_456332,EPI_ISL_456400 | Wellington SCL | Institute of Environmental Science and Research (ESR) | Matt Storey et al |
| EPI_ISL_1250695 | Wellington SCL (WN) | Institute of Environmental Science and Research (ESR) | Rachel Boyle et al |
| EPI_ISL_637085 | Wellington SCL (WN) | Institute of Environmental Science and Research (ESR) | Xiaoyun Ren et al |

| | | | |
|---|---|---|---|
| EPI_ISL_1255113,EPI_ISL_1255154,EPI_ISL_1255199,EPI_ISL_1255250,EPI_ISL_1255256,EPI_ISL_1255275 | West African Centre for Cell Biology of Infectious Pathogens (WACCBIP), University of Ghana, Accra, Ghana | West African Centre for Cell Biology of Infectious Pathogens (WACCBIP), University of Ghana, Volta Road, Legon-Accra, Ghana | Collins M. Morang'a et al |
| EPI_ISL_1255105 | West African Centre for Cell Biology of Infectious Pathogens (WACCBIP), University of Ghana, Volta Road, Legon-Accra, Ghana | West African Centre for Cell Biology of Infectious Pathogens (WACCBIP), University of Ghana, Volta Road, Legon-Accra, Ghana | Collins M. Morang'a et al |
| EPI_ISL_1740514 | WESTCHESTER MEDICAL CENTER | Wadsworth Center, New York State Department of Health | Kirsten St. George et al |
| EPI_ISL_450284,EPI_ISL_524017,EPI_ISL_524048,EPI_ISL_733264,EPI_ISL_1789541,EPI_ISL_1789542,EPI_ISL_1797437 | WHO National Influenza Centre Russian Federation | WHO National Influenza Centre Russian Federation | Andrey Komissarov et al |
| EPI_ISL_578688,EPI_ISL_1794346,EPI_ISL_1794352 | Wisconsin State Laboratory of Hygiene Communicable Disease Division | Wisconsin State Laboratory of Hygiene Communicable Disease Division | Kelsey R. Florek et al |
| EPI_ISL_708812 | World Medical Hospital | National Institute of Health, Department of Medical Sciences, Ministry of Public Health, Thailand | Pilailuk Okada et al |
| EPI_ISL_1909094,EPI_ISL_1909095,EPI_ISL_1909092 | WSSE Katowice | 1. Academic Center for Pathomorphological and Genetic-Molecular Diagnostics ltd, Bialystok, Poland 2. National Institute of Public Health - National Institute of Hygiene, Warsaw, Poland | Radosław Charkiewicz et al |

| | | | |
|---|---|---|---|
| EPI_ISL_2090620 | WSSE w Warszawie | National Institute of Public Health - National Institute of Hygiene | Wołkowicz Tomasz et al |
| EPI_ISL_454986 | Wuhan Chain Medical Labs (CMLabs) | State Key Laboratory of Biotherapy of Sichuan University | Baowen Du et al |
| EPI_ISL_1446719 | WVDHHR - Office of Laboratory Services | Centers for Disease Control and Prevention Division of Viral Diseases, Pathogen Discovery | Mili Sheth et al |
| EPI_ISL_540439 | WVDHHR - Office of Laboratory Services | Pathogen Discovery, Respiratory Viruses Branch, Division of Viral Diseases, Centers for Disease Control and Prevention | Ying Tao et al |
| EPI_ISL_462959,EPI_ISL_614248,EPI_ISL_614254,EPI_ISL_614264,EPI_ISL_1627062 | Wyoming Public Health Laboratory | Center for Global Health, University of New Mexico Health Sciences Center | Daryl Domman et al |
| EPI_ISL_1140085,EPI_ISL_1468663,EPI_ISL_1571784,EPI_ISL_1593776 | Wyoming Public Health Laboratory | Wyoming Public Health Laboratory | Jim Mildenberger et al |
| EPI_ISL_1272962 | WY Public Health Laboratory | Centers for Disease Control and Prevention Division of Viral Diseases, Pathogen Discovery | Krista Queen et al |
| EPI_ISL_1225681 | Yale Clinical Virology Lab | Grubaugh Lab - Yale School of Public Health | Joseph Fauver et al |
| EPI_ISL_684824 | Yamagata Prefectural Institute of Public Health | Pathogen Genomics Center, National Institute of Infectious Diseases | Tsuyoshi Sekizuka et al |
| EPI_ISL_1911195,EPI_ISL_1911250,EPI_ISL_1911196,EPI_ISL_1911197 | Zhejiang Provincial Center for Disease Control and Prevention Zhoushan Center for Disease Prevention and Contol | Zhejiang Province Center of Disease Control and prevention | Yanjun Zhang et al |

| | | | |
|---|---|---|---|
| EPI_ISL_660070 | Zurita & Zurita Laboratorios | Zurita & Zurita Laboratorios | Gabriela Sevillano Camilo Zurita-Salinas Karen Loaiza David Ortega-Paredes Jeannete Zurita et al |
